# A theory of brain-computer interface learning via low-dimensional control

**DOI:** 10.1101/2024.04.18.589952

**Authors:** J. A. Menéndez, J. A. Hennig, M. D. Golub, E. R. Oby, P. T. Sadtler, A. P. Batista, S. M. Chase, B. M. Yu, P. E. Latham

## Abstract

A remarkable demonstration of the flexibility of mammalian motor systems is primates’ ability to learn to control brain-computer interfaces (BCIs). This constitutes a completely novel motor behavior, yet primates are capable of learning to control BCIs under a wide range of conditions. BCIs with carefully calibrated decoders, for example, can be learned with only minutes to hours of practice. With a few weeks of practice, even BCIs with randomly constructed decoders can be learned. What are the biological substrates of this learning process? Here, we develop a theory based on a re-aiming strategy, whereby learning operates within a low-dimensional subspace of task-relevant inputs driving the local population of recorded neurons. Through comprehensive numerical and formal analysis, we demonstrate that this theory can provide a unifying explanation for disparate phenomena previously reported in three different BCI learning tasks, and we derive a novel experimental prediction that we verify with previously published data. By explicitly modeling the underlying neural circuitry, the theory reveals an interpretation of these phenomena in terms of biological constraints on neural activity.

## Introduction

A core property of mammalian motor systems is their capacity to adapt to novel environments. Through learning, mammals are able to tailor their movements to an astonishing variety of previously unexperienced tasks, often needing only minutes to hours of practice to do so.^1–7^ A particularly remarkable demonstration of this is offered by brain-computer interfaces (BCIs), where the movement of a cursor on a screen is determined by cortical activity via an external decoder.^8–10^ Despite the unfamiliarity of this motor task, human and non-human primates are capable of learning to control BCIs under a wide range of conditions, often with little practice. With a carefully calibrated BCI decoder, proficient control of the BCI cursor can be learned after only minutes of experience.^11–14^ But even effectively random BCI decoders can be learned as well, provided the subject undergoes a more extensive training procedure (e.g. a few weeks).^15,16^ The purpose of this study is to develop a theory of the algorithm(s) underlying this learning process.

Previous models of motor cortical BCI learning have postulated that synaptic plasticity within motor cortex underlies learning a BCI.^17–20^ Indeed, models of the synaptic connectivity required for a recurrent network to solve a BCI reaching task^19^ and the plasticity rules by which that connectivity might be learned^20^ can account for slow and fast learning of different BCI decoders. However, a fundamental limitation of synaptic plasticity is the curse of dimensionality: motor cortex contains trillions of synapses, so learning via optimization of their weights would entail solving an extremely high-dimensional optimization problem. In the best of cases – when the objective function and its gradient are explicitly known – solving such problems typically requires vast amounts of training data. In the case of BCI learning, the subject’s motor system has no explicit access to the BCI decoder, so the relationship between internal neural activity and movement – and, by extension, task performance – is unknown. This means that gradients of task performance with respect to internal biological parameters must be estimated through trial and error,^20,21^ which is notoriously slow in high dimensional spaces.^22,23^ Moreover, this estimation problem is made even more difficult by the biological constraints of neurons and synapses, which impose noise in the learning signals available to each synapse^24^ and preclude synaptic plasticity rules from back-propagating gradients through the many layers of neural circuitry.^25–27^ These considerations suggest that BCI learning by synaptic plasticity in motor cortex should be slow and highly limited.

Such slow and limited learning is inconsistent with the strikingly fast and flexible learning observed in many BCI experiments, where non-human primates are observed to achieve proficient control after only a single session of 10’s to 100’s of trials of practice.^11–14,28^ Moreover, the hypothesis that motor cortex undergoes substantial synaptic changes over learning is inconsistent with two additional observations. First, the statistical structure of motor cortical activity remains remarkably conserved after learning: the repertoire of activity patterns employed for BCI control is unchanged after training on a new decoder for a few hours,^18,29,30^ and single neuron tuning to manual reaches also remains largely unchanged after performing a BCI reaching task.^31^ Second, learning a BCI task can occur without interfering with natural limb control^31^ (but see^32^).

Together, these observations suggest that synaptic plasticity in motor cortex is not the primary mechanism underlying BCI learning, at least for the short timescales of learning observed in the studies cited above. Instead, they suggest that the brain might take a more parsimonious learning strategy, in which (1) learning is reduced to a low-dimensional optimization problem to enable data-efficient learning, and (2) the motor cortical machinery for natural movements is kept intact.

A learning strategy that satisfies these two criteria is that of “re-aiming”^11,12,33^ or “intrinsic variable learning”.^34,35^ Under this strategy, the animal exploits the pre-existing motor cortical circuitry by learning an association between intended BCI movements and internal motor commands that would otherwise be used during natural motor behavior. For example, if the BCI decoder were such that motor cortical activity generated during a *leftward* arm reach would lead to an *upward* BCI movement, then the animal would learn to employ the motor command usually reserved for *leftward* arm reaches to achieve this *upward* BCI movement (fig. 1a). This strategy satisfies criteria 1 and 2 above: the dimensionality of the learning problem is kept low because both BCI movements – typically movements of a 2D or 3D cursor – and natural motor commands^36–41^ are low-dimensional, and the motor cortical circuit can be kept intact because the patterns of activity used for manual and BCI control are the same.

**Figure 1:**
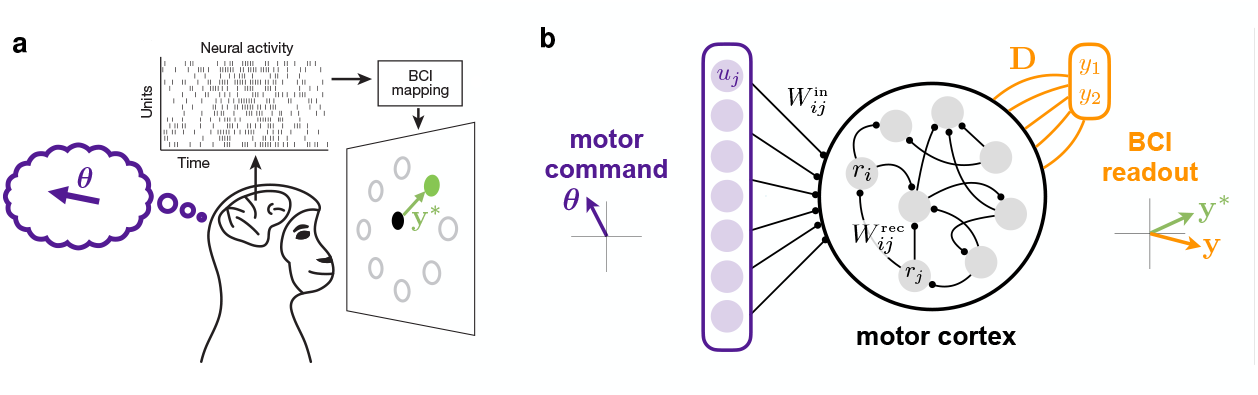
The re-aiming learning strategy. a. Re-aiming strategy for BCI learning. If activity evoked by imagining a leftward planar movement moves the BCI cursor right, then the animal learns to use this motor command to move the cursor to the right. Critically, the space of imagined planar movements is low-dimensional. b. Proposed model of re-aiming. Upstream inputs to motor cortex, *u*_*j*_, depend on a low-dimensional motor command vector, ***θ*** (depicted here as two-dimensional). The BCI readout, **y**, is a 2D linear readout of motor cortical firing rates through a decoding matrix, **D**. Re-aiming is formalized as identifying the motor command, ***θ***, that ensures the BCI readout gets as close as possible to a given target readout, **y**^*^ (cf. equation 4).

Previous experimental results have suggested that re-aiming can account for some^34,35^ but not all^11,12,33^ of the changes in motor cortical activity that occur after learning a novel BCI decoder. However, this evidence has typically been interpreted through the lens of a feed-forward spatial tuning curve model of motor cortex, which does not take into account the influence of additional motor variables beyond reach direction, and omits biological constraints on the dynamics of cortical circuits. Here, we address these limitations by modeling motor cortex as a non-linear recurrently connected network of neurons and modeling re-aiming as an optimization over low-dimensional motor commands driving this network. Via simulation and analysis, we derive predictions of this theory about how neural activity and behavior should change under a pure re-aiming learning strategy, for three distinct BCI learning tasks. These predictions reveal a potentially unifying explanation of disparate phenomena observed in BCI learning.

## Results

### 2.1 Re-aiming as optimization of low-dimensional inputs to motor cortex

We begin by modeling motor cortex as a recurrent neural network driven by an upstream population of neurons (fig. 1b),

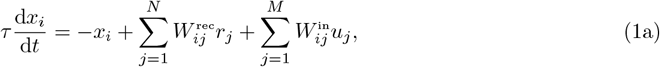

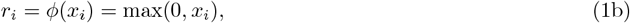

where *r*_1_, *r*_2_, …, *r*_*N*_ and *u*_1_, *u*_2_, …, *u*_*M*_ denote the firing rates of the motor cortical and upstream neurons, respectively. A rectified linear activation function *ϕ*(·) is used to ensure that firing rates are strictly non-negative. We assume that firing rates are low at the start of each trial of BCI control, and thus set the initial conditions to 0, *x*_*i*_(*t* = 0) = 0. The weights 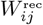 and 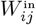 represent the strengths of the synaptic connections between neurons within motor cortex and from the upstream population to motor cortex, respectively. To avoid making any strong commitments about the structure of these connections, we use randomly connected networks throughout the main text; simulations with other, more realistic, connectivity patterns yield similar results (see Supplementary Figure S1).

Next, we consider the upstream inputs to motor cortex, {*u*_*i*_(*t*)}. Inspired by recent models and theories of motor cortex,^42–46^ we assume that the rich intrinsic dynamics of the local motor cortical circuit suffice to generate the complex patterns of cortical activity necessary to execute a given motor behavior. Which behavior is executed at a given time is selected by an upstream “motor command” that drives motor cortex via these upstream inputs. These inputs are therefore assumed to fluctuate on a much slower timescale than the motor cortical firing rates they drive. In the analysis presented below, we take these to be constant in time; results for more complex input dynamics are presented in Supplementary Materials Section S.1.6.

Motivated by the fact that motor behaviors are generally low-dimensional,^36–41^ we assume that the motor commands setting these inputs also have low dimensionality. We formalize this by representing the motor command as a *K*-dimensional vector, ***θ*** ∈ ℝ ^*K*^, constituted by *K ≪ N command variables θ*_1_, *θ*_2_, …, *θ*_*K*_. These command variables could correspond to extrinsic motor variables, such as reach speed or direction, or to more abstract motor-related information, such as parameters of prepared, observed, or imagined movements. Fundamentally, we make no commitments as to the nature of these intrinsic command variables beyond them influencing the upstream activity driving motor cortex. This assumption is formalized by having the upstream firing rates depend on the low-dimensional motor command via a set of encoding weights, *U*_*ij*_,

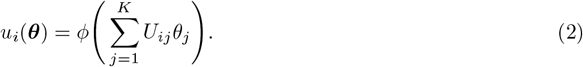

The rectified linear activation function, *ϕ*(·), is again used here to enforce non-negative firing rates. For simplicity, we set the encoding weights *U*_*ij*_ randomly.

During BCI control, motor cortical firing rates, **r**(*t*) = (*r*_1_(*t*) ···*r*_*N*_ (*t*)) ∈ ℝ ^*N*^, are directly translated to behavior of an external effector (e.g. a cursor on a screen) through a linear readout,

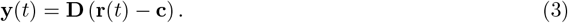

As is typically done in BCI experiments with linear decoders, we include a constant offset **c** to center the strictly positive firing rates (see Methods Section 4.8). The readout, **y**(*t*), determines behavior in the BCI task by specifying, for example, the position^47,48^ or velocity^11,12,14,15,49^ of a cursor. Regardless of how exactly how the readout maps to cursor movements, performing a given task (e.g. moving the cursor towards a target) demands a particular sequence of target readouts, which we denote by **y**^*^(*t*). A subject learning to perform a BCI task with a given decoder must therefore find a way to generate motor cortical activity patterns that will produce these target readouts.

Our hypothesis is that subjects do so only by optimizing the upstream motor commands, ***θ***. A key feature of this learning strategy is that it reduces the dimensionality of the learning problem. That reduction can be huge: from the number of synaptic weights to the number of command variables specifying the motor command, *K* – a factor that can easily reach 10^9^. Moreover, not all *K* command variables need to be optimized – we will argue below that, in certain settings, subjects may be optimizing only a subset of the task-relevant command variables, sometimes as few as 2. Such a reduction in the number of optimized parameters allows efficient learning in the absence of gradient information. However, it also limits the space of available solutions to the BCI task. Here we develop a formal theory of re-aiming to understand the implications of these limitations, and, importantly, show that they are consistent with empirical data.

We analyze a simplified model of re-aiming in which the motor command, ***θ***, is optimized to produce a target readout, **y**^*^, at a single endpoint time, *t*_end_,

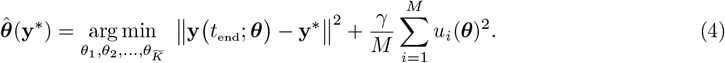

The vector **y**(*t*_end_; ***θ***) is the BCI readout at time *t*_end_ resulting from driving the model motor cortical network with the motor command ***θ***. The integer 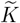 denotes the number of command variables optimized by re-aiming; for simplicity, the remaining command variables that are not optimized, 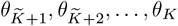, are set to 0. The second term on the right-hand side quantifies the metabolic cost of the upstream firing rates induced by the motor command, ***θ***, included in the objective function to ensure that only biologically plausible solutions are allowed.

Solutions to equation 4 constitute a concrete hypothesis about *what* subjects learn. In the following, we analyze these optimal motor commands to evaluate whether this hypothesis is consistent with empirical observations from BCI learning experiments. The question of *how* subjects might learn these optimal motor commands is left for future work. We also briefly acknowledge here that equation 4 constitutes an incomplete description of the true BCI learning problem, since controlling the BCI effector’s movement typically requires specifying a whole sequence of readouts over time (rather than at just one target time, *t*_end_) and relies on closed-loop feedback of the effector’s state.^50–52^ That said, this simplified model of re-aiming will prove useful to intuit general principles of the re-aiming learning strategy, which, as we show in Supplementary Materials Section S.1.6, extend to more complex settings such as closed-loop control. After all, being able to produce a target readout at a fixed future time is, loosely, a pre-requisite to solving the full closed-loop control problem.

### 2.2 Re-aiming implies neural constraints on short-term learning

We begin by modelling the BCI experiment designed by Sadtler et al. (2014).^14^ In this task, subjects learn to perform center-out movements with a 2D cursor on a screen, with the velocity of the cursor controlled by the readout from a linear BCI decoder, as in equation 3. Prior to learning, subjects first engage in a “calibration task”, in which neural activity is recorded while the subject passively views center-out cursor movements to eight radial targets (fig. 2a). Sadtler et al. observed that neural responses to these stimuli occupy a low-dimensional subspace, termed the “intrinsic manifold”. This subspace – identified via linear dimensionality reduction – is subsequently used to construct three types of BCI decoders.

**Figure 2:**
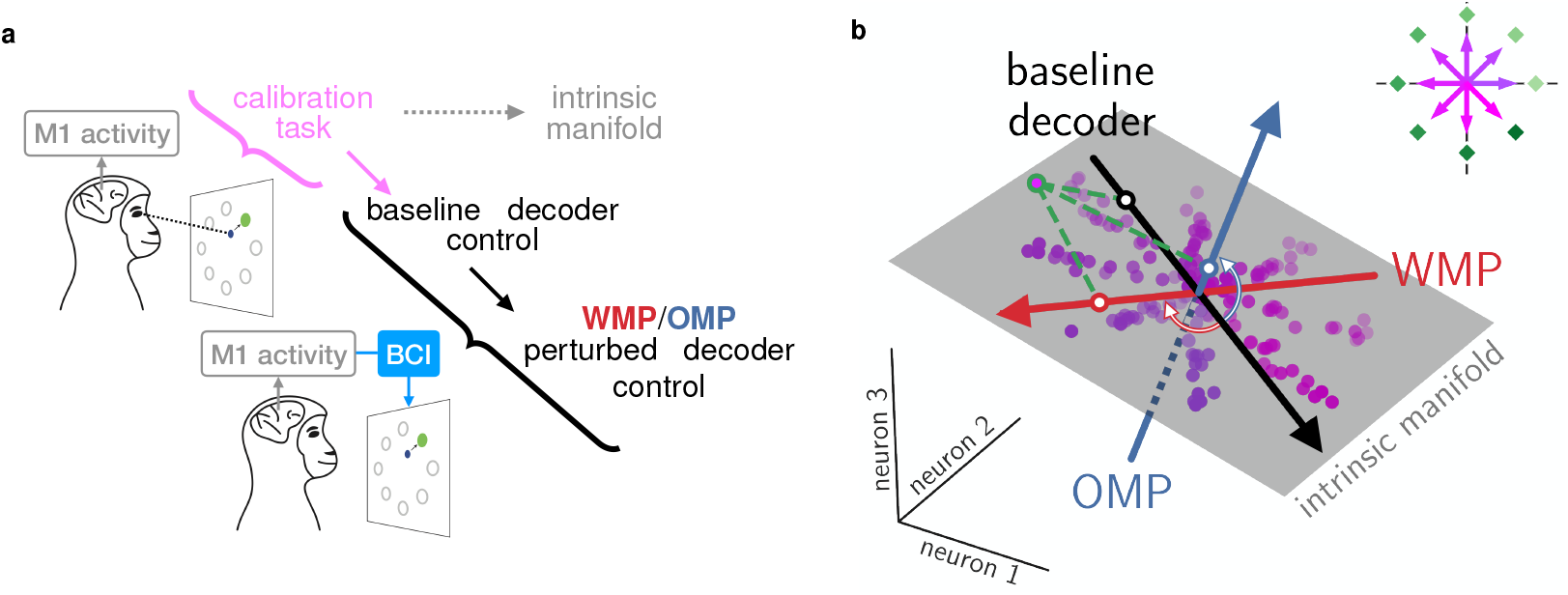
BCI learning task of Sadtler et al. (2014). a. Schematic of task structure. Subjects first engage in a “calibration task” whereby they passively observe center-out cursor movements on a screen. Recorded neural activity in motor cortex is used to construct the baseline decoder and estimate the intrinsic manifold. Subjects are then instructed to perform center-out cursor movements under BCI control, first using the baseline decoder and then with a perturbed decoder, constructed by perturbing the baseline decoder. This perturbation can either preserve the baseline decoder’s alignment with the intrinsic manifold (a within-manifold perturbation, or WMP) or disrupt it (an outside-manifold perturbation, or OMP). b. Low-dimensional illustration of the intrinsic manifold and its relationship to the decoders (defined in equation 3) used in this task. Colored dots represent activity patterns recorded during different trials of the calibration task, colored by the cursor velocity presented on that trial. The cursor velocities of these stimuli are depicted by color-matched arrows in the inset in the top right, with the cursor targets used in the subsequent cursor control task depicted by green diamonds. The evoked neural activity patterns reside predominantly within the two-dimensional plane depicted by the gray rectangle, the so-called intrinsic manifold. Three hypothetical one-dimensional decoders are depicted by colored arrows, labelled baseline decoder, WMP, and OMP. The corresponding component of the linear readouts, *y*_1_, from these decoders can be visualized by projecting individual activity patterns onto the corresponding decoder vector. This is illustrated for one activity pattern marked in green, whose projections onto each of the three decoders is shown. Because this activity pattern resides close to the intrinsic manifold, it yields a large readout (i.e. far from the origin, at the intersection of the three decoders) from the baseline decoder and WMP, which are both well aligned with the intrinsic manifold. In contrast, this activity pattern’s readout through the OMP is much weaker (i.e. its projection onto this decoder is much closer to the origin), since this decoder is oriented away from the intrinsic manifold. It is important to keep in mind that this illustration is a simplified cartoon of the true task, in which the intrinsic manifold is higher-dimensional (8-12D instead of 2D) and the BCI task depends on two readouts (*y*_1_, *y*_2_) rather than one.

First, a “baseline decoder” is constructed by fitting the decoding matrix, **D**, to the neural responses from the calibration task such that these activity patterns suffice to move the cursor towards the corresponding target in each trial. By construction, the baseline decoder is well aligned with the intrinsic manifold, such that activity patterns within this subspace can produce large readouts through this decoder (fig. 2b). Sadtler et al. found that, with this baseline decoder, non-human primate subjects can easily perform center-out cursor movements to the targets instantly, with no learning time required.

Next, the decoding matrix of the baseline decoder is perturbed and the subject is prompted to perform the same center-out cursor movements with the perturbed decoder. Two types of perturbations are used, which either preserve or disrupt the baseline decoder’s alignment with the intrinsic manifold: *within*-manifold perturbations (WMPs) randomly re-orient the baseline decoder *within* the intrinsic manifold, whereas *outside*-manifold perturbations (OMPs) randomly re-orient the baseline decoder *outside* the intrinsic manifold (fig. 2b). WMPs alter how neural activity *within* the intrinsic manifold subspace gets mapped to readouts, such that activity patterns in this subspace suffice to perform the task. Under an OMP, on the other hand, activity patterns within the intrinsic manifold are limited in the extent of readouts they can produce, so new activity patterns *outside* of the intrinsic manifold are needed to proficiently perform the task.

Sadtler et al. found that with 1-2 hours of practice (a few hundred trials), non-human primates can learn to successfully move the cursor to the targets with WMP decoders. In contrast, such short-term learning does not typically occur with OMP decoders, under which relatively little improvement is observed over this timespan. Here we argue that this limitation of short-term BCI learning is consistent with a re-aiming learning strategy. This low-dimensional learning strategy can account for the rapid learning achievable with WMPs, as well as the much slower learning exacted by OMPs.

To demonstrate this, we follow the experimental protocol outlined above, but with simulations of our motor cortical model (equations 1-2) rather than with animals. Our starting point, as with the experiments, is to estimate the intrinsic manifold from motor cortical firing rates recorded during the calibration task, in which the subject passively views center-out cursor movements to each of the eight radial targets, 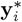.We simulated neural responses to these stimuli by driving the model network with command variables set to the cursor’s constant velocity on each trial: the first two command variables, *θ*_1_ and *θ*_2_, set to the coordinates of the given target, 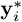,and the remaining command variables, *θ*_3_, *θ*_4_, …, *θ*_*K*_, set to 0. We then used Principal Components Analysis (PCA) to find the minimal subspace containing 95% of the variance over the resulting firing rates, which we found to be 8-dimensional (fig. 3g). We then defined the intrinsic manifold to be this subspace and used it to construct the baseline decoder and the two types of perturbed decoders (WMPs and OMPs), following the procedures of Sadtler et al. (see Methods Section 4.8).

**Figure 3:**
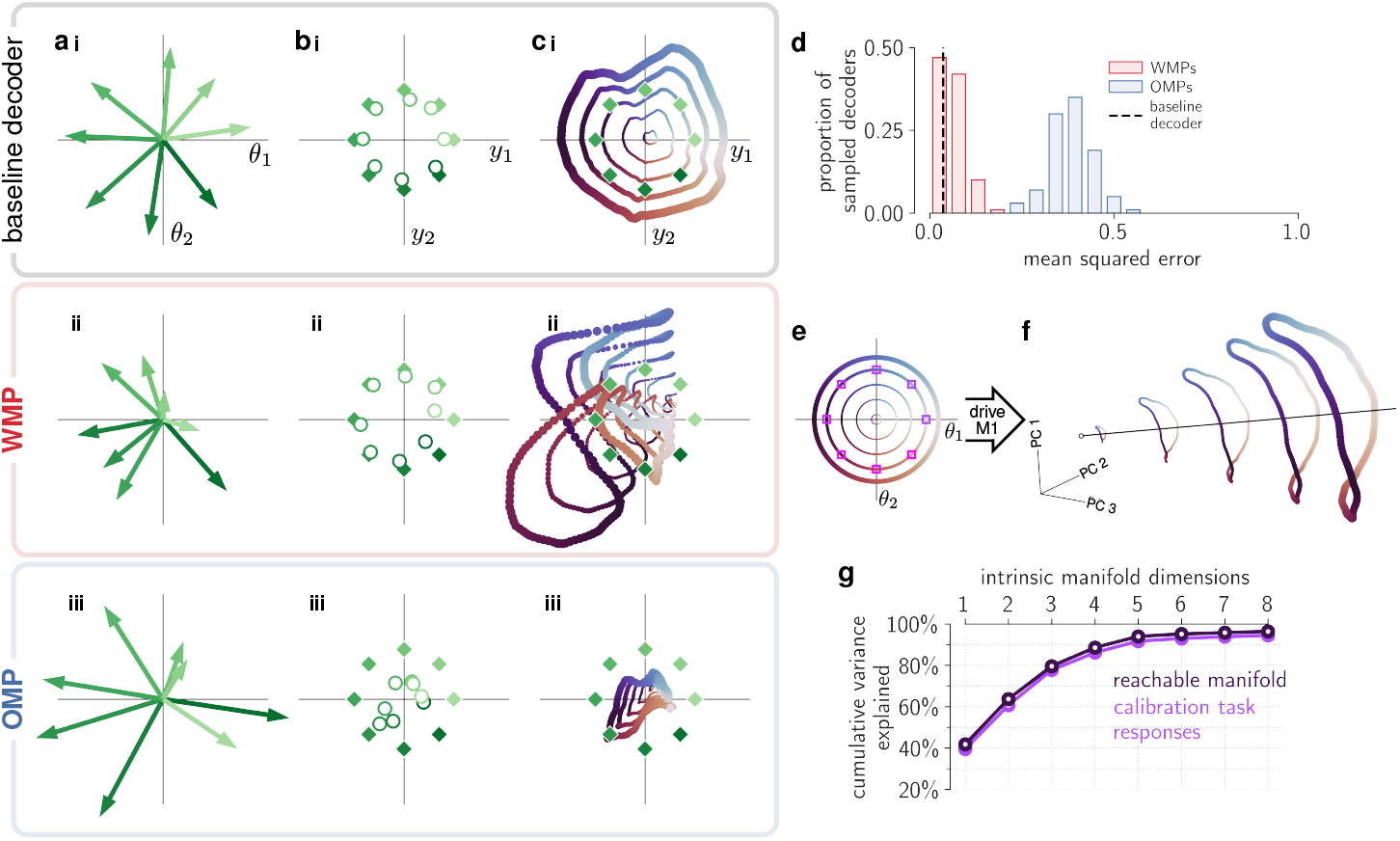
Re-aiming with two command variables suffices to learn good solutions for within-but not outside-manifold perturbations. a. Optimal motor commands, 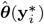,for the baseline decoder and one example WMP and OMP, plotted in *θ*_1_-*θ*_2_ space. The shade of green indexes the target readout, 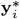,that each motor command is optimized for, corresponding to the target readouts plotted in the adjacent panel (green diamonds in fig. 3b). b. Readouts generated at time *t*_end_ by the optimal motor commands shown in the previous panel (fig. 3a), i.e. 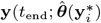.Green diamonds mark the eight target readouts for the center-out cursor control task, set to the directions of the eight radial cursor targets used by Sadtler et al. (2014). c. Readouts from each of the reachable manifold activity patterns plotted in fig. 3f, with matched marker colors and sizes. The diamonds denote the eight target readouts as in fig. 3b. Note that the reachable readouts closest to the targets do not necessarily match the readouts produced by the optimal motor commands (fig. 3b), as the optimal motor commands are optimized to minimize the metabolic cost of the upstream input as well the readout error (cf. equation 4). d. Distribution of mean squared error achieved by the optimal motor commands for 100 randomly sampled WMP’s and OMP’s. The mean squared error achieved by the optimal motor commands for the baseline decoder from which these perturbations are derived is marked by the vertical dashed black line. Target readouts are unit norm, so a mean squared error of 1.0 is equivalent to producing readouts at the origin. e. Motor commands covering a range of angles on the *θ*_1_ − *θ*_2_ plane and 5 norms, ∥***θ***∥ ∈ {0.1, 0.4, 0.7, 1.0, *s*_max_}, with *s*_max_ *≈* 1.25 (see Methods Section 4.4 for how this was chosen). The motor commands used to simulate the calibration task are indicated by the pink/purple squares. All other command variables, *θ*_3_, *θ*_4_, …, *θ*_*K*_, are fixed to 0. f. Activity patterns in the reachable manifold at endpoint time *t*_end_ = 1000ms. Each ring of activity patterns is generated by the corresponding ring of color- and size-matched motor commands in the previous panel. This ensemble of *N* - dimensional activity patterns is projected onto its top three principal components. The black line is drawn to facilitate visualization of the 3D structure of this conical manifold. Note that the points in this plot should not be thought of as spatiotemporal trajectories of activity; rather, they depict activity patterns *at the same timepoint* generated by different motor commands. g. Purple curve: cumulative variance in reachable manifold activity patterns along each intrinsic manifold dimension (equation 25). Gray curve: cumulative variance in calibration task neural responses. By construction, the intrinsic manifold contains 95% of the total variance of the calibration task neural responses (Methods Section 4.8).

Our hypothesis is that subjects learn to control the cursor by re-aiming with the same two command variables driving the calibration task responses, *θ*_1_ and *θ*_2_. We thus model BCI learning by optimizing *θ*_1_ and *θ*_2_ with respect to the re-aiming objection function (equation 4, with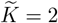), leaving the remaining command variables fixed to 0 as in the calibration task (*θ*_3_ = *θ*_4_ = … = *θ*_*K*_ = 0). As only two variables need to be optimized, learning should proceed very efficiently. The motor commands available for BCI control, however, are now severely constrained: only two command variables are free to change, and they are bounded by the metabolic cost incurred by the upstream firing rates (the second term in equation 4).

To see how this affects performance in this BCI learning task, we simulate re-aiming for WMP and OMP decoders. For each decoder and target readout, we solve equation 4 with 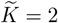 (setting *t*_end_ = 1000 ms, roughly matching the ∼700-1000 ms target acquisition times observed in experiments, and setting *γ* to its largest possible value guaranteeing good performance with the baseline decoder, cf. Methods Section 4.3) and drive the motor cortical network with the resulting optimal motor commands, 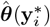.The optimal motor commands for the baseline decoder and an example WMP and OMP are shown in fig. 3a, as vectors in *θ*_1_-*θ*_2_ space. The readouts produced by driving motor cortex with these motor commands, 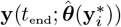,are shown in adjacent panels to the right (fig. 3b), with the corresponding target readouts underlaid. We find that, for the baseline decoder and WMP, most of these optimally driven readouts reach their targets; for the OMP, on the other hand, most of them fall far short. In fig. 3d, we repeat this simulation for 100 randomly sampled WMP and OMP decoders (see Methods Section 4.8 for the sampling procedure, closely matching that used by Sadtler et al.), and in each case quantify re-aiming success using the mean squared error between the optimally driven readouts and their corresponding targets. We find that the mean squared error is consistently lower for WMP decoders than for OMP decoders, as it was for the representative examples in fig. 3b.

Why does re-aiming fail to produce good readouts through OMPs? The answer lies in the constraints the re-aiming strategy imposes on the set activity patterns available in motor cortex for BCI control. To visualize and characterize these, we consider the *reachable manifold* : the set of all motor cortical activity patterns at a fixed endpoint time, **r**(*t*_end_; ***θ***), that can be reached by a motor command, ***θ***, which is accessible by re-aiming; that is, with *θ*_*k*_ = 0 for all *k >* 2 and with its norm, ∥***θ***∥, constrained by the quadratic metabolic cost (which we enforce here with a hard upper bound, ∥***θ***∥ ≤ *s*_max_, set to the maximum norm of the re-aiming solutions to all sampled decoder perturbations; cf. Methods Section 4.4). A large set of these accessible motor commands are shown in fig. 3e, and the corresponding activity patterns they generated are shown to the right in fig. 3f, projected down to three dimensions via PCA. Note that, despite the accessible motor commands being two-dimensional, the reachable manifold occupies more than two dimensions of state space, due to non-linearities in the dynamics of the motor cortical network. The three-dimensional projection in fig. 3f in fact contains about 80% of the variance over the *N* -dimensional activity patterns, revealing that the reachable manifold occupies in a moderately low-dimensional linear subspace – higher than that of the motor commands giving rise to it (two-dimensional) but significantly lower than that of its ambient state space (*N* -dimensional).

In fact, the reachable manifold is almost completely contained within the intrinsic manifold subspace. This is quantified in fig. 3g, which reveals that the eight dimensions of the intrinsic manifold subspace capture almost 100% of the variance in reachable activity patterns. This is unsurprising given that both the activity patterns in the reachable manifold and the activity patterns evoked by the calibration task – which define the intrinsic manifold – are generated by similarly low-dimensional motor commands ***θ***, in which only two command variables (*θ*_1_, *θ*_2_) are non-zero. Fig. 3e shows this directly by overlaying the calibration task inputs on the accessible motor commands. Ultimately, what this entails is that there are virtually no reachable activity patterns outside of the intrsinsic manifold; no activity patterns outside of the intrinsic manifold are accessible via re-aiming. This explains why this learning strategy would fail to produce large readouts through OMPs.

To confirm this, in fig. 3c we visualize the set of readouts reachable by re-aiming, for the baseline decoder, WMP, and OMP from fig. 3a and 3b. Specifically, we plot the readouts from each of the reachable activity patterns shown in fig. 3f, providing a comprehensive visualization of the space of readouts that can be reached through each decoder by re-aiming. As expected from the fact that the reachable manifold resides solely within the intrinsic manifold, we see that the readouts reachable under the baseline and WMP decoders cover a wider area than those reachable under the OMP decoder. The targets are thus enclosed by the baseline and WMP decoder reachable readouts, but remain out of reach of the OMP decoder. The re-aiming learning strategy therefore fails to solve the task with this OMP decoder, as none of the motor commands accessible under this learning strategy can reach the target readouts.

We conclude that re-aiming with only two variables (*θ*_1_ and *θ*_2_) can lead to successful BCI control with WMP decoders but not with OMP decoders. This offers an explanation for why only WMPs are learnable on the short timescale of a single experimental session. Because such low-dimensional re-aiming can’t succeed for OMPs, subjects must resort to an alternative – and presumably higher-dimensional – learning strategy, explaining why it requires substantially more training to learn these.^16^

### 2.3 Re-aiming predicts biases in short-term learning

A close look at fig. 3c reveals an important difference between the baseline and the WMP decoders: the readouts reachable with the baseline decoder cover the readout space symmetrically while those reachable with the WMP decoder do not (compare figs. 3ci and 3cii). In other words, larger readouts are reachable in some directions than in others. Such biases in reachable readouts are not unique to this particular WMP decoder; fig. 4a reveals similar asymmetries in the readouts reachable through three other representative WMP decoders.

**Figure 4:**
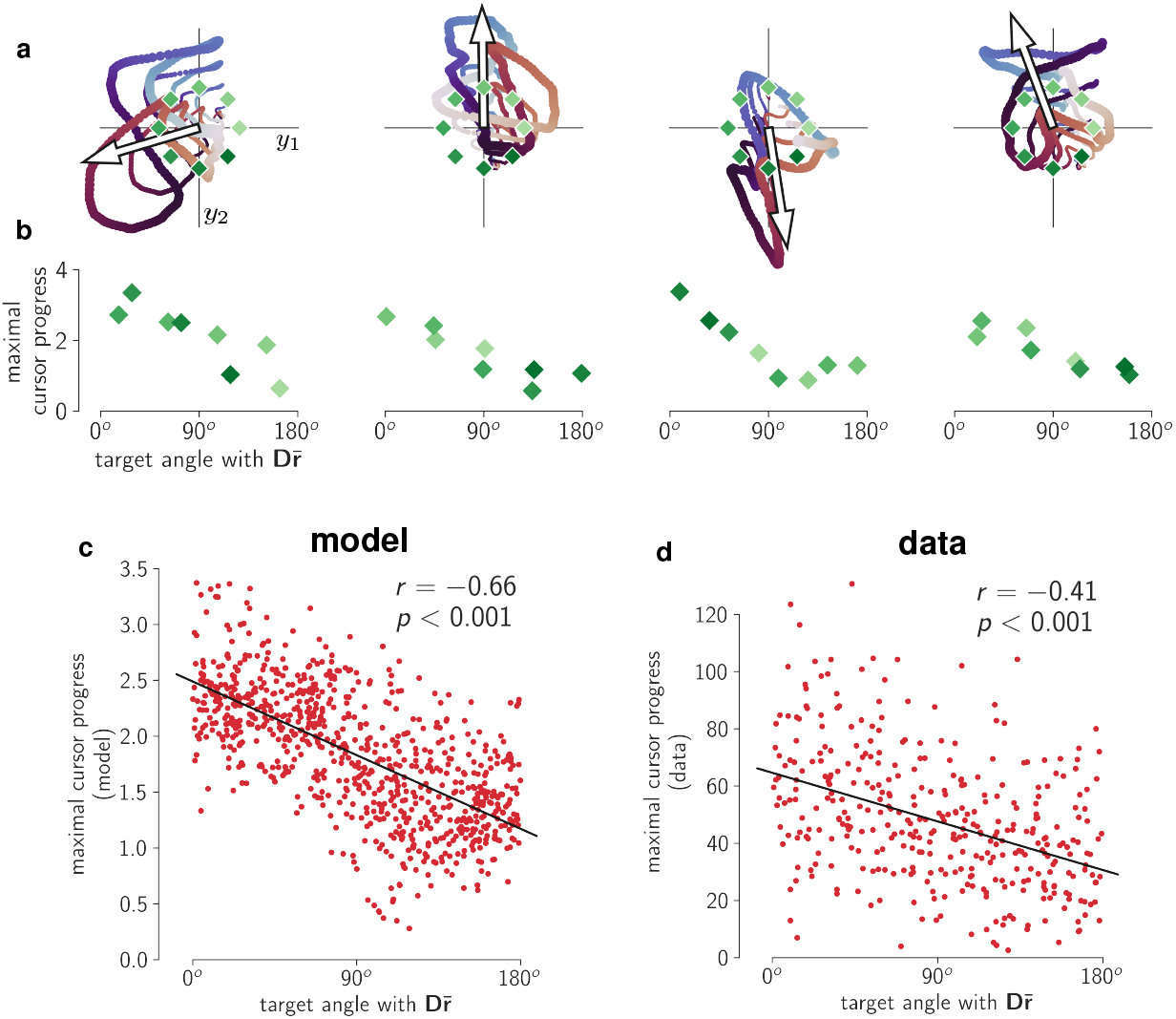
Re-aiming predicts biases in readouts after short-term learning of within-manifold perturbation (WMP) decoders. a. Readouts reachable through four representative WMP decoders, using the same color conventions as in fig. 3c. In each case, the four loops correspond to four distinct motor command norms, chosen to aid visualization. The leftmost panel corresponds to the example WMP decoder shown in fig. 3cii. The projection of the reachable manifold centroid, 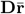,is overlaid as an open arrow, arbitrarily rescaled for visibility. b. Maximal cursor progress in each target direction as a function of angle with 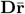, for the four example WMP decoders in panel a. c. Maximal cursor progress in each target direction as a function of angle with 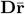,for all 100 sampled WMPs (Pearson *r* = −0.66, *p <* .001, *N* = 8 target directions × 100 sampled WMPs = 800). As was done for the experimental data in the next panel, the reachable manifold centroid, 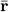, is estimated using simulated mean firing rates during baseline decoder control (see Methods Section 4.6). d. Maximal cursor progress in each target direction as a function of angle with 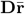,for all 46 sessions of WMP learning across three monkeys (Pearson *r* = −0.41, *p <* .001, *N* = 8 target directions × 46 experimental sessions with WMP control = 368). Maximal cursor progress is estimated using the average cursor progress over the 50 contiguous trials with lowest acquisition times. The reachable manifold centroid, 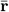, is estimated using mean firing rates over trials of baseline decoder control (see Methods Section 4.6).

The direction of this bias is moreover predictable: typically, the largest reachable readouts are in the direction of 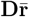 (arrow overlaid on each plot), where 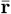 is the centroid of the reachable manifold. This bias arises because of the non-negativity of firing rates, which permits the population firing rate, **r**, to grow widely away from the origin, but shrink towards the origin only up to a point, where it is truncated by the non-negativity. It is this property that endows the reachable manifold its conical structure (fig. 3f), whose centroid, 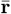, dictates the direction in which firing rates can grow the most under the re-aiming strategy. The projection of this direction through a given decoder, 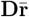,thus determines the direction in which the largest readouts can be reached by re-aiming (see Supplementary Materials Section S.1.2 for a more detailed analysis). In Supplementary Figure S1d, we show that – as long as firing rates are constrained to be non-negative (Supplementary Materials Section S.1.3) – this bias arises across a large variety of motor cortical connectivity patterns and dynamics, suggesting that it is an unavoidable consequence of the re-aiming learning strategy. The absence of such a bias in experimental data would therefore provide strong evidence against this theory of BCI learning.

To quantify this experimental prediction, we used the “cursor progress” metric, *ρ*, introduced by Golub et al. (2018) to measure the degree to which a given readout, **y**, pushes the BCI cursor in a given target direction, **y**^*^,

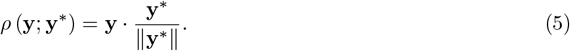

We then predict the maximum achievable cursor progress in each target direction,

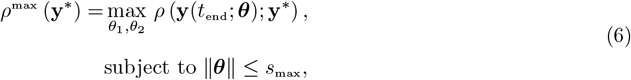

where, as above, *θ*_1_ and *θ*_2_ are the two command variables optimized by re-aiming and *s*_max_ is the bound on motor command norms imposed by the metabolic constraint in equation 4 (cf. Methods Section 4.4). In fig. 4b, we plot this maximal cursor progress for each target readout, 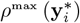,as a function of the target readout’s angle from 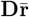,for each of the four example WMPs. The negative correlation in each case confirms our above observation: higher cursor progress is reachable in target directions more aligned with 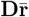.In fig. 4c, we plot the maximal cursor progress in each target direction for all 100 sampled WMP decoders, revealing a statistically significant negative correlation across all sampled decoders (Pearson *r* = −0.66, *p <* .001).

Does this predicted negative correlation also hold in the empirical data? To test this, we estimated the maximal cursor progress and reachable manifold centroid in each experimental session of WMP control. Maximal cursor progress in each target direction, 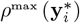,was estimated using the average cursor progress in that target direction over the 50 contiguous WMP control trials with fastest target acquisition times (see Methods Section 4.6). The reachable manifold centroid, 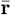,was estimated using mean motor cortical firing rates during the block of baseline decoder control (see Methods Section 4.6), which in our model is highly correlated with the true reachable manifold centroid. We then replicated fig. 4c by plotting the empirically measured maximal cursor progress for each target direction as a function of the angle between the target direction and 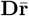,using our empirical estimates of maximal cursor progress and 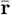 from each experimental session. The data over all sessions are plotted in fig. 4d, revealing a significant negative correlation (Pearson *r* = −0.41, *p <* .001) akin to that observed in our model. This confirms the existence of a statistically significant bias in the same direction predicted by our model of re-aiming.

### 2.4 Long-term BCI learning by generalized re-aiming

Although non-human primates struggle to control OMP decoders within a single experimental session (a few hundred trials),^14^ they can in fact learn to do so when trained over multiple days (thousands of trials).^16^ In this long-term learning paradigm, new motor cortical activity patterns emerge that allow the subjects to achieve good performance with OMP decoders. Could re-aiming play a role in the emergence of novel activity patterns over these longer timescales?

Since re-aiming with the two command variables evoked by the calibration task, *θ*_1_ and *θ*_2_, is not sufficient to produce the activity patterns required for OMP control, additional command variables will be required. We refer to a learning strategy that uses additional command variables as “generalized re-aiming”, and demonstrate below that this strategy can in fact achieve good performance with OMP decoders. Moreover, it can account for why learning is slower for these decoders: the search for optimal motor commands takes place in a higher-dimensional space beyond the narrow 2D space of command variables evoked by the calibration task.

To simulate generalized re-aiming, we simply increase the number of command variables used for re-aiming, 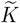,and solve the resulting 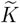 -dimensional optimization problem in equation 4. In fig. 5a we plot the mean squared error achieved by the re-aiming solutions for each OMP decoder for each value of 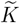.We find that as 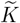 increases, a lower mean squared error is achieved, demonstrating that this learning strategy can be effective for OMP learning. For this model motor cortical network, re-aiming with about 15-20 command variables suffice to achieve a mean squared error as low as that achievable with WMP decoders using 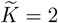.For other motor cortical models with different connectivity, fewer than 10 command variables suffice (Supplementary Figure S1e). These values of 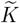 comfortably fall in the range of the total number of extrinsic motor variables known to influence motor cortical activity.^53–59^ However, they may be too high for naïve gradient-free optimization to succeed in solving equation 4 under biological limitations (e.g. on memory, motivation, and noise), which might explain why primates seem to only be able to learn to control OMP decoders when provided with a structured incremental training paradigm.^16^

**Figure 5:**
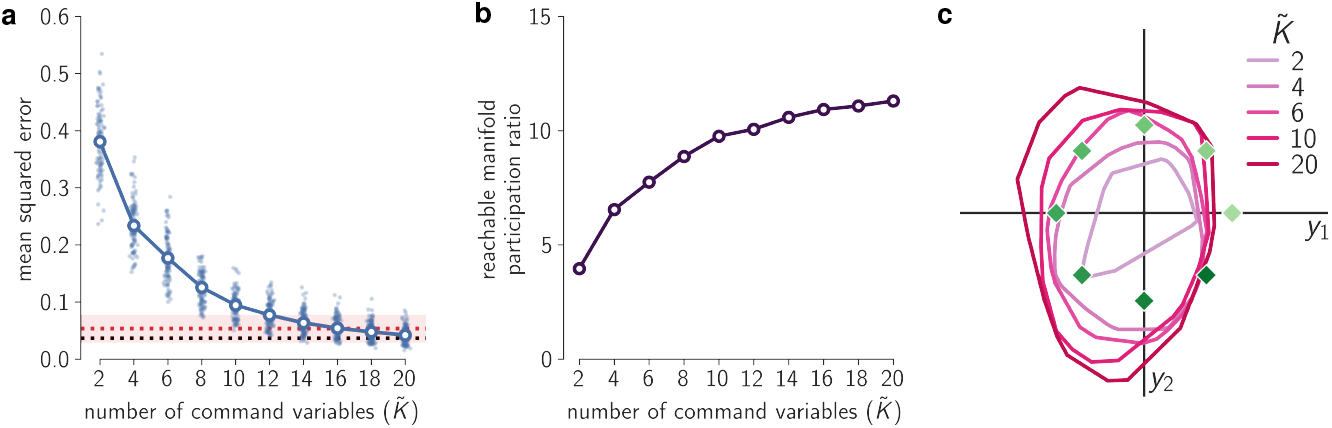
Generalized re-aiming produces good solutions for outside-manifold perturbations (OMPs). a. Mean squared error achieved by generalized re-aiming solutions for all sampled OMP decoders, plotted as a function of the number of command variables used for re-aiming,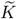. Lighter blue points show the mean squared error for individual OMP decoders, darker open circles show the median over all sampled OMPs. For reference, dotted horizontal lines show the mean squared error achieved by re-aiming solutions with 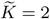 for the baseline decoder (black) and for WMP decoders (red); for WMP decoders, the median over all sampled decoders is shown with shading marking the upper and lower quartiles (corresponding to the values plotted in the red histogram in fig. 3d). b. Participation ratio of the reachable manifold covariance (a measure of the effective dimensionality of the reachable manifold; see Methods Section 4.4, equation 16) as a function of the number of command variables used for re-aiming, 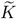. c. Convex hull of OMP readouts reachable with different number of command variables, 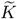,for the same OMP decoder shown in fig. 3c. The innermost ring 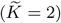 corresponds to the convex hull of the reachable readouts plotted in fig. 3ciii.

Why generalized re-aiming works can be understood by looking at how increasing 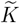 changes the set of activity patterns reachable by re-aiming. A larger number of learnable command variables permits a more diverse set of upstream inputs, which in turn implies that a more diverse set of activity patterns are reachable. This diversity is quantified in fig. 5b by the participation ratio of the covariance of the reachable manifold (see Methods Section 4.4, equation 17). The participation ratio measures the extent to which variability is spread out over many dimensions (high participation ratio) or concentrated to only a few (low participation ratio).^60^ We find that as 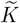 rises, the participation ratio of the reachable manifold covariance increases, indicating it occupies more and more dimensions of state space. That said, the participation ratio does begin to saturate at around 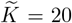,reflecting the fact that the reachable manifold is ultimately limited by the smooth dynamics of the motor cortical network.

This expansion in the reachable manifold leads to the inclusion of new activity patterns that are useful for OMP control. We can see this in fig. 5c, which shows the readouts reachable through the same OMP visualized in fig. 3ciii. The readouts reachable under different values of 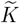 are plotted with different colors, revealing how re-aiming with a larger number of command variables allows the target readouts to be reached. As the reachable manifold expands, more and more activity patterns occupying dimensions relevant to OMP control become reachable, such that a wider set of readouts become reachable.

### 2.5 Illusory credit assignment by generalized re-aiming

We now turn to a different class of BCI decoder perturbation, termed the credit assignment rotation perturbation.^61^ We can think of the readout from a linear BCI decoder (equation 3) as summing together the *N* columns of the decoding matrix **D** (termed the “decoding vectors”), each one weighted by the activity of the corresponding neuron (fig. 6a, top). Under a credit assignment rotation perturbation, the decoding vectors of a random subset of neurons (the “rotated neurons”) are rotated by a given angle (fig. 6a, bottom). Errors induced by this decoder perturbation can be corrected by adjusting only the responses of the rotated neurons, while leaving the responses of the “non-rotated” neurons unchanged. But doing so would require solving the so-called credit assignment problem:^62^ identifying which neurons’ decoding vectors were rotated – a tall order given that the subject has no explicit knowledge about the BCI decoder or the few motor cortical neurons (among millions) it records from.

**Figure 6:**
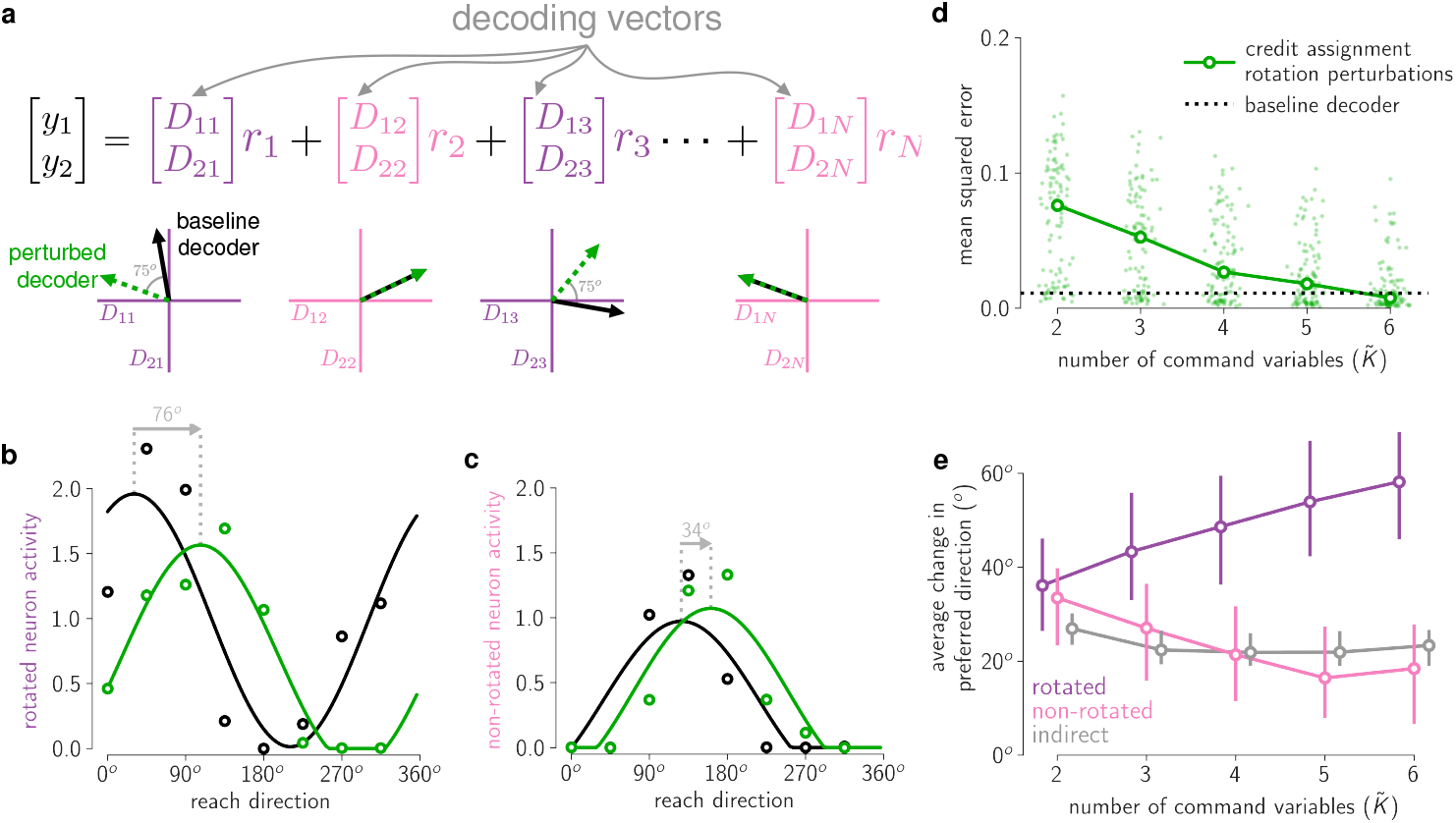
Generalized re-aiming solutions reproduce motor cortical tuning changes observed under credit assignment rotation perturbations. a. A linear BCI readout (equation 3) can be interpreted as summing together columns of the decoding matrix **D**, each weighted by the firing rate of the corresponding neuron (the centering term **c** has been dropped here for simplicity). These columns are called the neurons’ decoding vectors, and they are plotted on the axes below the equation. Under a credit assignment rotation perturbation, the decoding vectors of a subset of neurons (marked in purple) are rotated by a fixed angle (in this case 75^*o*^ counter-clockwise). The neurons’ decoding vectors under this perturbed decoder are shown by dashed green arrows. The neurons whose decoding vectors are rotated are termed “rotated” neurons (in purple), the rest of the neurons that are recorded by the BCI are termed “non-rotated” neurons (in pink). Neurons that are not recorded by the BCI (i.e. whose decoding vectors are just a vector of 0’s, not depicted here) are termed “indirect” neurons. b. Tuning curve of a representative example rotated neuron of our model, during cursor control with the baseline decoder (black) and with a credit assignment rotation perturbation (green). The dots show the time-averaged activity over *t*_end_ = 1000ms while the motor cortical network is driven by the re-aiming solutions for each respective decoder, using 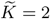 for the baseline decoder and 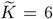 for the perturbed decoder. Curves show tuning curves fit to these responses (Methods Section 4.9). The vertical dotted gray lines mark the preferred direction under each decoder, with an arrow labeling the change in preferred direction. c. Tuning curve of a representative example non-rotated neuron of our model, under the same two decoders. All conventions exactly as in the previous panel. Note that this neuron’s preferred direction changes less than that of the rotated neuron in the previous panel. d. Mean squared error achieved by generalized re-aiming solutions for 100 random credit assignment rotation perturbations, plotted as a function of the number of command variables used for re-aiming, 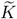.Light green dots denote individual decoder perturbations, overlaid darker open circles denote medians over all 100 sampled decoder perturbations. Black dotted horizontal line shows the mean squared error achieved by re-aiming solutions to the unperturbed baseline decoder with 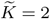. e. Average change in preferred direction of rotated, non-rotated, and indirect neurons between simulated cursor control with the baseline decoder and each perturbed decoder, plotted as a function of the number of command variables used for re-aiming, 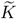.For each decoder perturbation, the changes in preferred direction are averaged over all neurons in each sub-population, and the median over all sampled perturbations is plotted. Error bars mark the upper and lower quartiles. Positive angles indicate a counter-clockwise rotatation, consistent with the direction of rotation of the decoding vectors of the rotated neurons.

Despite these challenges, multiple studies have shown that non-human primates can learn to control such decoder perturbations.^11,12,61^ These studies used the same 2D cursor control task described above (fig. 2a), in which subjects first control a BCI cursor using a baseline decoder fit to motor cortical activity recorded during a calibration task, and then learn to control the cursor using a perturbed decoder with rotated decoding vectors. Subjects’ motor cortical activity changes after learning to control the perturbed decoder, and this can be characterized by the change in neurons’ tuning to cursor direction during BCI control with the baseline and perturbed decoders. Each of these studies found that, after learning, tuning curves of both rotated (fig. 6b) and non-rotated neurons (fig. 6c) shift in the same direction as the decoding vectors. For example, if the decoding vectors are rotated counter-clockwise, tuning curves also shift counter-clockwise. Notably, however, tuning curves of rotated neurons shift more on average than those of non-rotated neurons (compare the simulated examples in fig. 6b and fig. 6c). This observation could be interpreted to support the hypothesis that the motor system is able to solve the credit assignment problem: it has identified which neurons’ decoding vectors were rotated, and accordingly modified their responses more so than the others’ (e.g. via Hebbian plasticity^17^). Here we consider an alternate hypothesis: that this phenomenon could arise from generalized re-aiming, a global learning strategy entirely unconcerned with modifying individual neurons.

To test this, we follow the same procedure as above: we simulate motor cortical activity during the calibration task, use it to construct a baseline decoder, and then sample 100 random credit assignment rotation perturbations (Methods Section 4.9). Following the experimental procedures of Zhou et al. (2019), the perturbed decoders are constructed by applying a 75° counter-clockwise rotation to a random selection of 50% of the columns of the decoding matrix, **D**. For each decoder, we then compute the optimal motor commands for each target (equation 4) and drive the network with them to simulate center-out cursor movements learned by re-aiming. By comparing neurons’ directional tuning under the optimal motor commands for the baseline decoder and the perturbed decoder, we can determine how directional tuning would change after learning by re-aiming.

Reflecting the fact that the baseline decoder is easy to learn, we used 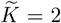 to compute re-aiming solutions for it. For the perturbed decoders, we simulated generalized re-aiming with 2 to 6 command variables. We find that re-aiming with about 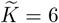 command variables is necessary to achieve the same performance as with the baseline decoder (fig. 6d). That said, re-aiming with only 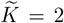 suffices to achieve a relatively low mean squared error (around 0.1; compare to fig. 3d), suggesting ordinary 2D re-aiming could still be a viable learning strategy for this task.

As in the experiments, we measured neurons’ preferred directions (i.e. the direction at the tuning curve peak, cf. fig. 6b and fig. 6c) under the optimal motor commands for the baseline decoder and for each perturbed decoder, and calculated each neuron’s change in preferred direction. We then averaged the change in preferred direction separately over rotated and non-rotated neurons. Figure 6e shows the median of this average change in preferred direction over all sampled perturbed decoders, as a function of the number of command variables used for re-aiming, 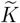.Consistent with the experimental results, we find that re-aiming leads to a global counter-clockwise shift in motor cortical tuning curves congruent with the rotation of the decoding vectors. Importantly, we find that generalized re-aiming with 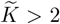 command variables replicates the credit assignment effects seen in the experiments, whereby the preferred directions of rotated neurons shift on average more than their non-rotated counterparts. This is true despite the fact that the credit assignment problem was never truly solved: no neuron-specific parameters were modified under this learning strategy.

Because we have complete access to the full population of neurons in our motor cortical model, we can also measure tuning changes in the sub-population of “indirect” neurons not recorded by the BCI (i.e. neurons whose decoding vectors in **D** comprise a vector of 0’s). These are plotted in fig. 6e with a gray line. Under generalized re-aiming, indirect neurons’ tuning curves shift less on average than rotated neurons’. Whether they shift more or less than non-rotated neurons’ tuning curves, on the other hand, depends on the specific value of 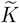 and varies considerably across different perturbed decoders.

These two results are roughly consistent with the observations of Zhou et al. (2019), who examined this phenomenon in two non-human primate subjects. They found that, in both subjects, the average change in tuning curves was larger for rotated neurons than for indirect neurons, as predicted by our model. But when comparing non-rotated neurons to indirect neurons, they found that the average change in tuning curves was larger for the former in one subject and larger for the latter in the second subject, consistent with the variability observed in our simulations. To our knowledge, this is the only experimental study on indirect neurons’ responses before and after learning a credit assignment rotation perturbation; more studies are needed to fully test the predictions of our model.

An important additional prediction of our model is that credit assignment effects do not arise under ordinary 2D re-aiming (fig. 6e,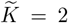). This is consistent with prior modeling work showing that two-dimensional re-aiming does not suffice to account for empirically observed changes in motor cortical tuning curves after learning a credit assignment perturbation.^33^ Interestingly, it is also consistent with recent experimental work showing that differences between rotated and non-rotated neurons seem to arise gradually over multiple days of training.^61^ Our model suggests that this timecourse of learning reflects a change in learning strategy, whereby subjects initially engage in low-dimensional re-aiming to rapidly reduce gross cursor movement errors, and then turn to generalized re-aiming to further refine

BCI control over a longer timescale, resulting in more marked credit assignment effects later in learning. We briefly remark, however, that Zhou et al. did not observe changes in the preferred directions of non-rotated neurons after the first day of training. In our simulation, on the other hand, the non-rotated neuron tuning curves shift back towards their starting values under the baseline decoder as the number of re-aimed command variables increases (see decreasing pink line in fig. 6e). This discrepancy between our model and the experimental data could be explained by subjects using suboptimal re-aiming solutions deviating from the optimal one predicted by our model, possibly due to the difficulty of solving this optimization problem when 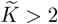,or due to the lack of motivation to find it (since even 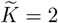 suffices to achieve relatively low error, fig. 6d). This could also explain why the amount of change in preferred directions is significantly larger in our simulation (30 − 60^*o*^) than in the experimental data (20 − 40^*o*^).

### 2.6 Operant conditioning of individual neurons by re-aiming

The third and final BCI task we study is the operant conditioning of individual motor cortical neurons. In this task, subjects are rewarded for simply increasing the activity of one group of motor cortical neurons over another.^28,63–66^ The fact that primates and rodents are capable of solving such tasks is often cited as evidence that the motor system can learn to specifically modulate the responses of individual neurons. Classical models of single-neuron operant conditioning have argued that these changes happen via reward-modulated plasticity at their synapses.^18,67^ Here we explore the extent to which these observations could instead be explained by re-aiming.

We begin by considering the classic operant conditioning task of Fetz and Baker (1973). In this task, the subject is rewarded for increasing the firing rate of one neuron – termed the “target” neuron – while simultaneously decreasing that of another neuron – termed the “distractor” neuron. Remarkably, Fetz and Baker found that non-human primates are often able to do this with only minutes of practice. Moreover, the identity of the target and distractor neurons could be flipped midway through a recording session, and the subject would subsequently adapt to this new target assignment within tens of minutes, increasing the activity of the neuron whose activity was previously suppressed. Could low-dimensional re-aiming explain this behavior?

The answer depends on the reachable manifold. If the reachable manifold contains activity patterns in which neuron *a* is more active than neuron *b*, as well as activity patterns in which neuron *b* is more active than neuron *a*, then good re-aiming solutions will exist for both target assignments. This is illustrated in fig. 7a, which shows two neurons’ endpoint firing rates, *r*_*i*_(*t*_end_), at various points on the reachable manifold, following the same conventions as in fig. 3f. On this plane, activity patterns below the diagonal are rewarded when neuron *a* is the target neuron; activity patterns above the diagonal are rewarded when neuron *b* is the target neuron. Because there are reachable activity patterns on both sides of the diagonal, good re-aiming solutions exist for both target assignments. We calculated optimal re-aiming solutions for each target assignment by maximizing the firing rate difference between the target and distractor neurons, subject to a metabolic cost (equation 29). The activity produced by these optimal solutions are marked by the open red and green circles. These evidently satisfy each target assignment, with neuron *a* achieving a higher firing rate under one re-aiming solution (green circle) and neuron *b* achieving a higher firing rate under the other (red circle).

**Figure 7:**
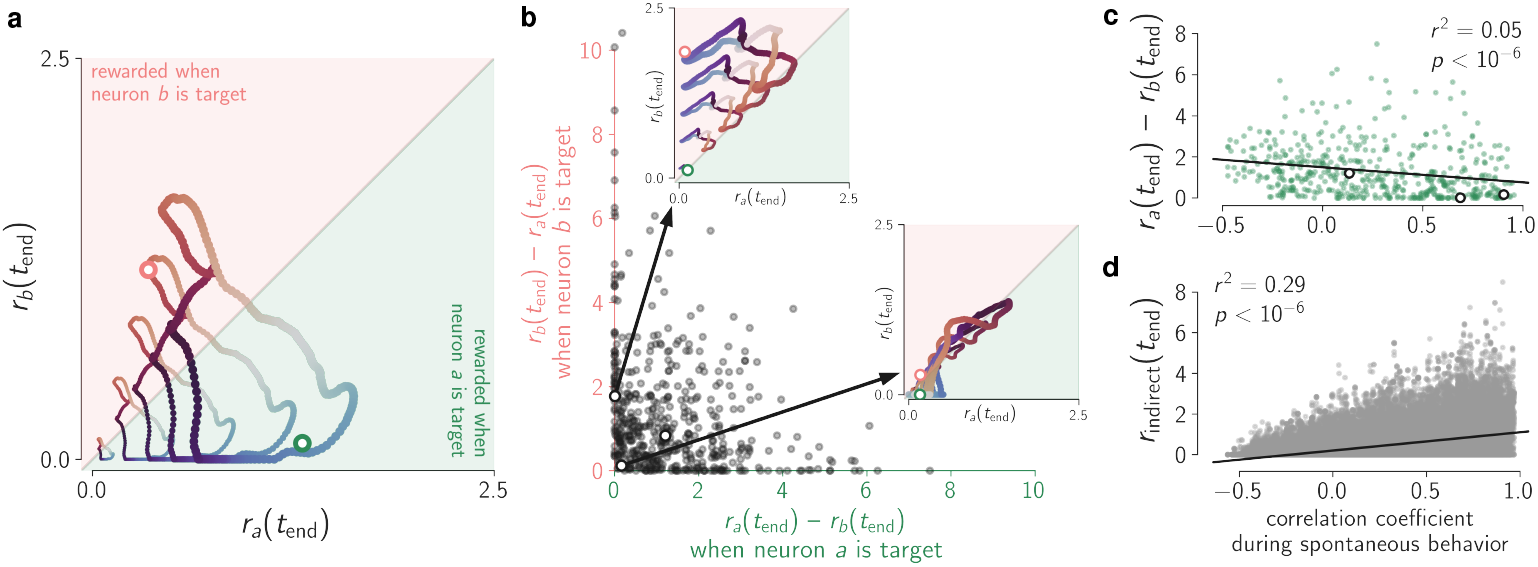
Operant conditioning of individual neurons by re-aiming. a. Activity of two model neurons at various points on the reachable manifold, at *t*_end_ = 1000ms with 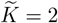, following the same conventions as fig. 3f. As in that figure, each ring of activity patterns is generated by the corresponding ring of color- and size-matched motor commands in fig. 3e. Activity patterns below the diagonal are ones where neuron *a* is more active than neuron *b*, satisfying the task demands when neuron *a* is the target neuron; the reverse holds for activity patterns above the diagonal. The green and orange open circles denote the activity patterns produced by the optimal re-aiming solutions for the two respective target assignments. Note that, due to the metabolic cost term in the re-aiming objective function, these do not necessarily correspond to the points on reachable manifold that are furthest away from the diagonal. b. Difference in activity produced by re-aiming solutions for each target assignment (at time *t*_end_ = 1000 ms, the endpoint time the re-aiming solutions were optimized for), for 500 random pairs of neurons from the same model motor cortical network used in previous simulations. Each dot corresponds to one pair of neurons. The pair of neurons shown in previous panel is marked by an open circle. Two additional examples are marked by open circles. Insets show the activity of those neuron pairs at various points on the reachable manifold, following same conventions as the previous panel. c. Difference in activity (at *t*_end_ = 1000 ms) produced by the re-aiming solutions optimized for neuron *a* being the target, for the same 500 random pairs of neurons, plotted as a function of the correlation between the two neurons during simulated spontaneous behavior. The three example pairs of neurons highlighted in the previous panel are again highlighted here with open circles. A quantitatively similar trend is observed for *r*_*b*_ − *r*_*a*_ with re-aiming solutions optimized for neuron *b* being the target (data not shown). d. Activity of indirect neurons (at *t*_end_ = 1000 ms) for the same neuron pairs and re-aiming solutions in previous panel, plotted as a function of correlation with neuron *a* during simulated spontaneous behavior. A quantitatively similar trend is observed for correlations with neuron *b* with re-aiming solutions optimized for neuron *b* being the target (data not shown).

More generally, we can think of this plot as a particular two-dimensional projection of the reachable manifold, specified by the pair of neurons *a* and *b*. A given neuron pair thus admits good re-aiming solutions for this task whenever the corresponding projection of the reachable manifold covers the appropriate side(s) of the diagonal. Framed in this way, it is easy to intuit that for most random pairs of neurons a good solution will generally exist for at least one target assignment – random two-dimensional projections of the reachable manifold are unlikely to lie exactly on the diagonal. We verify this intuition by sampling 500 random pairs of neurons from our model motor cortical network (see Methods Section 4.10) and checking whether the re-aiming solutions for the two target assignments produce higher firing rates for the target neuron than for the distractor neuron. The difference in the two neurons’ firing rates produced by the optimal re-aiming solutions are plotted in fig. 7b. For most neuron pairs, we see that at least one of the two neurons can be activated more than the other. In many cases, both neurons can be activated more than the other, meaning that both target assignments could be learned by re-aiming.

For some neuron pairs, however, the optimal re-aiming solutions do not produce a large difference in firing rates under either target assignment. These infelicitous neuron pairs are ones where the two neurons are highly correlated across all activity patterns on the reachable manifold, such that the relevant projection doesn’t deviate strongly from the diagonal and no good re-aiming solutions exist. One such example is shown in the inset on the right, where the reachable manifold lies largely right on the diagonal, and thus both re-aiming solutions lead to near-0 difference in firing rates. A different kind of exampe is shown in the inset on the top, where the two neurons are correlated in such a way that the reachable manifold resides on only one side of the diagonal. In this case, a good re-aiming solution exists for one target assignment but not for the other. These observations reveals a tight relationship between neural correlations and operant conditioning performance: the more correlated a pair of neurons is across reachable activity patterns, the more difficult it should be to selectively activate one more than the other by re-aiming.

Is this prediction of our model consistent with experimental observations? Without empirical access to the reachable manifold, we cannot directly measure correlations across reachable activity patterns. But if we assume that neural activity during a prior “calibration task” is driven by the same command variables used subsequently for re-aiming – as we did in our simulation of WMP/OMP learning –, then we should expect neural correlations during this calibration task to approximately match correlations across the reachable manifold used for re-aiming. This predicts that neural correlations during a prior calibration task should be predictive of subsequent operant conditioning performance. This prediction is in fact consistent with observations from a study by Clancy et al. (2014), in which the conditioned neurons’ correlation was measured during a period of spontaneous behavior (the “calibration task”) just prior to performing operant conditioning. Consistent with our model’s prediction, they observed that the stronger the correlation during spontaneous behavior prior to operant conditioning, the worse the mouse tended to perform the subsequent operant conditioning task.

To quantify this prediction, we simulated the experiment of Clancy et al. We first simulated motor cortical activity during spontaneous behavior by driving the model motor cortical network with randomly sampled motor commands in which only two command variables, *θ*_1_ and *θ*_2_, were allowed to vary (all other command variables were set to 0). We sampled 50 such motor commands to simulate 50 bouts of spontaneous activity (cf. Methods Section 4.10). For each conditioned pair of neurons, we measured their correlation coefficient over all activity across the 50 bouts, and then simulated re-aiming with the same two command variables, *θ*_1_ and *θ*_2_. We quantified the efficacy of re-aiming with the firing rate difference between the target and distractor neurons at the optimized endpoint time, *t*_end_. Mirroring the experimental results of Clancy et al., we find that the spontaneous activity correlations are weakly but significantly predictive of re-aiming efficacy (fig. 7c). We briefly remark here that the operant conditioning task of Clancy et al. differs from our simulations in that pairs of ensembles of up to 11 neurons were conditioned, rather than pairs of single neurons. In Supplementary Materials Section S.1.4, we show that the same results hold in this setting as well.

Finally, we consider what happens to the “indirect” neurons – neurons that are neither a target nor a distractor. Clancy et al. observed that, after learning, indirect neurons *strongly* correlated with the target neuron during spontaneous behavior remained highly active during performance of the subsequent operant conditioning task (supplementary figure 9a in^65^). This is consistent with our model of re-aiming, in which re-aiming tends to drive indirect neurons proportionally to their correlation with the target neuron during spontaneous behavior (fig. 7d). Our model is not, however, consistent with another observation by Clancy et al.: indirect neurons that were *moderately* correlated with the target neuron were highly active only in the early stages of learning, but by the end of learning became silent. A possible explanation for this inconsistency is that subjects re-aim with 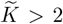 command variables. If spontaneous behavior during the calibration period were driven by more than 2 command variables, we would expect subjects to re-aim with more as well. Given the complexity of spontaneous behavior, this is a reasonable explanation, but we leave for future work a more comprehensive study of generalized re-aiming with 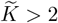 command variables in operant conditioning tasks.

## Discussion

In this study, we proposed and investigated the hypothesis that motor cortical BCI learning proceeds via a learning strategy we refer to as generalized re-aiming. Under this strategy, internal motor commands are manipulated to control the BCI using the same motor cortical circuitry used during natural motor behaviors. Because only a few command variables need to be manipulated to achieve this goal, learning can proceed rapidly and flexibly, and, because the motor cortical circuitry is conserved, the operation of motor cortex during natural motor control is conserved as well.

To study the neural and behavioral consequences of this learning strategy, we formulated a mechanistic model of re-aiming in which the internal command variables specify upstream inputs to motor cortex. By analyzing how these inputs get transformed into motor cortical activity patterns through the circuit’s nonlinear recurrent dynamics, we were able to demonstrate that re-aiming can in fact account for a wide range of experimental observations about BCI learning. This model can explain the different timescales of learning required for different BCI decoders,^14,16^ selective changes in motor cortical tuning curves over learning,^11,12,61^ and the seemingly astonishing ability of mammals to flexibly modulate the activity of single neurons via operant conditioning.^28,63,65^ The model also makes a novel experimental prediction about behavioral biases during short-term learning (fig. 4), which we were able to corroborate in previously published data.^14^ The success of this model at replicating these empirical phenomena provides an explanation in terms of the biological dynamics of neural circuits.

### 3.1 Intrinsic variable learning vs. individual neuron learning

An important debate in the BCI learning literature has been whether human and non-human primates are able to precisely learn and control the contribution of individual neurons to a given BCI decoder readout – the so-called “individual neuron learning” hypothesis. Several studies have been directly aimed at testing this hypothesis, leading to evidence in favor^11,12,33,61^ and against it.^34,35^ The alternative hypothesis is often referred to as “intrinsic variable learning”,^9^ whereby subjects learn to control the BCI using the same constrained set of activity patterns usually used for natural motor control, unable to independently control the activity of single neurons. Our model of re-aiming is a particular formalization of this latter hypothesis, with the command variables 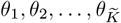 acting as the so-called intrinsic variables.

Our simulations of generalized re-aiming show that many experimental results traditionally attributed to some form of individual neuron learning^17,67^ can be accounted for by intrinsic variable learning. In particular, even classical single neuron operant conditioning results can be reproduced by our model. Our simulations show that the dynamics of recurrently connected neural circuits are capable of generating the activity patterns required by these BCI tasks, without the need to optimize parameters specific to individual neurons or synapses.^68^ This suggests caution in underestimating the role of macroscopic cognitive strategies^69,70^ when observing what may look like highly specific, microscopic, changes to the activity of single neurons.

### 3.2 The role of synaptic plasticity in BCI learning

All the results we replicated here have been previously replicated by various models of synaptic plasticity within motor cortex. As argued in the introduction, however, learning by optimizing synaptic parameters entails solving an extremely high-dimensional optimization problem with no access to explicit gradients, which would limit learning to be slow and brittle. Several of these previously proposed models worked around this problem by using small and simplified feed-forward models of motor cortex^17,68^ or biologically implausible learning rules.^19,20,71^ A few have demonstrated that, for simple tasks like operant conditioning, biologically plausible learning rules can in fact succeed in biologically relevant regimes despite these obstacles.^18,67^ However, to our knowledge, none have comprehensively accounted for all three sets of experimental results considered here, including the observed effects on non-recorded neurons and the dependence of operant conditioning performance on neural correlations.

That said, the present study does not by any means rule out the possibility that synaptic plasticity within motor cortex may play an important role in BCI learning; rather, it reveals the surprising capabilities of a pure re-aiming strategy. The true mechanisms underlying BCI learning most likely comprise a mixture of both re-aiming as well as synaptic plasticity, and future work will be needed to tease apart the contributions of these two learning mechanisms and understand how they are coordinated.

One natural possibility is that synaptic plasticity operates on a much slower timescale than reaiming.^72^ This could help explain selective changes to motor cortical responses that only arise late in learning and are not replicated by our model. For example, Clancy et al. (2014) observed in their operant conditioning experiments that indirect neurons not strongly correlated with the target neuron become silent late in learning.^65^ Ganguly et al. (2011) similarly observed that indirect neurons become less tuned to reach direction after days of practice with a given BCI decoder. Jarosiewicz et al. (2008) reported a similar effect in rotated neurons after a credit assignment rotation perturbation (although note that this effect seems to disappear when increasing the proportion of neurons rotated, see [12, 61]). These selective changes in tuning strength are not reproduced by our model of re-aiming (Supplementary Figure S5a), but previous theoretical work has demonstrated that they can be reproduced by reward-modulated Hebbian plasticity in a simplified model of motor cortex.^17^

An additional set of observations that are not well accounted for by our model come from a few recent studies demonstrating long-term changes in motor cortical activity after short-term learning.^73,74^ In particular, Losey et al. (2024) found that motor cortical activity during baseline decoder control changed before and after learning a WMP within a single experimetal session. Our model could account for this if the upstream population driving motor cortex encoded not only the motor commands relevant for control but also additional variables indexing the current behavioral context,^73,75^ or a memory trace of the current task.^74^

### 3.3 The “intrinsic manifold” of population activity

A simple but important takeaway from this study is that the low-dimensional structure of activity in a population depends not only on the intrinsic dynamics and connectivity within that population, but also on the structure of its upstream input. The observation that population activity is confined to a low-dimensional subspace – often termed the “intrinsic manifold”^14,19,76^ or the “neural modes”^58^ – does not mean that the circuit connectivity prevents it from generating activity patterns outside of this subspace. It is likely that many more activity patterns outside of this subspace are accessible, but that only a low-dimensional subset are accessed by the inputs evoked by the subjects’ behavior during the recording session.^60,77^

This insight leads to a novel interpretation of the observation that outside-manifold perturbations require a longer time to learn than their within-manifold counterparts.^14,16^ Previous models of this phenomenon have assumed that the longer learning time reflects the challenge of modifying the motor cortical connectivity to permit the production of activity patterns outside of the intrinsic manifold.^19,20,68,71^ Our simulations demonstrate that this isn’t necessary, and that in many cases it may suffice to simply exploit additional input dimensions beyond those evoked by the calibration task (fig. 5a, Supplementary Figure S1e). Under this model of learning, the longer learning time required for OMPs reflects the fact that these new input dimensions need to be discovered from scratch, as the calibration task provides little prior information about them.

Another important aspect of the intrinsic manifold that this study highlights is its nonlinear structure. Because firing rates are bounded from below by 0, activity patterns are confined to the positive orthant of state space. This constraint imposes a conical structure on population activity within the intrinsic manifold (fig. 3f, Supplementary Figure S2), which we show in Supplementary Materials Section S.1.3 is in fact necessary to account for experimentally observed behavioral biases in WMP learning. Given the strong behavioral repercussions this structure can have on BCI control, understanding and identifying such nonlinear structure in motor cortical activity may prove crucial both for understanding BCI learning as well as for designing better BCI decoders.

### 3.4 The role of the calibration task in BCI learning

From a more practical perspective, our theory of re-aiming highlights the role of the calibration task in BCI learning. The calibration task is typically seen as a way to calibrate the decoding parameters; that is, as a source of information for constructing the BCI *decoder*. Here we suggest that it additionally serves as a source of information for the subject itself, that is, for the BCI *learner*.^78,79^ For example, in modeling WMP learning, we assumed that subjects re-aimed with the two command variables modulated by the calibration task; in modeling operant conditioning, we assumed that subjects re-aimed with the same command variables driving spontaneous behavior prior to the operant conditioning task. If any other two command variables had been optimized instead, the re-aiming strategy would not have succeeded in solving the task. It is the prior information provided by the calibration task that allows efficient learning by telling the subject which command variables to re-aim with. This hypothesis is consistent with various BCI learning studies demonstrating that subjects learn to control BCIs using the same patterns of activity evoked by the task they were engaged in just prior to BCI learning.^18,29^

Importantly, it predicts that the calibration task can influence subjects’ ability to learn a given BCI decoder, and therefore that careful design of this task could help improve subjects’ learning speed. For example, the calibration task should evoke changes in as few command variables as necessary, so that subjects subsequently re-aim by optimizing only those 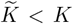 command variables and avoid wasting time exploring modifications to other command variables. A prediction of our model is that learning should be slower when the calibration task evokes changes in more command variables.

### 3.5 How are the re-aiming solutions learned?

The theory presented here treats the question of *what* solutions subjects learn, and makes no claims about *how* they are learned and subsequently maintained. That said, a strong assumption we made in motivating the re-aiming learning strategy was that learning could operate within the low-dimensional space of the command variables. It is this low dimensionality that we claimed would be critical for efficient learning; if the command variables were learned by simply optimizing the connectivity of an upstream circuit, then the limitations of learning by synaptic plasticity would also apply to learning by re-aiming.

One intriguing resolution to this problem would be that the command variables are stored and updated in the *activity* – rather than the *synapses* – of an upstream circuit, as in the pre-frontal cortex model of Wang et al. (2018).^80^ In this model, a recurrent neural network implicitly stores a behavioral policy in its internal state, which, through the network’s dynamics, is updated over time as it interacts with the environment and observes which actions are rewarded in which states. A similar architecture might operate upstream of motor cortex, whereby an upstream circuit continually stores and updates a sensorimotor policy for selecting low-dimensional motor commands. This learning circuit would likely encompass additional populations beyond those directly driving motor cortex, such as the basal ganglia, which are well known to play an important role in BCI learning.^64,81–83^

We finally remark that the short timescale of WMP learning closely mirrors that of motor adaptation, in which subjects adapt their natural movements to a systematic environmental perturbation. Learning these tasks typically requires 100’s of trials of practice,^2,6,7,73^ similar to the time it takes non-human primates to learn WMPs. Neural recordings during these tasks have suggested that changes in neural activity during motor adaptation are driven by changes in the preparatory input from dorsal pre-motor cortex to primary motor cortex.^7,84^ Moreover, changes in the preparatory state of motor cortex (presumably set by upstream inputs^43,44,46^) have been shown to play a critical role in motor adaptation tasks under manual control^73^ as well as BCI control.^32^ These results are consistent with the idea that, much like in our model of re-aiming, motor adaptation involves modifications to the inputs driving motor cortex while motor cortex itself remains unchanged. Our model of re-aiming may thus be relevant to more general and naturalistic forms of sensorimotor learning beyond BCIs.

## Methods

### 4.1 Motor cortical dynamics

Motor cortical activity was simulated by integrating equation 1 using a standard 4th order Runge-Kutta method with step size 0.1ms, implemented with the torchdiffeq Python package.^85^ Reachable activity patterns, **r**(*t*_end_; ***θ***), were computed by integrating this equation from the initial condition at time *t* = 0 to the endpoint time *t* = *t*_end_, with constant inputs determined by the given motor command, ***θ***, using equation 2. We assumed silent initial conditions, *x*_*i*_(0) = 0, to enable a computationally efficient solution to the re-aiming objective function (see equation 12). Decoding from the reachable activity patterns using equation 3 yields the reachable readouts, **y**(*t*_end_; ***θ***).

In all simulations in the main text, sparse random recurrent weights, 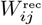 were used: only 10% of these weights were set to non-zero values, which were independently sampled from a zero-mean Gaussian, 𝒩 (0, 1*/N*), where *N* is the total number of motor cortical neurons in the network. Input weights were all sampled from a zero-mean Gaussian, 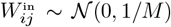, where *M* is the total number of inputs. Encoding weights were sampled randomly from the standard Gaussian distribution, *U*_*ij*_ ∼ 𝒩 (0, 1) (any normalization is taken care of by the metabolic cost term in equation 4 when computing re-aiming solutions). Other connectivity patterns are considered in Supplementary Figure S1. We used *τ* = 200ms, as in the motor cortical model of Hennequin et al. (2014). To enable efficient numerical simulation, network size was set to *N* = *M* = 256. Simulations with larger networks (up to *N* = *M* = 2048 neurons) produced similar results (data not shown).

### 4.2 Computing re-aiming solutions

Concatenating the command variables into a 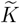 -dimensional vector that contains only the command variables being optimized, 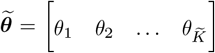,we can treat equation 4 as an optimization over all 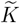 -dimensional vectors 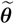 in 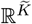.We can simplify this optimization problem by first analytically solving for the optimal magnitude of 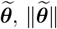 given its direction. Once we have this optimal magnitude, all that remains is an optimization over its direction – an optimization over unit vectors on the 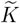-dimensional unit hypersphere. This is a 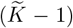-dimensional manifold that, importantly, is bounded, so we can hope to find the optimal direction efficiently by brute-force search, avoiding the difficulties of non-convex gradient-based optimization in high dimensions.

Formally, we decompose the re-aiming optimization (equation 4) into an optimization of the norm, *s*, and direction, 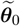,of 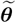,

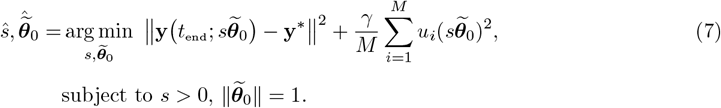

We can analytically solve for the optimal magnitude, ŝ, by exploiting two simplifications afforded to us by the rectified linear activation function *ϕ*(·) of the motor cortical dynamics (equation 1b). The first is the scale-invariance of the activation function (*ϕ*(*sx*) = *sϕ*(*x*) for any *s* ≥ 0), which accordingly endows the motor cortical dynamics with scale invariance,

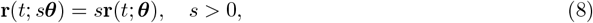

whenever *x*_*i*_(0) = 0 (see Supplementary Materials Section S.2.1 for a formal proof), which we assumed to be the case in our simulations. The second simplification is to approximate the quadratic cost term by its large *M* limit

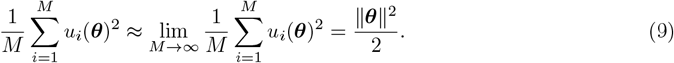

The equality holds whenever the encoding weights (*U*_*ij*_ in equation 2) are independent and identically distributed with zero mean and unit variance, as they are here, so that the law of large numbers can be invoked to replace the sum with an expectation over the encoding weight distribution (the factor of 1*/*2 arises from the fact that only half of each axis counts towards the expectation due to the linear rectification, see Supplementary Materials Section S.2.2 for a formal proof). Inserting these two equations into equation 7 together with the BCI readout equation 3, we obtain

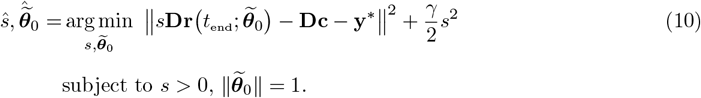

It is then straightforward to solve for ŝ in terms of 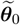,yielding the following closed set of equations:

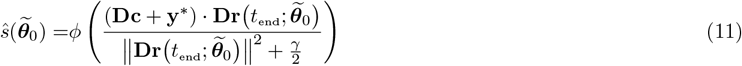

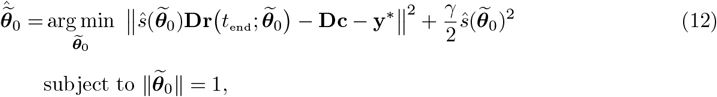

where the ·notation denotes the Euclidean dot product. We have thus reduced what was an optimization over all vectors in 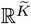 (equation 4) to an optimization over all vectors living on the 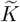-dimensional unit hypersphere (equation 12).

We can therefore approximately solve this new optimization problem via brute-force search, by simply uniformly sampling a large number of 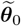’s on the unit hypersphere and identifying the one that produces the smallest value of the loss function in equation 12. Evaluating 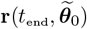 for a large number of 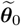’s can be done efficiently by using a GPU to integrate in parallel the dynamics driven by each 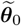,Note that, once these activity patterns have been calculated, they can be re-used to perform the brute-force search optimization for any given value of *γ*, without having to again integrate the dynamics.

For simulations with 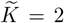, this brute-force search algorithm sufficed to produce good re-aiming solutions. In this case, the relevant hypersphere is the unit circle, from which it is straightforward to sample densely and uniformly. For simulations of generalized re-aiming, however, we took an additional step to ensure the obtained solutions were as good as they could be even for the larger values of 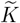, where it becomes more difficult to sample densely from the corresponding unit hypersphere. We first performed a brute-force search over 2^17^ vectors sampled uniformly from the unit hypersphere, as just described, to obtain an initial estimate of the re-aiming solution. This initial estimate was then used as a starting point for the L-BFGS algorithm,^86^ which we then applied to minimize the re-aiming loss function (equation 4) with respect to the raw command variables 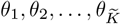. We found that this additional step was essential when 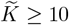.

In all simulations in the main text, we used an endpoint time of *t*_end_ = 1000 ms, reflecting the typical time it takes for trained primates to complete center-out reaches under BCI control.^11,14^ The results of simulations with other endpoint times are shown in Supplementary Figure S1b. To simulate a center-out reaching task, the target readouts **y**^*^ were set to eight equally spaced unit vectors on the unit circle (cf. fig. 3c). Mean squared error is calculated as

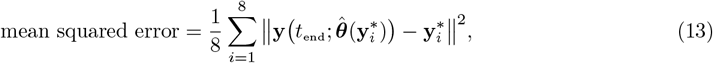

where 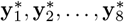 correspond to the eight radial target readouts. Because the targets are unit norm, a mean squared error of 1.0 corresponds to that achieved by readouts at the origin.

In the case of operant conditioning, there is no “target readout” per se, as subjects are simply instructed to modulate firing rates as much as possible in a given direction. In this case, a different re-aiming objective was used, see the section “Simulation of operant conditioning” for details.

### 4.3 Setting the metabolic cost weight

The metabolic cost weight parameter *γ* was picked to ensure that low mean squared error would be achieved under the baseline decoder with 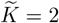.We calculated re-aiming solutions with 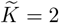 for the baseline decoder under a wide range of values of *γ*. We took advantage of the fact that the brute-force search algorithm outlined above allows us to easily evaluate solutions for different values of *γ* with only a single forward pass of the model. Once we had re-aiming solutions for each value of *γ*, we calculated the error achieved by these re-aiming solutions for each target readout, and found the largest value of *γ* that permitted a squared error of less than .05 for each of the eight targets. *γ* was then fixed to this value for simulations with all the decoder perturbations.

### 4.4 Characterizing the reachable manifold

The reachable manifold is the set of activity patterns at time *t*_end_ that can be generated by any motor command allowable under the re-aiming strategy. We assume that these allowable motor commands are bounded, reflecting the fact that (i) actual extrinsic motor variables are finite and bounded and (ii) upstream firing rates are bounded. Formally, we enforce this by assuming an upper bound on the motor command norm, ∥***θ***∥ ≤ *s*_max_. In our simulations of short-term learning of WMP’s and OMP’s, we set the value of this bound to the maximum norm over all 2D re-aiming solutions to all decoders. Specifically, we computed re-aiming solutions for each target readout and decoder perturbation (8 target readouts × (100 WMP’s + 100 OMP’s + baseline decoder) = 1,608 re-aiming solutions) with 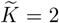,calculated their norms, and set *s*_max_ to their maximum. For the randomly connected network presented in the main text, we found this value to be approximately 1.25.

In figure 3f, we drove the motor cortical network with motor command vectors with five distinct norms, ∥***θ***∥ ∈ {0.1, 0.4, 0.7, 1.0, *s*_max_}, chosen to aid visualization of the reachable manifold. We picked 256 equally spaced angles between 0 and 2*π* and constructed 2D vectors with each angle and each norm to obtain the command variable pairs, *θ*_1_, *θ*_2_, shown in fig. 3e (all other command variables were set to zero). We then simulated the motor cortical network with each of these motor commands to obtain a large ensemble of activity patterns on the reachable manifold, **r**(*t*_end_; ***θ***), and projected these onto their top three principal components to obtain the visualization in figure 3f. Figure 3c plots the readouts of each of these activity patterns from three different decoders. Figure 4a plots the readouts from four different WMP decoders, in this case using activity patterns generated from motor commands with four different norms equally spaced between 0.1 and *s*_max_ (thus producing four loops of readouts instead of five). In figure 7a, five motor command norms equally spaced between .1 and the maximum norm of the re-aiming solutions for that pair of neurons were used. These choices were all made to aid visualization of the reachable manifold’s structure.

To obtain the centroid, 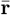,and covariance, **Σ**_*r*_, of the reachable manifold for 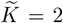, we computed expectations over a uniform distribution on the reachable manifold. Note that an expectation over uniformly distributed activity patterns on the reachable manifold is not the same as an expectation over activity patterns generated by uniformly distributed command variables. These two distributions of activity patterns are related via the Jacobian of the mapping from command variables, *θ*_1_, *θ*_2_, to activity patterns, **r**(*t*_end_; ***θ***), which was used to derive the following two expressions for 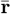 and **Σ**_*r*_ (see Supplementary Materials Section S.2.3 for full mathematical derivation):

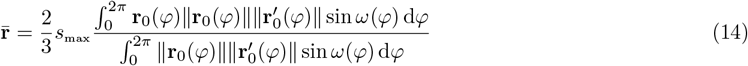

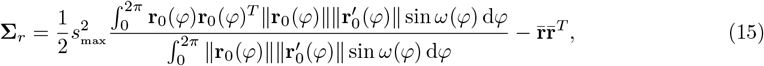

where **r**_0_(*φ*) is the population activity at time *t*_end_ generated by a pair of non-zero command variables *θ*_1_, *θ*_2_ with angle *φ* and unit norm, 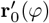 is its derivative with respect to *φ*, and *ω*(*φ*) is the angle between **r**_0_(*φ*) and 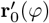. We used a finite-differences approximation for the derivative 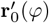 and computed these integrals numerically by summing over a dense range of values of *φ* ∈ [0, 2*π*]. This estimate of the reachable manifold centroid, 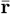,is plotted in figures 4a, 4b, and S2a. This estimate of the reachable manifold covariance, **Σ**_*r*_, is used for the variance explained curve plotted in figure 3g.

Analagous calculations for 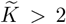 quickly become numerically intractable, as the derivatives and integrals become multivariate as the number of polar coordinates increases. We thus chose to characterize the dimensionality of the reachable manifold under *generalized* re-aiming by the covariance over activity patterns produced by uniformly distributed motor commands, which we denote by **Σ**_*θ*_. As already noted, this is not the same as the covariance over activity patterns uniformly distributed on the reachable manifold manifold, but these two covariances are strongly related. This covariance is given by (see Supplementary Materials Section S.2.4 for derivation)

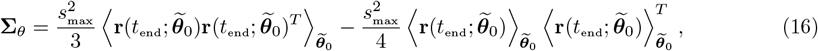

where 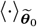 denotes an expectation over a uniform distribution on the unit-norm 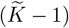-sphere. These expectations were estimated numerically by uniformly sampling 2^17^ vectors 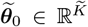 from the unit hypersphere, setting the 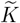 non-zero command variables to these values, calculating the activity patterns generated by these motor commands at time *t*_end_ = 1000ms, 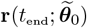,and then averaging over the resulting ensemble of activity patterns.

This estimate of the reachable manifold covariance, **Σ**_*θ*_, is used to compute the participation ratio plotted in figure 5b as a function of 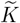,using the formula

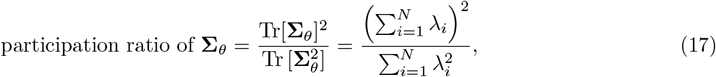

where *λ*_1_, *λ*_2_, …, *λ*_*N*_ are the eigenvalues of **Σ**_*θ*_.

### 4.5 Quantifying biases in BCI readouts

To quantify behavioral biases, we used the maximal cursor progress metric defined in equation 6. This equation was solved by again exploiting the same re-parameterization of the motor commands we used for calculating re-aiming solutions (equation 7). Specifically, we decompose the vector of non-zero command variables, 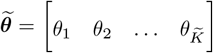,into a magnitude and direction, 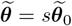 where *s >* 0 and 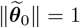. Plugging in the readout equation (equation 3) into the definition of cursor progress (equation 5) and exploiting the scale invariance of the motor cortical dynamics (equation 8), we have that the maximal cursor progress is given by

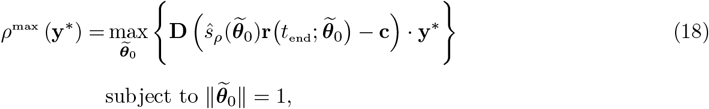

where

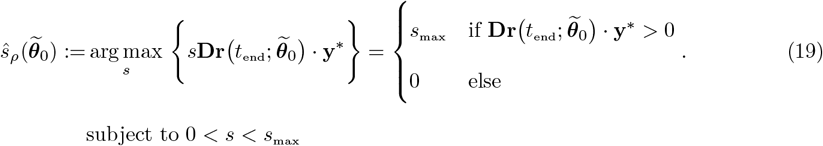

Since 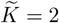 in these simulations (and thus 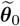 is just a 2D unit vector), we were able to effectively solve the optimization problem in equation 18 by brute-force search over densely and uniformly sampled 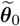’s from the unit circle.

### 4.6 Re-analysis of data from Sadtler et al. (2014)

To quantify behavioral biases in the experimental data, we estimated the maximal cursor progress in each experimental session by using the cursor progress values observed in the window of 50 contiguous trials of WMP control with shortest average reach completion times. At each timestep in each trial, we calculated the cursor progress in the direction of the target relative to the cursor’s position at that time. We then binned the per-timestep relative target directions into 45^*o*^ bins centered at the eight radial reach target angles, and averaged the cursor progresses in each bin to obtain an average cursor progress for each target direction. We take these averages to be estimates of the subject’s *maximal* cursor progress with that session’s WMP decoder, as they are taken from the 50 trials with fastest reaches.

To predict the maximal cursor progress in each session from the target direction angle with 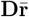,we sought an estimate of the reachable manifold centroid, 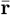,that could be empirically measured from the recorded neural activity, without access to the underlying reachable manifold. To do this, we noticed that, in our model, the time- and trial-averaged population activity generated by the re-aiming solutions for the baseline decoder – which we denote by 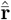– was highly correlated with the true reachable manifold centroid, 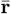.We therefore used 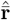 to estimate 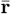,since 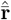 can be easily estimated in the experimental data by simply averaging motor cortical activity during the baseline decoder control block in each session. We calculated target-specific means by averaging motor cortical activity over all trials and time during reaches to each target, and then averaged these target-specific means over targets to obtain 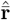.This was the estimate of the reachable manifold centroid – and its projection through the respective WMP decoder in each session, 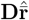-used in the analysis presented in figure 4d.

To keep the analysis of the data and the model consistent, we also used an analogous estimate of the reachable manifold centroid for the analysis of the model in figure 4c. In this case, 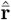 was measured by simulating reaches to each target by driving the motor cortical network dynamics with the re-aiming solutions for the baseline decoder, and then averaging the motor cortical firing rates over all time and over all eight reach directions. We found that the resulting negative correlation was similar regardless of whether the true reachable manifold centroid, 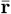,(data not shown) or its estimate, 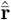,was used.

### 4.7 Simulation of the calibration task

The calibration task was simulated by setting the first two command variables *θ*_1_, *θ*_2_ to the coordinates of the reach direction 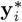 being presented in each trial (a 2D unit vector pointing in one of eight equally spaced angles), and setting all other command variables to zero (*θ*_3_ = *θ*_4_ = … = *θ*_*K*_ = 0). To simulate noise in the neural responses, we added noise in the dynamics, in the motor commands, and in the initial conditions on each trial. At each timestep, zero mean Gaussian noise with standard deviation 0.05 was sampled and added to the single neuron potentials *x*_*i*_(*t*) and to the two command variables *θ*_1_, *θ*_2_. Initial conditions in each trial were sampled randomly from a 0-mean isotropic Gaussian with standard deviation 0.1. The network was driven for 1000ms in each trial, matching the duration of each trial in the experiment.

For simulations with WMPs and OMPs, we simulated 10 trials of each reach direction, replicating the structure of the calibration task used by Sadtler et al. (2014). For simulations with credit assignment rotation perturbations (fig. 6), the calibration task was identical except that only a single trial of each reach direction was simulated, to mimic the decoder initialization procedure of Zhou et al. (2019). Note that in all cases re-aiming with 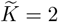 command variables implies re-aiming with the same two command variables driving the calibration task neural responses, *θ*_1_ and *θ*_2_.

### 4.8 Within- and outside-manifold perturbations

In the BCI system used by Sadtler et al. (2014) and Oby et al. (2019), 96-channel microelectrode arrays were used to record neural activity. Spikes were detected by threshold crossings in the recorded voltage signals at each electrode, resulting in a series of spike trains at each electrode. Spike trains at each electrode could therefore contain spikes from multiple neurons near the electrode site, as no spike sorting was performed. In total, about 100 neurons were likely to have been recorded, constituting a small fraction of the total population of neurons in motor cortex. To simulate this, we ensured that the BCI decoder in our simulations only had access to a linear mixture of firing rates from *N*_*r*_ = 99 neurons (so as to be divisible by *ℓ* + 1 = 9, to group neurons by modulation depth, see below). This was done by first multiplying the firing rates with a *N*_*r*_ × *N* “recording matrix”, **H**, which had the following tri-diagonal structure

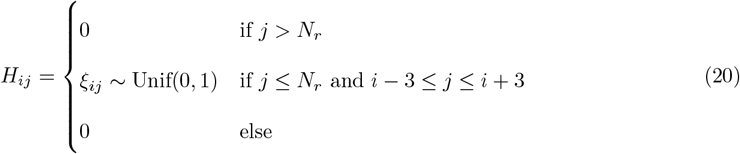

Thus, each “neural unit” in the vector **Hr** is composed of a linear mixture of seven neurons, with “neural units” with adjacent indices mixing together overlapping sets of neurons.

Following Sadtler et al., we next z-scored the activity recorded by each neural unit with respect to its statistics during the calibration task,

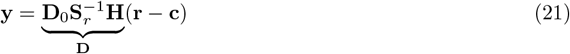

where **c** is an *N* -dimensional vector with the mean firing rate of each neuron and **S**_*r*_ is an *N*_*r*_ × *N*_*r*_ diagonal matrix with the standard deviation of each “neural unit”, measured from the simulated activity during the calibration task. Readouts, **y**, were obtained by decoding from the *N*_*r*_-dimensional vectors of z-scored mixed firing rates. It is the 2 × *N*_*r*_ *effective* decoding matrix, **D**_0_, that is perturbed by the WMP and OMP decoder perturbations (see below). Note that the full 2 × *N* decoding matrix, **D**, is such that only its first *N*_*r*_ columns are non-zero, reflecting the fact that only a subset of the full population of motor cortical neurons is recorded by the BCI.

For the baseline decoder, the effective decoding matrix **D**_0_ was constructed following the methods of Sadtler et al., with the exception that we used Principal Components Analysis instead of Factor Analysis to estimate the intrinsic manifold. This choice was made purely for the sake of numerical convenience, as Principal Components Analysis has a closed-form solution that can be computed more efficiently. The full procedure for estimating the intrinsic manifold and constructing the baseline decoder is outlined in detail in Supplementary Materials Section S.3. In short, the baseline decoder effective decoding matrix can be expressed as a product of a 2 × *ℓ* matrix **K** and an *ℓ* × *N*_*r*_ matrix **L**, where *ℓ* is the dimensionality of the intrinsic manifold,

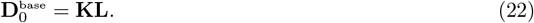

The matrix **L** projects *N*_*r*_-dimensional activity patterns down to the *ℓ*-dimensional intrinsic manifold; its rows span the subspace defined by the intrinsic manifold (Supplementary Materials Section S.3.2, equation 71). In our simulations we used *ℓ* = 8, as we found that the top 8 principal components contained 95% of the variance in the simulated calibration task responses. The matrix **K** then translates the resulting *ℓ*-dimensional dimensionality-reduced activity patterns into 2-dimensional BCI readouts. This matrix is fit to the calibration task data, by fitting a Kalman filter that accurately decodes the calibration task stimuli from the dimensionality-reduced calibration task neural responses at each timestep and trial (Supplementary Materials Section S.3.2, equation 72).

Within-manifold perturbations (WMPs) perturb the baseline decoder in such a way that the row space of **L** remains intact, so as to conserve the decoder’s relationship with the intrinsic manifold. This is done by simply shuffling the rows of **L** without modifying them, via pre-multiplication with a random *ℓ* × *ℓ* permutation matrix **P**,

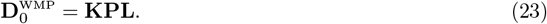

Outside-manifold decoders, on the other hand, directly disrupt the row space of **L**. This is done by randomly shuffling the components of each of its rows, via post-multiplication with a random *N*_*r*_ × *N*_*r*_ permutation matrix **P**,

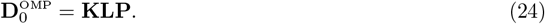

It is important to keep in mind that both WMPs and OMPs can change the readouts in complex ways, beyond a simple rotation like that depicted by the cartoon in figure 2b (for examples, see fig. 4a here and supplementary figure 2 in^29^). Once the baseline decoder was constructed, we randomly sampled 100 within-manifold and 100 outside-manifold perturbations by randomly selecting 100 *ℓ* × *ℓ* and *N*_*r*_ × *N*_*r*_ permutation matrices, respectively.

To minimize any differences between these two types of decoder perturbations that would go beyond their opposing relationship to the intrinsic manifold, we imposed several restrictions on the selected permutation matrices, as was done by Sadtler et al. (see Supplementary Materials Section S.3.3 for details). First, we enforced that the mean principal angle between the row space of the baseline effective decoding matrix and the row space of each perturbed effective decoding matrix was between 60^*o*^ and 80^*o*^. Second, we enforced that population activity produced by the re-aiming solutions for the baseline decoder would produce readouts through each perturbed decoder that resulted in a mean squared error between 0.6 and 0.8. Finally, we fit tuning curves to the neural activity generated by the re-aiming solutions for the baseline decoder, and asked how much the preferred directions would need to change to produce the same readouts under the perturbed decoder, following the same procedure employed by Sadtler et al. We enforced that this change be between 30^*o*^ and 45^*o*^. We typically found that about 100-200 permutations out of all possible permutation matrices satisfied these criteria. We then randomly sampled 100 of them.

Following the procedure used by Sadtler et al. with monkey L, we did not consider all possible *N*_*r*_ × *N*_*r*_ permutation matrices for OMPs (as there are 99! of them). Rather, we grouped all *N*_*r*_ neural units into *ℓ* groups, and then considered all *ℓ*-dimensional permutations of these groups (of which there are 8!). In other words, rather than permuting all *N*_*r*_ columns of **L**, *ℓ* groups of columns were permuted. This ensured that the total number of possible decoder perturbations was the same for WMPs and OMPs. The *ℓ* groups were assigned as follows: for each neural unit in **Hr**, we fit a cosine tuning curve to its calibration task responses to obtain its modulation depth (equation 26), assigned the *N*_*r*_*/*(*ℓ* + 1) neurons with smallest modulation depths to a small-modulation group not to be permuted, and randomly assigned the remaining neurons to *ℓ* high-modulation groups to be permuted; following Sadtler et al., the small-modulation group was never permuted to avoid cases in which an inactive or noisy neuron could get assigned a large decoding weight.

In figure 3g, we define the “dimensions” of the intrinsic manifold as a set of orthonormal basis vectors **f**_1_, **f**_2_, …, **f**_*ℓ*_ spanning the intrinsic manifold (Supplementary Materials Section S.3.1). We then calculated the variance explained by each dimension by

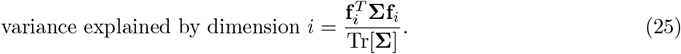

For the gray curve, **Σ** was set to the sample covariance of the simulated calibration task responses. For the purple curve, **Σ** was set to the reachable manifold covariance **Σ**_*r*_ (defined in equation 15). In each case, the cumulative variance explained was calculated by ordering the dimensions by variance explained and then summing them in that order.

### 4.9 Credit assignment rotation perturbations

In the BCI system used by Zhou et al. (2019), recorded activity was sorted by matching spike waveforms to identify spikes from single neurons, resulting in the identification of 10–12 individual neurons. Importantly, each neuron had reliable tuning to reach direction during the calibration task. In our simulation, we modelled this by randomly choosing 80 neurons, fitting tuning curves to their activity during the calibration task, and selecting only those with modulation depth greater than 0.5 (see below for how modulation depth is measured). This typically resulted in 10–15 neurons being included in the BCI decoder (i.e. being assigned non-zero weights in the decoding matrix **D**); no “recording matrix” **H** was used in these simulations. For the particular network model used in the simulations reported in the main text, this selection procedure resulted in *N*_*r*_ = 11 neurons being included.

Tuning curves were fit to time-averaged simulated firing rates in the calibration task, 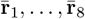,in response to presentation of each of the eight radial reach targets, 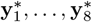 (cf. section “Simulation of the calibration task”). An *N* × 3 matrix of tuning weights, **T**, was fit to predict these average responses from the respective reach target coordinates,

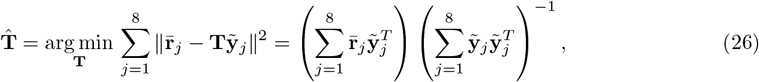

where 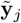 is a 3D vector with the coordinates of the direction of the *j*th reach target, 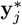,as its first two components and a constant 1 in its third component, included to model baseline tonic firing rates of each neuron. Thus, only the first two columns of the tuning weight matrix **T** model how the *i*th neuron’s firing rate depends on the reach target’s direction, whereas its third column models activity independent of the reach target. To extract from these weights the directional tuning of neuron *i*, we take the 2D vector comprising the first two components of the *i*th row of 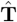. We notate this 2D vector by *m*_*i*_**p**_*i*_, where **p**_*i*_ is a unit vector pointing in its direction and *m*_*i*_ is its norm. The angle of **p**_*i*_ is neuron *i*’s preferred direction, and *m*_*i*_ is its modulation depth.

Following the methods of Zhou et al. (2019), the baseline decoder was constructed as follows. First, raw firing rates were baseline-subtracted and normalized by their modulation depths,

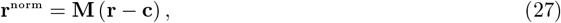

where **c** is given by the third column of 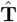, containing the baseline firing rates estimated from the linear regression fit (equation 26), and **M** is an *N*_*r*_ × *N* diagonal rectangular matrix containing the inverse modulation depths *m*^−1^ for each of the *N*_*r*_ neurons recorded by the BCI on the diagonal across the first *N*_*r*_ columns 0 everywhere else. These normalized firing rates were then transformed into 2D readouts by a 2 × *N*_*r*_ effective decoding matrix 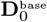 containing the preferred direction vectors **p**_*i*_ of each of the *N*_*r*_ recorded neurons in their corresponding columns. More precisely, the *i*th column of 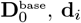,is given by

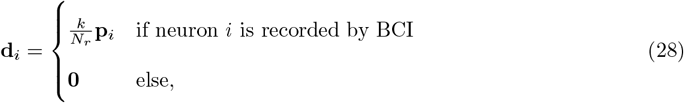

where the scaling constant *k* is chosen to minimize the mean squared error between the readouts from the calibration task activity and the target readouts. This is the classic population vector algorithm (PVA).^87^ The full 2 × *N* decoding matrix of the baseline decoder was thus 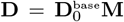,which has non-zero weights only in the *N*_*r*_ columns corresponding to the *N*_*r*_ recorded neurons.

Credit assignment rotation perturbations were constructed by simply picking a random subset of the columns of 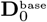 and rotating them. In our simulations, we picked a random 50% of these columns and rotated them 75^*o*^ counter-clockwise, as was done in the decoder perturbations used by Zhou et al. (2019). We sampled 100 random perturbations in this way, in each case rotating a different subset of columns. The normalization matrix **M** and baseline subtraction parameters **c** are kept the same for all decoders.

To measure the tuning changes predicted by re-aiming, we simulated cursor reaches with each decoder by driving the motor cortical network with the re-aiming solutions for that decoder. In each case, noise was applied to the dynamics, exactly as in the calibration task. We then fit tuning curves to each neuron’s time-averaged activity, using linear regression exactly as described in equation 26, and extracted the preferred direction of each rotated, non-rotated, and indirect neuron. For each perturbed decoder, we then determined each neuron’s change in preferred direction by calculating the difference between its preferred direction under the re-aiming solutions for the perturbed decoder and its preferred direction under the re-aiming solutions for the baseline decoder (cf. figures 6b, 6c). These changes were then averaged over all neurons in each sub-population (rotated, non-rotated, or indirect). Figure 6e shows the percentiles (median and quartiles), over all 100 sampled decoder perturbations, of this average tuning change. An analagous analysis of the changes in modulation depth is shown in Supplementary Figure S5a.

### 4.10 Simulation of operant conditioning

In an operant conditioning task, there is no “target readout” per se. The objective is to simply increase the activity of one neuron over another, as much as possible. We can thus express the objective as maximizing the difference in firing rate between the two neurons, which can be thought of as a one-dimensional linear readout from the population. Formally, we calculate readouts in this task by a dot product between the firing rate vector **r** and a decoding vector **d** which has a +1 for the target neuron, a −1 for the distractor neuron, and 0’s everywhere else. This one-dimensional readout indicates how much more active the target neuron is than the distractor neuron. The goal in an operant conditioning task is to maximize this readout.

Adding in a metabolic cost, the objective function we use for re-aiming is

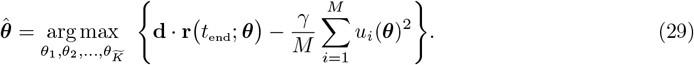

Letting 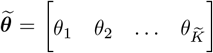 denote the non-zero command variables, we again re-parameterize this optimization problem into an optimization over the magnitude, *s*, and direction, 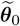,of 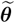,

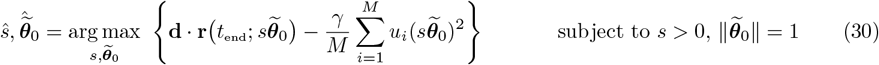

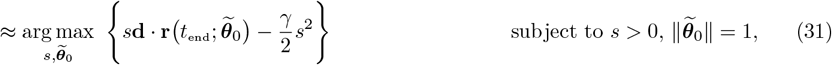

where the approximation follows from the application of the scale invariance of the network dynamics (equation 8) and the mean-field approximation of the metabolic cost (equation 9). This approximation allows us to analytically solve for the optimal magnitude ŝ,

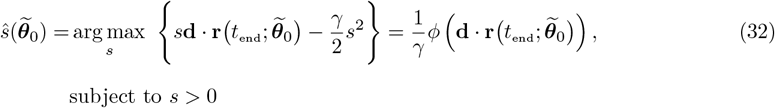

which in turn allows us to solve for the optimal direction, 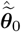,via optimization over the 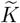-dimensional unit hypersphere,

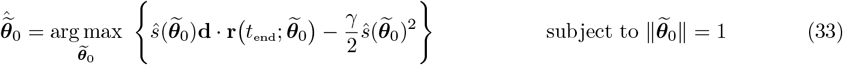

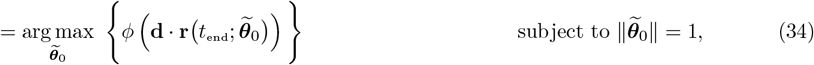

where the second equality follows from plugging in equation 32 and simplifying. In all our operant conditioning simulations, we used 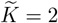,which enabled us to easily solve this optimization problem via brute force search over the unit circle. As in other simulations, we used *t*_end_ = 1000ms.

Note that the optimal readout achieved by the re-aiming solution is

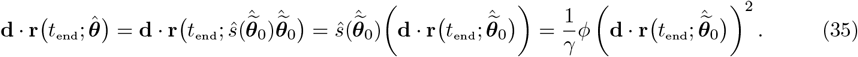

Thus, changing the exact value of *γ* only re-scales the re-aiming solutions and the readouts they produce. We thus simply set it to *γ* = 1 in all our simulations of this task.

In classic operant conditioning experiments,^28^ neurons selected for operant conditioning had to be active prior to the conditioning task to be identified by the recording electrode. We imposed a similar constraint in our simulation by first driving the network with 50 random *K* = 100-dimensional motor commands for *t*_end_ = 1000ms, and identifying the 50% of neurons with highest average firing rate over motor commands and time. The neuron pairs used for operant conditioning were sampled from this sub-population.

To simulate neural activity during a baseline period of spontaneous behavior prior to operant conditioning, we used a similar procedure but now driving the network with 50 random 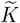-dimensional motor commands, with 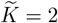.This allowed us to ask whether operant conditioning performance under re-aiming could be predicted from correlations arising during spontaneous behavior driven by the same command variables used subsequently for re-aiming. In figure 7c, correlations were measured by correlation coefficient between the two conditioned neurons. In figure 7d, correlation coefficients between each indirect neuron and the target neuron are plotted against the firing rate of the indirect neuron at *t*_end_ = 1000ms when driving the network with the re-aiming solution.

### 4.11 Table of simulation parameters

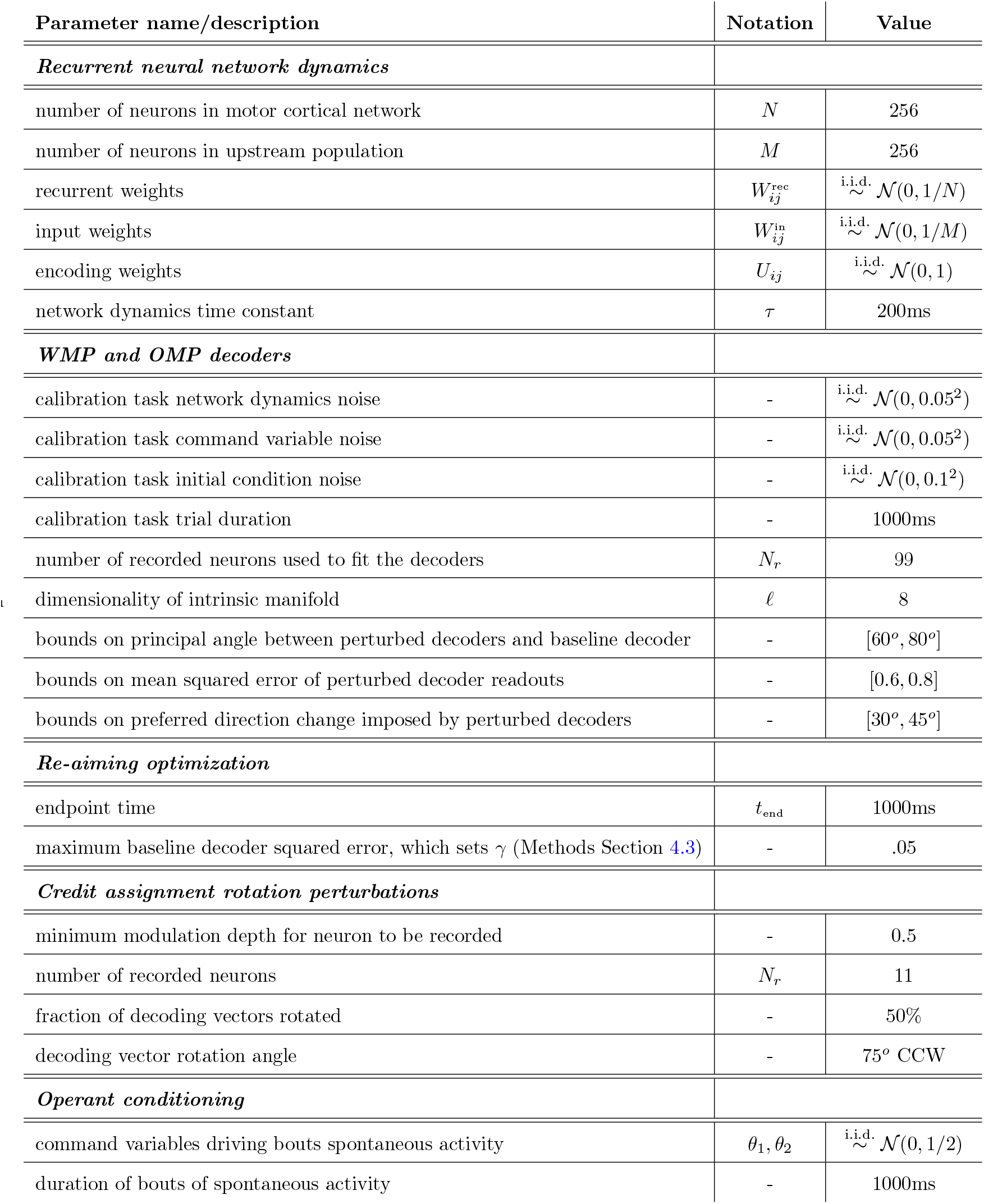

## Acknowledgments

This work was supported by the Gatsby Charitable Foundation (J.A.M., P.E.L.), Wellcome Trust 110114/Z/15/Z (P.E.L.), University College London Research Excellence Scholarship (J.A.M.), NIH K99/R00 MH121533 (M.D.G.), NIH R01 NS129584 (A.P.B., S.M.C., B.M.Y.), NSF NCS DRL 2124066 and 2123911 (B.M.Y., S.M.C., A.P.B.), Simons Foundation 543065 and NC-GB-CULM-00003241-05 (B.M.Y.).

## Supplementary Matrials

### S.1 Extended modeling results

#### S.1.1 Structured motor cortical connectivity

In addition to the randomly connected network architecture used in the results presented in the main text, we simulated re-aiming in the task of Sadtler et al. (2014) with alternative motor cortical connectivity profiles, described below. In each case, we simulated the calibration task, sampled decoder perturbations, and computed re-aiming solutions as described in the main text, except for a few minor modifications noted below. Results of these simulations are shown in figure S1.

##### Random excitatory/inhibitory (E/I) connectivity

a random sparse and balanced E/I recurrent connectivity matrix was constructed following the sampling procedure described in.^1^ In short, all ex1316 citatory weights had the same strength, all inhibitory had the same strength (re-scaled relative to the excitatory weights to account for the different number of excitatory and inhibitory neurons), and each row of the weight matrix was enforced to be 0 mean to enforce so-called E/I balance. We used a sparsity of 10% (i.e. only 10% of weights were non-zero), with 80% of the neurons in the population being excitatory. Input and encoding weights were sampled randomly as for the randomly connected network in the main text.

**Figure S1:**
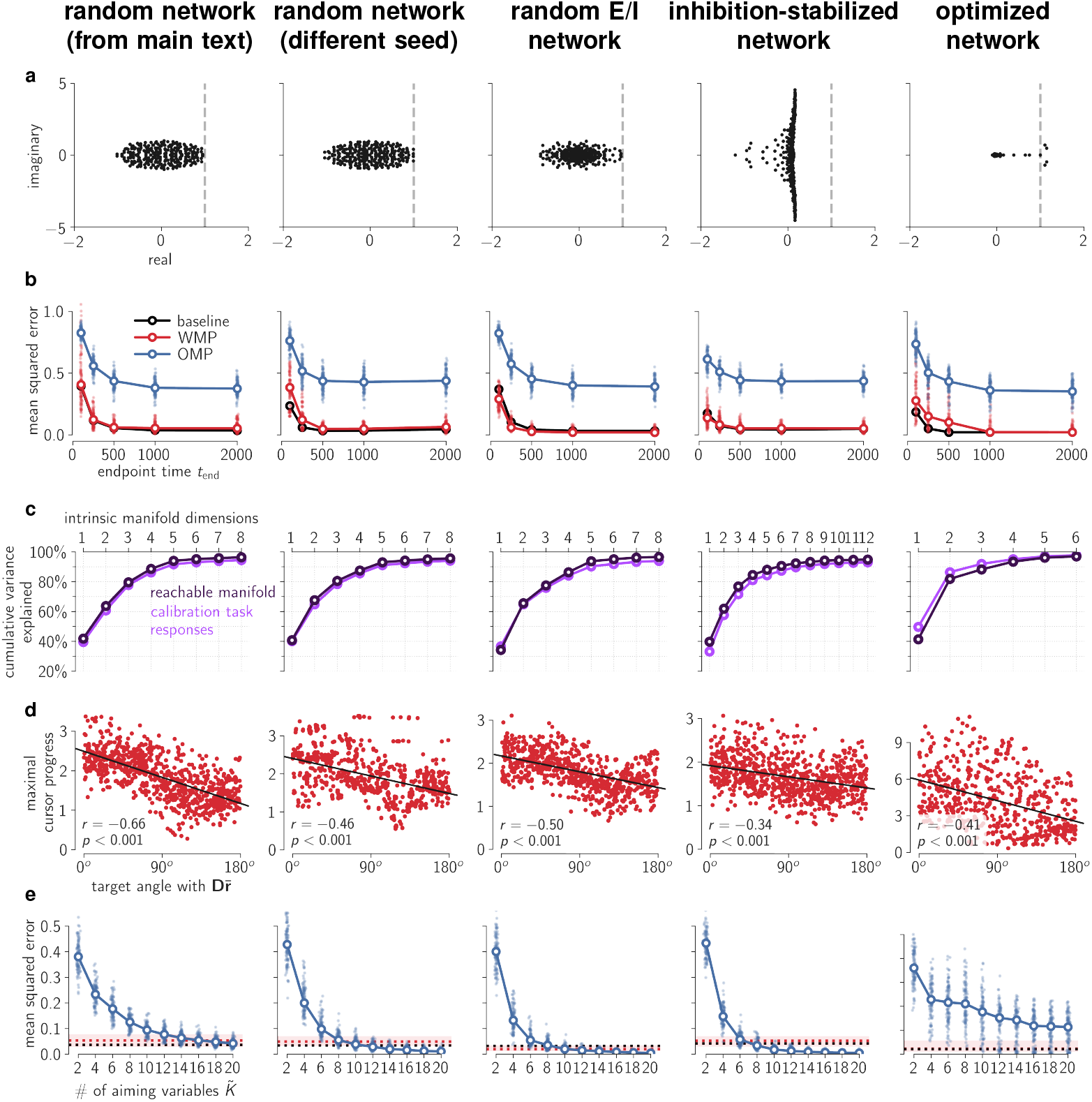
Each column shows simulation results for a different network: (1) the randomly connected network used in the main text; (2) another randomly connected network, with weights sampled in exactly the same way; (3) a network with random E/I connectivity; (4) a network with inhibition-optimized E/I connectivity; (5) network with connectivity optimized for delayed center-out reaching. a. Eigenspectra of recurrent weight matrices of each network, plotted on the complex plane. Note that the optimized network has low-rank connectivity: almost all eigenvalues are clustered at 0. Dashed vertical line marks the linear stability boundary of a real value of 1.0. b. Mean squared error achieved by re-aiming solutions for different endpoint times, *t*_end_, for each decoder. Lighter markers correspond to individual decoders, darker open markers (connected by lines) show medians over all decoders. Note that the metabolic cost weight, *γ*, is fixed to the same value, which was picked to guarantee low error under the baseline decoder for *t*_end_ = 1000ms only (see Methods Section 4.3). Thus, the error rises as the endpoint time decreases below this, as higher magnitude motor commands are necessary to achieve low error faster. However, note that the difference between baseline decoder, WMPs, and OMPs remains the same even at these lower values of *t*_end_. c. Calibration task response and reachable manifold variance cumulatively accounted for by each dimension of the intrinsic manifold, as in fig. 3g. Intrinsic manifold was found to be about 12-dimensional with inhibition-optimized connectivity and about 6-dimensional with connectivity optimized for delayed center-out reaching. d. Maximal cursor progress for each target and WMP as a function of target direction angle with 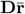,as in fig. 4c. e. Mean squared error achieved for each OMP by re-aiming with different numbers of command variables, 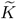, as in fig. 5a.

##### Inhibition-optimized excitatory/inhibitory connectivity

a sparse and balanced E/I recurrent connectivity matrix was constructed following the optimization procedure described in.^2^ In short, the excitatory weights were initialized to be very strong, and then inhibitory weights were optimized to ensure the dynamics were stable (by minimizing the spectral abscissa of the full connectivity matrix). Half of the neurons were assigned to be excitatory, and the inhibitory weights were enforced to be on average three times stronger than the excitatory weights. The only difference with^2^ was that the weight matrices were initialized with a spectral radius of 5, rather than 10. This was necessary as we found that an initial spectral radius of 10 lead to chaotic dynamics under constant input. Input and encoding weights were sampled randomly as for the randomly connected network in the main text.

Because of their highly non-normal dynamics, these networks were highly sensitive to changes in initial conditions, even with the reduced initial spectral radius. We therefore reduced the standard deviation of the initial conditions by half when simulating the calibration task (see Methods Section 4.7). These networks produced much higher-dimensional calibration task responses than the randomly connected network, so a 12-dimensional intrinsic manifold was used for constructing WMP’s and OMP’s (i.e. *ℓ* = 12).

##### Connectivity optimized for delayed center-out reaching

network weights were optimized to produce joint torques for performing delayed center-out reaches with a biomechanical arm model. The architecture and optimization scheme followed that used by,^3^ in which the recurrent network is driven by two distinct inputs. The first input is a one-dimensional signal reflecting a go cue that indicates when the reach should be performed (go time). This was built into our model by setting *θ*_*K*_ to 1 at the start of the trial and then setting it to 0 at go time, 1000ms after trial start. The other input is a two-dimensional signal reflecting the visual presentation of the target to reach towards, presented prior to go time to prepare the subject (or network) to perform the delayed center-out reach. This was built into our model by setting *θ*_1_, *θ*_2_ to the coordinates of the reach target at a randomly sampled target presentation time before the go cue, and then setting it back to 0 at the same time the go cue input is shut off. All other command variables are set to 0 (*θ*_3_ = *θ*_4_ = … = *θ*_*K*−1_ = 0). We chose to encode the go cude with the very last command variable, *θ*_*K*_, to reflect the hypothesis that subjects would not re-aim with this non-directional command variable, neither in 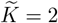 -dimensional re-aiming nor in generalized re-aiming with up to 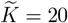 command variables.

Two joint torques were read out from the network through a set of readout weights, which were optimized along with the input, recurrent, and encoding weights. The weights were optimized to produce the joint torques required to move the endpoint of a planar two-link arm model^4^ to the cued reach target, in 500ms with a bell-shaped speed profile. Following the methods of,^5^ these target joint torques were computed by backpropagating through the arm model dynamics to minimize mean squared error between the arm endpoint velocity and the desired velocity profile for each reach target. We then trained the network weights so that in each trial it would produce 0 torque until go time, followed by the optimal reaching torque corresponding to the reach target on that trial.

The loss function used to optimize the network weights was a combination of the mean squared error plus L2 regularization on all weights and on network firing rates, to encourage naturalistic solutions to this task.^3^ This was minimized via stochastic gradient descent using the Adam optimization algorithm^6^ with learning rate set to .001. Since only three command variables were non-zero during this task, only the three corresponding columns of the encoding weights *U*_*ij*_ were optimized by this procedure. The remaining columns were thus fixed to their random initialization, as for the randomly connected network in the main text.

As is often observed in networks trained to perform a single task,^7^ the resulting optimized recurrent connectivity matrix had low rank (Supplementary Figure S1a). Its activity was consequently constrained to a much lower-dimensional subspace than that of the other networks.^8^ This network thus produced much lower-dimensional calibration task responses than the randomly connected networks did. We therefore used a 6-dimensional intrinsic manifold for constructing WMP’s and OMP’s in simulations with this network (i.e. *ℓ* = 6). This meant that only 6! − 1 = 719 possible decoder perturbations existed (as opposed to 40,319), so far fewer decoder perturbations satisfied the stringent criteria outlined in Methods Section 4.8 for sampling WMP’s and OMP’s. We thus loosened these criteria to include WMP’s and OMP’s with mean squared error going up to 1.2. These networks were also found to be highly sensitive to noise, so we reduced the standard deviation of the noise in the dynamics and in the initial conditions during simulation of the calibration task to 0.02 and 0.005, respectively.

Because of the low-rank recurrent connectivity, we found that re-aiming with even 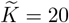 command variables could not yield good solutions for OMP control with this network (Supplementary Figure S1e). In other words, the low-rank connectivity did not permit the generation of activity patterns outside of the intrinsic manifold, even when re-aiming with a large nubmer of command variables. It is important to keep in mind, however, that in reality motor cortical connectivity is likely optimized to perform a wide variety of motor behaviors, rather than a single center-out reaching task. This assumption is implicit in our choice of high-rank connectivity structure, as in several other recent models of motor cortical function.^2,5,9^ Note also that such high-rank connectivity was necessary for our model to produce calibration task responses with dimensionality near the dimensionality of 10 observed by Sadtler et al. (2014) (Supplementary Figure S1c).

#### S.1.2 Analysis of WMP bias

This section provides a more detailed analysis of why WMP reachable readouts are biased in the direction of 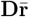 (fig. 4c). Specifically, we show that the reachable readouts are centered away from the origin, and the direction of this displacement is dictated by the relationship between the reachable manifold and the calibration task activity that the decoders are fit to, leading to the observed bias.

We begin by calculating the centroid of the reachable readouts, 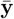,which is given by the readout of the reachable manifold centroid, 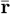,

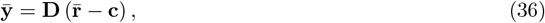

where **c** is the decoder centering vector (equation 3), set to the mean population response during the calibration task (Methods Section 4.8). As long as the reachable readouts are somewhat symmetrically distributed around their centroid, then they will be biased in the direction of their centroid. This equation shows that the direction and magnitude of this bias thus depends on the difference between between the mean calibration task response, **c**, and the reachable manifold centroid, 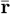.

Because the calibration task responses are driven by a subset of the motor commands used to define the reachable manifold (fig. 3e), these two directions are highly aligned. This is shown empirically for our simulations in Supplementary Figure S2a, where we overlay the network’s firing rates during individual trials of the calibration task on top of the reachable manifold, along with the mean population response and the reachable manifold centroid. Population activity during the calibration task evidently evolves along the same directions in state space occupied by the reachable manifold, and thus its mean, **c**, is highly aligned with the manifold’s centroid, 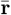.

Next, note that the reachable manifold centroid has a larger norm than the calibration task mean. That is because the reachable manifold contains activity patterns generated by motor commands with larger norms than those driving the calibration task responses (fig. 3e), so its centroid comprises higher firing rates. The underlying reason for this is that, in our simulations, we selected the metabolic cost weight, *γ*, such that it guaranteed high re-aiming performance with the baseline decoder (see Methods Section 4.3). Because the baseline decoder fit is not perfect (due both to the noise and to the non-linear mapping from calibration task stimuli to neural activity), stronger firing rates than those evoked by the calibration task are required to achieve such high performance, so the resulting metabolic cost weight permits stronger motor commands than the ones driving the calibration task.

Putting these two observations together, we have that

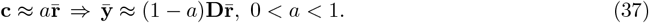

In other words, the reachable readout centroid points in the direction of 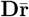.This explains why WMP readouts are biased in that direction.

An important remaining question, however, is why *baseline* decoder readouts are not biased in that direction. The baseline decoder shares the same centering vector, **c**, so, by the above logic, should inherit the same bias. The reason it does not is that the reachable manifold centroid is orthogonal to the baseline decoder, 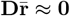, so the reachable readouts are centered at the origin despite equation 37 holding true. Because the baseline decoder is fit to predict the calibration task stimuli from the neural responses they elicit (Methods Section 4.8), by construction it ignores any directions of calibration task activity that do not provide information about the stimulus. One such direction is their mean, 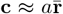.This can be appreciated from Supplementary Figure S2a, where it is evident that the trajectories of activity during different trials of the calibration task all evolve identically along this direction, despite being evoked by different stimuli. Thus, decoding from this direction is useless for decoding the stimulus identity, so the baseline decoder ignores it by spanning an orthogonal subspace. However, while this direction may not contain information about the calibration task stimuli, it does contain a lot of the variance of the calibration task neural responses. Consequently, it resides within the intrinsic manifold, and thus WMP’s – which are essentially randomly oriented within the intrinsic manifold – are likely to be aligned with it by chance.

**Figure S2:**
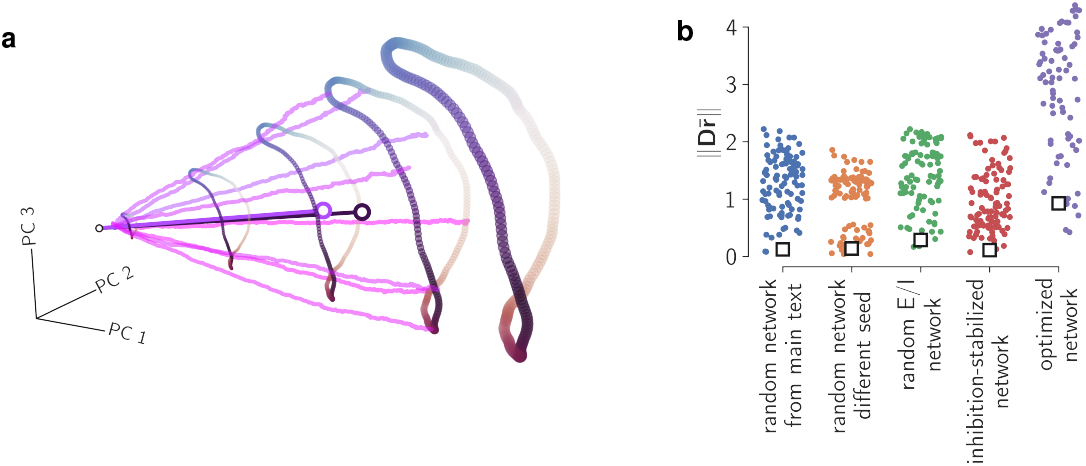
Analysis of WMP readout bias in the direction of 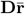. a. Simulated motor cortical responses in the calibration task, color-matched to the motor commands in fig. 3e driving these responses. These are plotted together with the same reachable manifold activity patterns from fig. 3f, projected onto the same three principal components. The open circles in the interior of this conical structure show the calibration task mean, **c**, in light purple and the reachable manifold centroid, 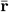, in dark purple, each connected to the origin by a line for visualization purposes. Note that, by definition, the calibration task neural responses at time *t*_end_ = 1000ms (the last point in each trajectory) lie almost exactly on the reachable manifold, slightly offset only because of noise in the response dynamics (see Methods Section 4.7). b. Norm of 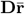 for each sampled WMP decoder, **D**, for each simulated model motor cortical network (see Supplementary Materials Section S.1.1). Overlaid with an open black square is the norm of 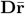 for the baseline decoder of each corresponding simulation.

This is confirmed empirically in fig. S2b, where we plot the norm of 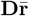 for the baseline decoder and each of the sampled WMPs for each simulation we studied. The norm is evidently much higher for WMP decoders than for the baseline decoder in each simulation, explaining why the reachable readout bias manifests itself in WMP decoders but not in the baseline decoder.

#### S.1.3 Non-negative firing rates are necessary to replicate biases in WMP learning

Here we demonstrate that removing the non-negativity constraint on firing rates precludes our model from reproducing the behavioral biases in WMP reachable readouts. We test this by replacing the activation function with the identity function, *ϕ*(*x*) = *x*, and repeating the analysis of fig. 4c to check if the maximal cursor progress under each WMP is highest in the direction of 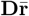.We find that they in fact are not (Supplementary Figure S3a)

**Figure S3:**
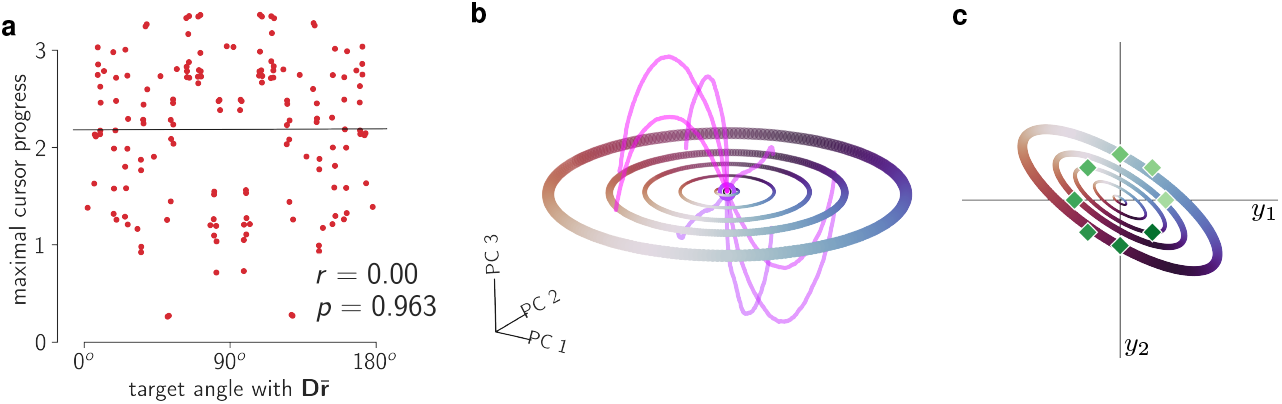
Removing the non-negativity constraint fails to reproduce experimentally observed biases in WMP learning. a. Maximal cursor progress in each target direction as a function of angle with 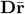,for 20 sampled WMPs. Only 20 WMP’s were used as we found that the intrinsic manifold of the linear network was only *ℓ* = 5-dimensional, so we correspondingly adjusted the criteria for subsampling WMP’s and OMP’s (cf. Supplementary Materials Section S.3.3) and found that only 20 WMP’s and 60 OMP’s satisfied them. As was done in fig. 4c, the reachable manifold centroid, 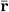, is estimated using simulated mean firing rates during baseline decoder control (see Methods Section 4.6). Maximal cursor progress was calculated exactly as in equation 6, following the same procedure as in the main text for selecting *s*_max_ (cf. Methods Section 4.4). A total of 8 target directions × 20 sampled WMPs = 160 points are plotted. b. Activity patterns in the reachable manifold at endpoint time *t*_end_ = 1000ms, with 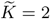non-zero command variables. Calibration task responses to each of the eight radial reach stimuli are overlaid in shades of pink, following exactly the same conventions as in Supplementary Figure S2a. These *N* -dimensional activity patterns are projected onto the top two principal components of the reachable manifold (PC1 and PC2) and the orthogonal dimension capturing the most calibration task response variance (PC3). Because the network dynamics are linear, the reachable manifold is exactly 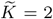-dimensional, so PC1 and PC2 capture 100% of the variance in activity patterns within it. The small open black circle at the center marks the origin of the state space. The light and dark purple open circles at the origin mark the calibration task mean **c** the reachable manifold centroid 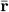 in purple, respectively. The latter is barely visible because they overlap almost completely. c. Readouts reachable through an example WMP, following the same color conventions as in fig. 3c. The green diamonds show the eight target readouts from the radial cursor reaching task.

The reason why can again be gleaned from looking at the relationship between the reachable repertoire centroid, 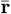,the mean calibration task response, **c**. In this case, because firing rates are allowed to be negative, both the reachable manifold and the calibration task neural responses are centered near the origin, and thus the reachable readouts are as well (equation 36),

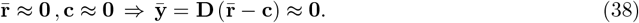

We can see this empirically in Supplementary Figure S3b, where we plot a projection of the reachable manifold with the calibration task neural responses overlaid (analogous to Supplementary Figure S2a). We see that both are centered around the origin, with their means exactly on top of each other. The reachable readouts through a representative WMP are shown in Supplementary Figure S3c, illustrating the fact that they are consequently also centered at the origin. The maximal cursor progress is higher in some directions than in others (in this case in the NW and SE directions), but the bias is not unidirectional as it is in the model with non-negative firing rates (fig. 4a, fig. 4c) or in the experimental data (fig. 4d).

**Figure S4:**
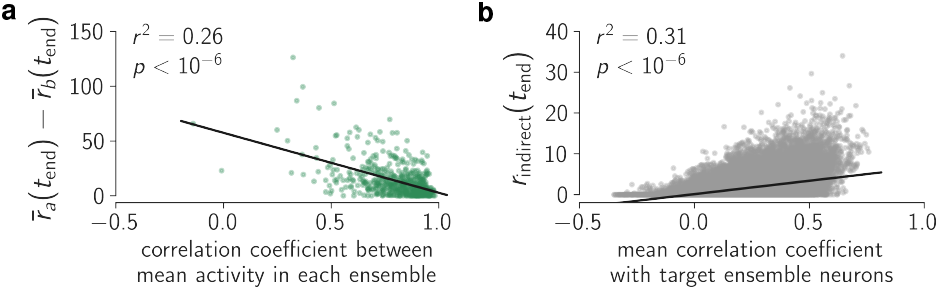
Operant conditioning of ensembles of 10 neurons by re-aiming. a. Difference in ensembles’ summed firing rates (at *t*_end_ = 1000 ms) produced by the re-aiming solutions optimized for ensemble *a* being the target, for 500 randomly sampled ensembles of 10 neurons, plotted as a function of the correlation between the two ensembles’ summed firing rates during simulated spontaneous behavior. Here, 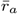 and 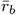 denote the summed firing rates of neurons in ensemble *a* and *b*, respectively. b. Activity of indirect neurons (at *t*_end_ = 1000 ms) for the same ensembles and re-aiming solutions in previous panel, plotted as a function of mean correlation coefficeint with neurons in ensemble *a* during simulated spontaneous behavior.

#### S.1.4 Operant conditioning of ensembles of neurons

Clancy et al. (2014) conditioned ensembles of up to 10 neurons, rather than only pairs of neurons. Here we repeat our simulations of operant conditioning but with a pair of target/distractor *ensembles* of 10 neurons, rather than a pair of single neurons. We simulate re-aiming in exactly the same way via equation 29, but where now the decoding vector **d** is a vector with a +1 for each target ensemble neuron and a -1 for each distractor ensemble neuron, and 0’s everywhere else. Selection of neurons used for operant conditioning and simulation of spontaneous behavior was done exactly as described in Methods Section 4.10.

We find that the main results from the main text are replicated in this setting as well: (1) the correlation coefficient of ensembles’ mean firing rates during simulated spontaneous behavior is correlated with the difference in mean firing rates achieved by the optimal re-aiming solutions (Supplementary Figure S4a); and (2) indirect neuron firing rates produced by these optimal re-aiming solutions are correlated with their mean correlation coefficient with target ensemble neurons during simulated spontaneous behavior (Supplementary Figure S4b).

#### S.1.5 Changes in modulation depth of motor cortical tuning curves

In a study employing credit assignment rotation perturbations in a 3D cursor reaching task, it was observed that both non-rotated and rotated neurons reduced their modulation depth after learning the perturbed decoder, and that rotated neurons reduced their modulation depth more^10^ (see text surrounding Methods Section 4.9, equation 26 for how modulation depth is defined and measured). Figure S5a reveals that our model of generalized re-aiming does not reproduce this result, at least for the values of 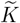 we tested. Generalized re-aiming with up to 6 command variables seems to lead to slight *increases* in the modulation depths of both rotated and non-rotated neurons, with marginal differences between rotated and non-rotated neurons.

While we did not simulate re-aiming with 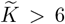 command variables in the context of credit assignment rotation perturbations, we did simulate this in the context of OMP learning, where we found that modulation depths of indirect neurons decreased as 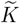 increased (figure S5b). The fact that only generalized re-aiming with a large number of command variables – but not regular re-aiming with only 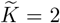 variables – can reproduce selective changes in modulation depth is consistent with the separate observation that indirect neurons show selective decreases in modulation depth only after days of practice with a given BCI decoder, but not within a single session.^11^

**Figure S5:**
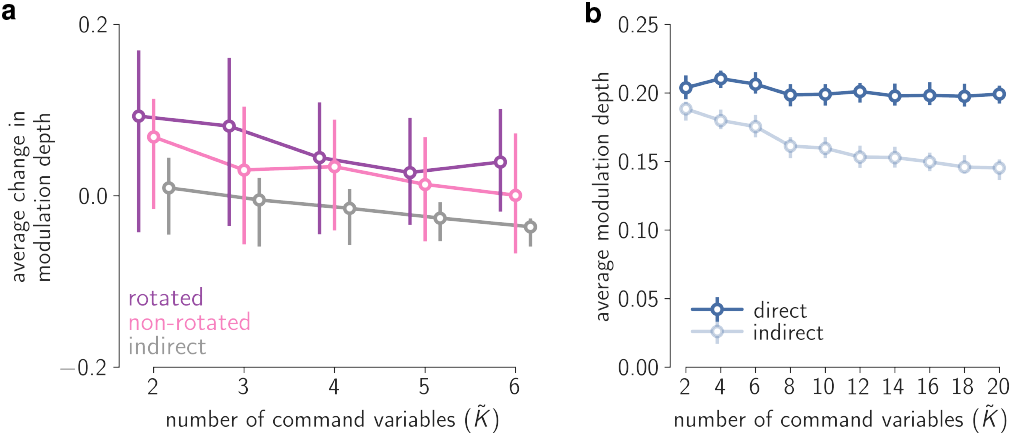
Changes in modulation depth under generalized re-aiming. a. Average change in modulation depth of rotated, non-rotated, and indirect neurons between simulated reaches with the baseline decoder and each perturbed decoder, plotted as a function of the number of command variables used for re-aiming 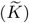.As in fig. 6e, the changes in modulation depth are averaged over all neurons in each sub-population, and the median over all 100 sampled credit assignment rotation perturbations is plotted. Error bars mark the upper and lower quartiles. b. Average modulation depth of direct neurons (neurons recorded by the BCI, with non-zero decoding weights in **D**) and indirect neurons (neurons not recorded by the BCI) under generalized re-aiming solutions for OMP’s. Modulation depths are averaged over all neurons in each sub-population, and the median over all 100 sampled OMPs is plotted. Error bars marks the upper and lower quartiles.

#### S.1.6 Closed-loop feedback control

All of the models we have considered so far are models of *open-loop* control: once the optimal motor command is specified, it is used to drive the motor cortical population for the duration of the movement, unchanged until the pre-specified endpoint time *t*_end_. Any errors encountered along the way – either due to noise or suboptimal motor commands – are thus ignored. A better strategy would be *closed-loop* control, wherein errors observed via sensory feedback are used to adaptively modify the motor command online. Under this strategy, errors that are encountered along the way can be corrected, thus improving the accuracy of the desired BCI output. Such closed-loop feedback control strategies are well known to be optimal in the presence of noise,^12,13^ and substantial evidence exists that non-human primates utilize them during BCI control.^14–16^

In this section, we consider a re-aiming-based model of closed-loop feedback control in which the motor command is continuously updated in response to sensory feedback. We evaluate this model on the Sadtler et al. (2014) BCI learning task, and confirm that closed-loop re-aiming suffers from the same limitations as open-loop re-aiming: the set of reachable activity patterns is limited by the number of command variables used for control, such that OMP’s cannot be learned with a small number of them.

We assume an error feedback controller architecture of the following form,

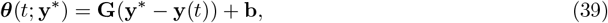

where the command variables, 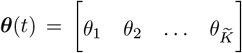,vary continuously in time according to an affine transformation of the instantaneous error, **y**^*^ − **y**(*t*). As in the open-loop control simulations, all additioanl command variables beyond the first 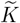 are fixed to 0. For simplicity, we assume a linear encoding of the motor command in the upstream inputs,

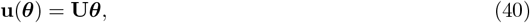

For a given decoder, **D**, we postulate that the subject learns a feedback controller, 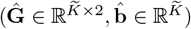,that minimizes the following loss function:

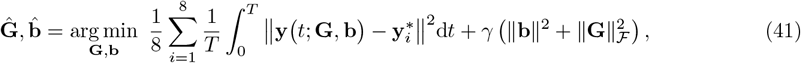

where **y** *t*; **G, b** denotes the readout produced at time *t* under the closed-loop dynamics 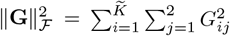 is the squared Frobenius norm, and 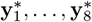 are the eight radial reach targets in the BCI cursor control task. The time window of control was set to *T* = 1000ms.

To compute 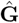 and 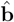,we used gradient descent on the above loss function, using the Adam optimization algorithm^6^ with a initial learning rate of .01. To facilitate numerical optimization, deterministic dynamics were used (no noise in the dynamics or in the initial conditions, which were fixed to 0). To avoid poor local optima (which was often a problem with WMPs in particular), we ran gradient descent from five different random initializations and used the best solution from these five runs.

We computed 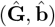 for the randomly connected non-linear network analyzed in the main text (equation 1), for each of the sampled baseline/WMP/OMP decoders used in the simulations presented in the main text. For the baseline decoder we performed this optimization over multiple values of the metabolic cost weight *γ* so as to identify the largest value of *γ* that permitted a time-averaged squared error of less than .05 for all eight target readouts under this decoder (analogous to how *γ* was set in the open-loop simulations in the main text, Methods Section 4.3). We then fixed *γ* to this value for all the decoder perturbations.

Simulations of cursor control with 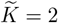 command variables are shown in Supplementary Figure S6a, where we plot the mean squared error over target readouts as a function of time. Each trace corresponds to the mean squared error achieved by the optimal feedback controller for a given decoder, with one trace for the baseline decoder and one for each of the 100 sampled WMPs and OMPs. We find that for almost all decoders, the mean squared error decreases to a certain level and remains low for the rest of this time window of 1000ms. However, this asymptotic error value is typically higher for OMP’s than for WMP’s (Supplementary Figure S6b), replicating the analogous result observed for the open-loop control model presented in the main text (fig. 3d).

**Figure S6:**
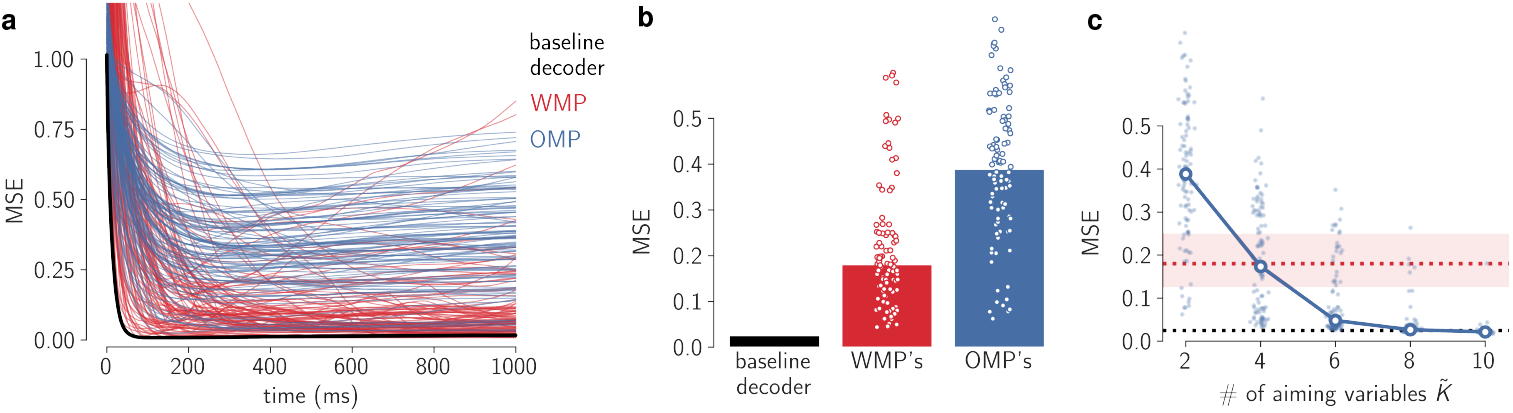
Closed-loop re-aiming reproduces the differences in WMP and OMP learning. a. Mean squared error (mean over target readouts) achieved by closed loop control with 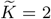 command variables, as a function of time. Each line corresponds to performance on a different decoder, with a correspondingly optimized feedback controller, 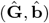 (equation 41). b. Mean squared error (mean over target readouts and over time) achieved by error feedback controllers with 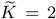 command variables. Each point corresponds to a different decoder, with medians over all decoders in each class marked by the height of the bars. c. Mean squared error (mean over target readouts and over time) achieved by error feedback controllers optimized for each OMP, with 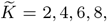 and 10 command variables. Light blue points denote this quantity for individual OMP’s, larger open circles on top show the median. For reference, dotted horizontal lines show the mean squared error achieved by optimized error feedback with 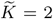 command variables for the baseline decoder (black) and WMP’s (red); the red dotted line shows the median over all sampled WMP’s with shading marking the upper and lower quartiles.

This again reflects the limitations of re-aiming with only 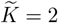 command variables. In this case, this manifests itself in restricting how the error can be fed back into the network: the error gets mapped to a 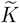-dimensional vector through equation 39 before being fed back into the network. As we saw occurs for the open-loop controller, this results in a restriction of how population activity can be modulated, making it difficult to generate the patterns of activity required to produce the target readouts under OMP’s. Supplementary Figure S6c shows that these restrictions can be relaxed by increasing the number of command variables used for re-aiming, 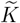.In this case, re-aiming with only 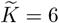 command variables suffices to obtain a mean squared error less than 0.1 with OMP’s. Interestingly, this is substantially less than the 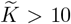 command variables that are necessary to achieve the same level of performance with open-loop re-aiming (fig. 5a).

### S.2 Mathematical derivations

#### S.2.1 Scale-invariance of RNN dynamics with rectified linear activation function

Here we prove that, whenever *x*_*i*_(0) = 0,

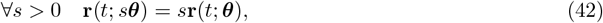

where **r**(*t*; *s****θ***) = *ϕ*(**x**(*t*; *s****θ***)) and **x**(*t*; *s****θ***) is the solution to equation 1 with inputs defined by equation 2. The function *ϕ*(·) is the rectified linear activation function defined in equation 1.

We begin by demonstrating that, when *x*_*i*_(0) = 0,

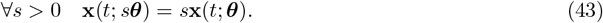

We prove this by showing that the dynamics of *s***x**(*t*; ***θ***) are the same as those of **x**(*t*; *s****θ***):

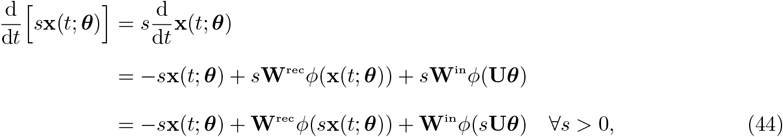

where in the second line we plugged in equation 1 and equation 2 for the dynamics and upstream inputs, respectively, and in the third line we used the scale-invariance of the rectified linear activation function,

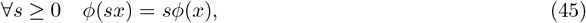

It is easy to see that equation 44 exactly matches equation 1 but with *s***x** substituted in for **x**; that is, the dynamics of these two quantities are the same. Therefore, whenever the initial conditions match, *s***x**(0; ***θ***) = **x**(0; *s****θ***), then their trajectories will too. It is easy to see that this condition holds for any *s* if *x*_*i*_(0) = 0, thus proving equation 43.

Along with the scale invariance of the activation function (equation 45), equation 43 implies equation 42:

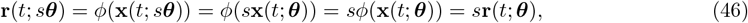

thus completing our proof.

#### S.2.2 Large *M* limit of quadratic metabolic cost

Here we derive the large *M* limit of the quadratic metabolic cost term in the reaiming objective function (equation 4),

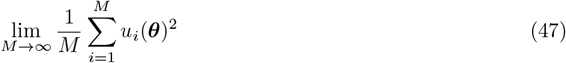

We first note that each term in the sum depends on a sum over the randomly sampled encoding weights (equation 2),

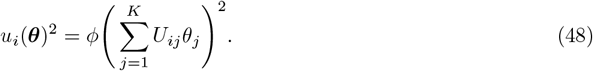

If the encoding weights, *U*_*ij*_, are independent and identically distributed, then each of the terms in this sum is also independent and identically distributed. By the law of large numbers, then, as *M → ∞* their sum will approach an expectation over this distribution,

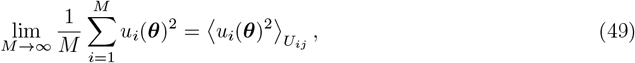

where 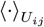 denotes an expectation over the probability distribution of the encoding weights, *U*_*ij*_.

This expectation can be evaluated by first defining the random variable 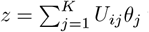 to express the expectation as an integral over *z*, and then exploiting the rectified linear activation function (equation 1) to simplify this integral,

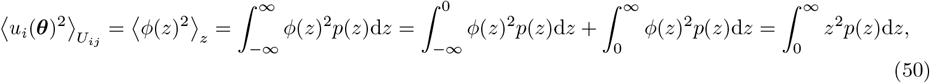

where we simply exploited the fact that *ϕ*(*z*) = 0 when *z <* 0 and *ϕ*(*z*) = *z* when *z* ≥ 0. If the distribution of the encoding weights *U*_*ij*_ is symmetric around 0, then the distribution of *z* is as well and we have that

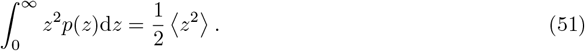

Finally, if the encoding weights are zero-mean and independent, we have that

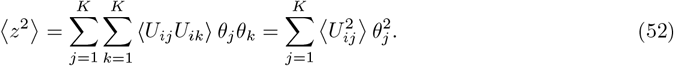

If *U*_*ij*_ additionally have unit variance, 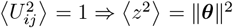.Putting this all together, we arrive at the equality in equation 9:

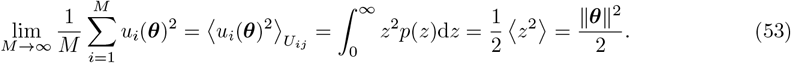

#### S.2.3 Reachable manifold moments for 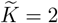

In our analysis of the reachable manifold, we characterized its location and shape via its centroid and covariance, which were evaluated as expectations over a uniform distribution on the manifold. Here we derive the probability density function of this distribution and use it to calculate these expections.

We begin with the case of 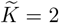 non-zero command variables, which we parameterize by their polar coordinates,

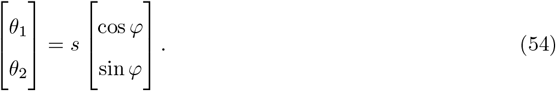

We then formally define the reachable manifold as follows:

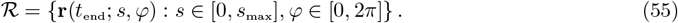

where **r**(*t*_end_; *s, φ*) is the motor cortical activity pattern at time *t*_end_ produced by a pair of command variables *θ*_1_, *θ*_2_ with angle *φ* and norm *s*, with all other command variables set to 0 (*θ*_3_ = *θ*_4_ = … = *θ*_*K*_ = 0). The function **r**(*t*_end_; *s, φ*) can be thought of as a function mapping 2D command variables, (*s, φ*) ∈ [0, *s*_max_] × [0, 2*π*], to activity patterns, **r** ∈ ℝ _*N*_, on the 2D surface constituting the reachable manifold (the conical surface shown in fig. 3f).

The probability density function of the uniform distribution on this 2D surface in ℝ _*N*_ is given by its area element, *dV* (*s, φ*), divided by its total area,

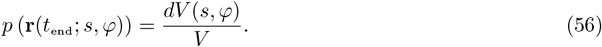

The area element and total area are given by

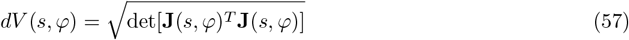

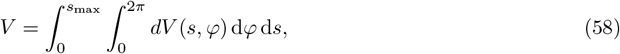

where det[°] denotes the matrix determinant and **J** denotes the *N* × 2 Jacobian of the mapping from command variables to the reachable manifold,

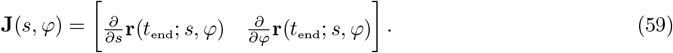

To evaluate the probability density function, we must first calculate these derivatives.

To do so, we again resort to the scale invariance property of the rectified linear activation function (equation 45),

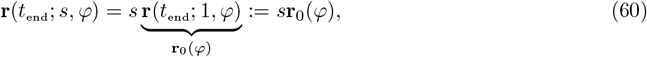

where we have defined **r**_0_(*φ*) to be the activity generated by a pair of command variables with angle *φ* and unit norm. The Jacobian is thus given by

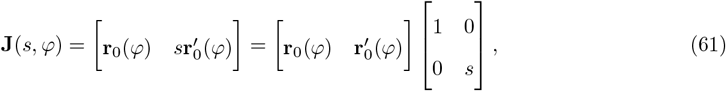

where 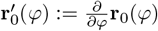.Plugging this into equation 57, we have that the area element is given by

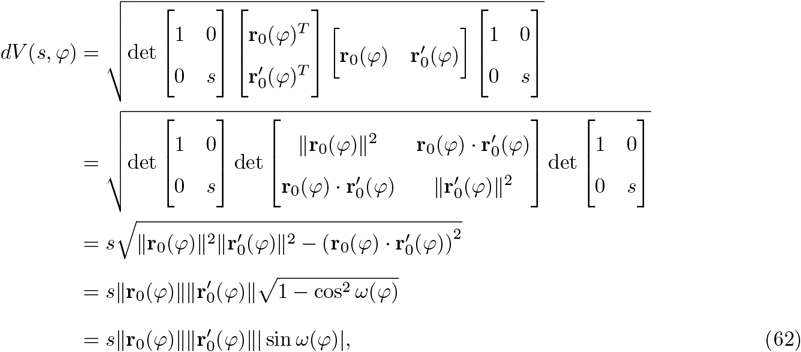

where *ω*(*φ*) is the angle between **r**_0_(*φ*) and its derivative at 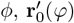.The total area of the manifold is thus

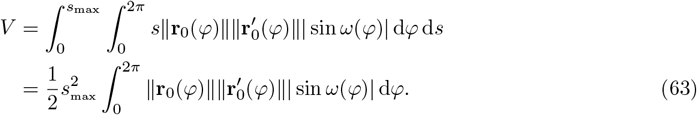

With these two expressions in hand, we can analytically express expectations over the probability density function in equation 56. The mean, corresponding to the manifold centroid, 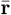,is given by

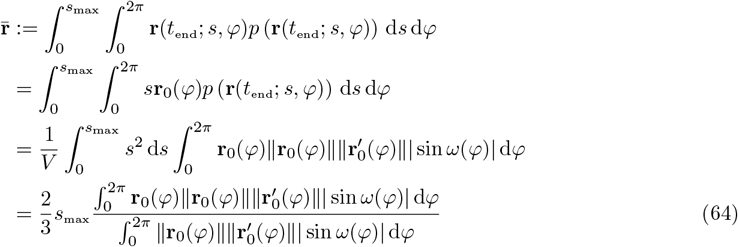

Its covariance, **Σ**_*r*_, is given by

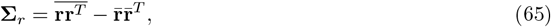

where 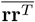 is the matrix of second moments,

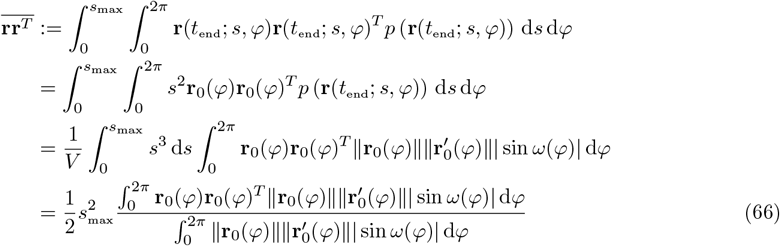

Because the integrals and derivatives in these expressions are all univariate, we can estimate them accurately with discrete approximations.

#### S.2.4 Reachable manifold moments for 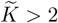

Analagous expressions can be derived for the case of 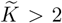 but in these cases good estimates of the integrals and derivatives quickly become numerically intractable as the number of variables increases. For these cases, we therefore resorted to moments with respect to the probability distribution of activity patterns *generated by uniformly distributed motor commands*, instead of the probability distribution of activity patterns *uniformly distributed on the reachable manifold*.

We can express the covariance of this simpler distribution, which we denote by **Σ**_*θ*_, by parameterizing the non-zero command variables, 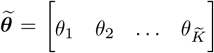,in terms of a magnitude and direction, 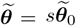,where 0 ≤ *s* ≤ *s*_max_ and 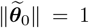. This allows us to factorize the uniform distribution over motor commands into a scalar uniform distribution for the magnitude, *s* ∼ Unif[0, *s*_max_], and a uniform distribution over the unit radius 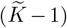-sphere for the direction, 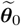. The expectations in the covariance thus factorize as follows:

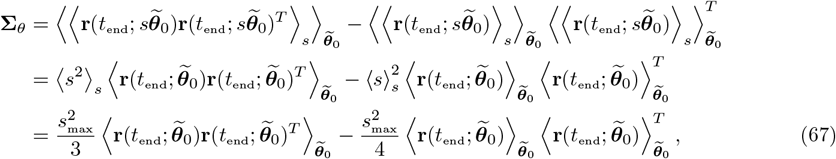

where we used the scale invariance of the motor cortical dynamics (equation 8) to write 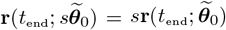 in the second line, and in the third line we simply inserted expressions for the first and second moments of *s*. The expectations over 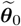 can be approximated using Monte Carlo methods by uniformly sampling vectors from the corresponding unit radius 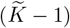-sphere.

### S.3 Extended methods

#### S.3.1 Estimating the intrinsic manifold

To estimate the intrinsic manifold, we fit a Probabilistic PCA (PPCA) model^17^ to the mixed and z-scored calibration task responses (see Methods Section 4.8),

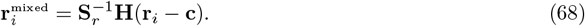

Here, *i* indexes a particular timestep and trial during the calibration task. The PPCA generative model assumes that each of these data points are generated from a corresponding set of *ℓ* uncorrelated latent variables 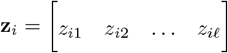 as follows,

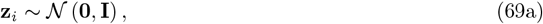

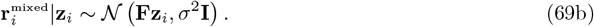

The model thus assumes that the activity patterns 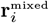 are concentrated within the column space of the factor loading matrix **F** – it is the columns of this matrix that define the intrinsic manifold. These parameters are fit to the mixed and z-scored calibration task data,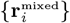, by maximum likelihood:

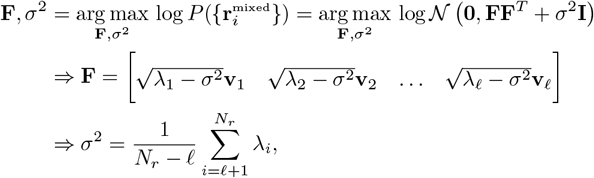

where 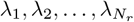 are the eigenvalues of the sample covariance of the calibration task activity, 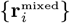, ordered from largest to smallest (i.e. *λ*_1_ is the largest eigenvalue), and 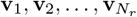 are their associated eigenvectors (i.e. the principal components, ordered from most to least variance explained).

Note, however, that the columns of **F** define the dimensions of the intrinsic manifold in mixed and z-scored neural activity space (i.e. the space defined by the coordinates of the 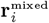 vectors). To convert these to dimensions of the full *N* -dimensional state space, where each coordinate corresponds to the activity of an individual neuron (i.e. the space defined by the coordinates of the **r**_*i*_ vectors), we invert equation 68 to obtain

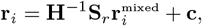

where we define **H**^−1^ as the *N* × *N*_*r*_ matrix containing the inverse of the tri-diagonal component of **H** in its first *N*_*r*_ rows and 0’s filling all subsequent rows. We then apply this linear transformation to the columns of **F** to obtain an analagous *N* × *ℓ* factor loading matrix **F**_*r*_ defined in the full *N* -dimensional state space,

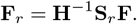

Note that, since the bottom *N* − *N*_*r*_ rows of **H**^−1^ are filled with 0’s, those same rows of **F**_*r*_ are also filled with 0’s. This reflects the fact that the intrinsic manifold is orthogonal to the dimensions of activity corresponding to neurons not recorded in the experiment. Finally, we defined an orthonormal basis **f**_1_, **f**_2_, …, **f**_*ℓ*_ ∈ ℝ _*N*_ for the intrinsic manifold by taking the left singular vectors of **F**_*r*_. These are the vectors used in equation 25 for figure 3g.

This method for estimating the intrinsic manifold is almost the same as that used by Sadtler et al., which differs only in that a Factor Analysis model was used instead of a PPCA model. In that case, the maximum likelihood estimates of the model parameters cannot be evaluated in closed form and must be computed via an iterative optimization algorithm (the Expectation Maximization algorithm). We found that using a Factor Analysis model instead of PPCA had no noticeable effects on our results (data not shown), so we reported only results with the more easily fit PPCA model.

#### S.3.2 Construction of the baseline decoder

As described in the Methods section, the baseline decoder has the following form

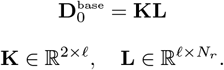

We term **L** the *dimensionality reduction matrix* and **K** the *velocity readout matrix*. Here we describe in greater detail how these two matrices are fit to the calibration task data. Unless otherwise noted, these procedures are exactly as those described in^18^ and.^19^

The dimensionality reduction matrix **L** is derived from the mode of the posterior distribution of the PPCA model (equation 69),

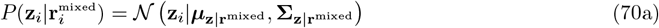

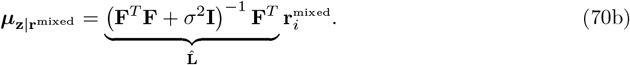

The *N*_*r*_ × *ℓ* matrix 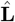 thus yields a linear transformation from *N*_*r*_ dimensions to *ℓ* dimensions. The z-scored and mixed activity patterns 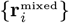 from the calibration task can thus be reduced to *ℓ* dimensions via multiplication with 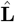,resulting in a corresponding set of dimensionality-reduced activity patterns 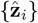 (as above, here and in the rest of this section the index *i* jointly indexes a timestep and trial of the calibration task).

To complete the construction of the dimensionality reduction matrix **L**, these dimensionality-reduced activity patterns are then z-scored. The standard deviations of each component of the 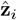 vectors are calculated over all timesteps and trials of the calibration task, and collected in a diagonal matrix **S**_*z*_. Note that mean subtraction is not necessary since the activity vectors 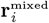 have already been z-scored so are mean 0. The final dimensionality reduction matrix is then given by

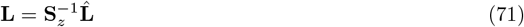

This second z-scoring step is necessary to ensure that controlling the BCI does not require neurons to produce firing rates beyond the range exhibited during the calibration task.

The dimensionality reduction matrix used by Sadtler et al. differed from ours in that 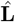 was constructed from the posterior distribution under a Factor Analysis generative model, rather than a PPCA generative model. Like in PPCA, the mode of the posterior distribution of a Factor Analysis model can also be expressed as a linear transformation of 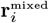,yielding a very similar expression for 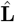.

The velocity readout matrix **K** is also chosen by maximum likelihood fit of a generative model. In this case, we assume that the z-scored dimensionality-reduced activity patterns from the calibration task, 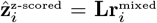,depend on the observed cursor velocities, **y**_*i*_, via the following latent Gaussian state space model,

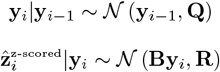

where *i*−1 indexes the previous timestep in the same trial. Note that the cursor velocities **y**_*i*_ are constant within each trial of the calibration task, so within a given trial **y**_*i*_ = **y**_*i*−1_. As was done in the original experiment of Sadtler et al., we set

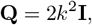

where *k* = 1*/*.15 denotes the ratio of the cursor speeds used in our simulation (∥**y**_*i*_∥ = 1) and the cursor speeds used in the original experiment (∥**y**_*i*_∥ = .15 m/s). Maximum likelihood estimates of the remaining parameters are given by

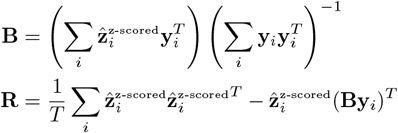

where *T* denotes the total number of data points in the calibration task data: the number of timesteps in each trial times the total number of trials.

**Figure S7:**
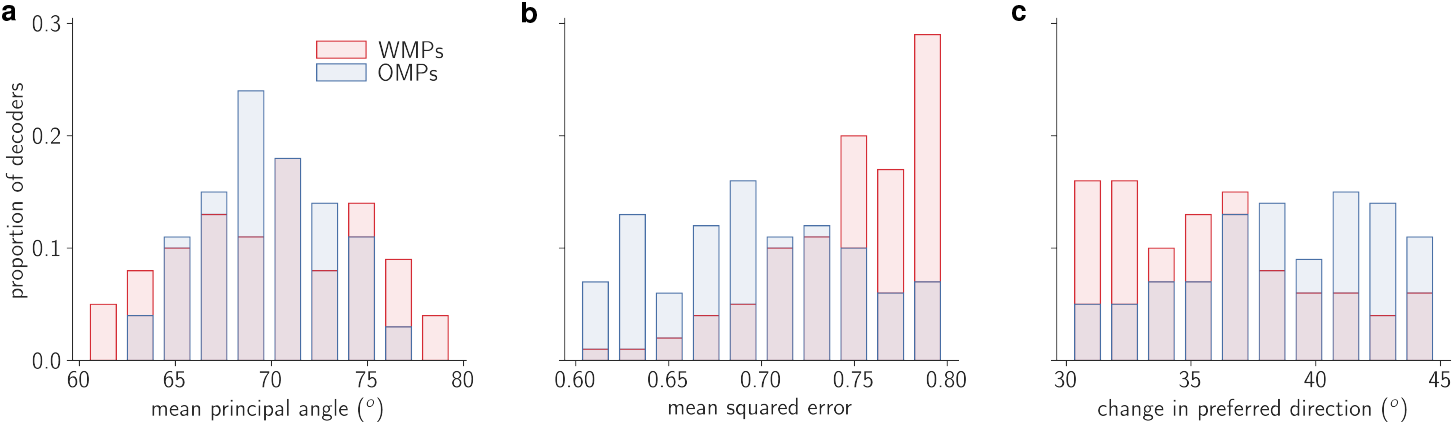
Differences between sampled decoder perturbations and the baseline decoder. a. Distribution of mean principal angle between row space of baseline decoder and row space of each perturbed decoder. b. Distribution of mean squared error achieved by mean calibration task responses under each perturbed decoder. c. Distribution of minimal absolute change in preferred direction needed to produce the same readouts with each perturbed decoder as with the baseline decoder.

The velocity readout matrix is then derived from the mode of the of the posterior distribution 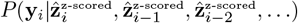,where the ellipses go back to the first timestep of the given trial. We use the posterior distribution at steady state, whose mode is given by

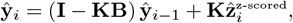

where **K** is the so-called steady-state Kalman gain matrix. This matrix is given by

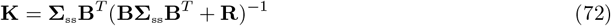

where **Σ**_ss_ is the steady-state posterior covariance, given by the solution to the discrete-time algebraic Riccatti equation

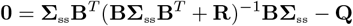

The 2 × *ℓ* velocity readout matrix used for the baseline decoder is thus set to the steady-state Kalman gain matrix, **K**.

#### S.3.3 Subsampling WMPs and OMPs

As mentioned in the methods, we attempted to minimize any differences between within- and outside-manifold perturbations that would go beyond their opposing relationship to the intrinsic manifold. To do this, we first calculated every possible WMP and OMP, corresponding to each *ℓ*-dimensional permutation. Since we set *ℓ* = 8, this resulted in *ℓ*! −1 = 40, 319 decoder perturbations of each type (minus 1 to exclude the identity permutation). We then quantified how different each of these perturbations were from the baseline decoder with three different metrics, and eliminated all decoder perturbations for which one or more of these metrics fell outside a specific range.

The first metric is the angle between the perturbed decoder’s row space and the baseline decoder’s.

For each decoder perturbation, 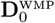 or 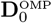,we calculated the two principal angles^20^ between its row space and that of the baseline decoder effective decoding matrix, 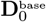,and averaged these two angles. Any decoder perturbations for which this mean principal angle was greater than 80^*o*^ or less than 60^*o*^ was eliminated (fig. S7a).

The second metric is the mean squared error that would be achieved if the subject were to simply reproduce the neural activity from the calibration task. Analagous to the procedure followed by Sadtler et al., we averaged the calibration task responses over time and over trials for each reach target,

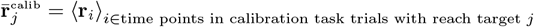

and then computed the readouts from these time- and trial-averaged firing rate vectors under each decoder perturbation, 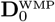 or 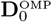.We then discarded all decoder perturbations where the mean

squared error between these readouts and the target readouts was greater than 0.8 or less than 0.6 (fig. S7b).

The third metric is to ask how much the mean calibration task responses would have to change to produce the same readouts under the perturbed decoder as under the baseline decoder. We first calculated the time- and trial-averaged z-scored and mixed firing rates from the calibration task

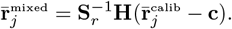

For each perturbed decoder, 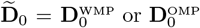,we then computed the activity patterns closest to 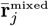 that would produce the same readouts through that decoder as the original activity patterns would through the baseline decoder, 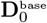,

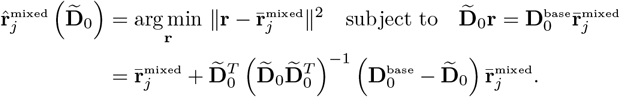

We then quantified the difference between 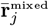 and 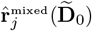 by fitting tuning curves and asking how much the preferred direction changed. Tuning curves were fit by least-squares regression, exactly as described in Methods Section 4.9 equation 26 (but with 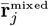 or 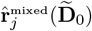 plugged in for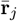), and preferred directions were extracted from the fitted tuning weights as described in that section. For each decoder perturbation, we then computed the mean absolute difference of the preferred directions of the computed activity patterns 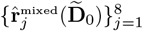 from those of the observed calibration task mean responses 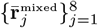. Any perturbed decoders that resulted in a mean absolute difference of more than 45^*o*^ or less than 30^*o*^ were discarded (fig. S7c).

We typically found that about 100-200 permutations out all possible decoder perturbations satisfied these criteria. We then randomly sampled 100 of them. The distributions of these three merics for the 100 sampled WMPs and OMPs used in the main text are shown in figures S7a, S7b, and S7c.

## References

[1] R. Shadmehr and F. A. Mussa-Ivaldi. “Adaptive Representation of Dynamics during Learning of a Motor Task”. en. In: Journal of Neuroscience 14.5 (May 1994), pp. 3208–3224. issn: 0270-6474, 1529-2401. doi: 10.1523/JNEUROSCI.14-05-03208.1994.

[2] Chiang-Shan Ray Li, Camillo Padoa-Schioppa, and Emilio Bizzi. “Neuronal Correlates of Motor Performance and Motor Learning in the Primary Motor Cortex of Monkeys Adapting to an External Force Field”. en. In: Neuron 30.2 (May 2001), pp. 593–607. issn: 0896-6273. doi: 10.1016/S0896-6273(01)00301-4.

[3] Rony Paz et al. “Acquisition and Generalization of Visuomotor Transformations by Nonhuman Primates”. en. In: Experimental Brain Research 161.2 (Feb. 2005), pp. 209–219. issn: 1432-1106. doi: 10.1007/s00221-004-2061-4.

[4] Gonçalo Lopes et al. “A Robust Role for Motor Cortex”. en. In: bioRxiv (May 2017), p. 058917. doi: 10.1101/058917.

[5] Risa Kawai et al. “Motor Cortex Is Required for Learning but Not for Executing a Motor Skill”. en. In: Neuron 86.3 (May 2015), pp. 800–812. issn: 0896-6273. doi: 10.1016/j.neuron.2015.03.024.

[6] Mackenzie Weygandt Mathis, Alexander Mathis, and Naoshige Uchida. “Somatosensory Cortex Plays an Essential Role in Forelimb Motor Adaptation in Mice”. en. In: Neuron 93.6 (Mar. 2017), 1493–1503.e6. issn: 0896-6273. doi: 10.1016/j.neuron.2017.02.049.

[7] Matthew G. Perich, Juan A. Gallego, and Lee E. Miller. “A Neural Population Mechanism for Rapid Learning”. en. In: Neuron 100.4 (Nov. 2018), 964–976.e7. issn: 0896-6273. doi: 10.1016/j.neuron.2018.09.030.

[8] Matthew D Golub et al. “Brain–Computer Interfaces for Dissecting Cognitive Processes Underlying Sensorimotor Control”. In: Current Opinion in Neurobiology. Neurobiology of Cognitive Behavior 37 (Apr. 2016), pp. 53–58. issn: 0959-4388. doi: 10.1016/j.conb.2015.12.005. (Visited on 04/27/2020).

[9] Richard A. Andersen, Tyson Aflalo, and Spencer Kellis. “From Thought to Action: The Brain–Machine Interface in Posterior Parietal Cortex”. In: Proceedings of the National Academy of Sciences 116.52 (Dec. 2019), pp. 26274–26279. issn: 0027-8424, 1091-6490. doi: 10.1073/pnas.1902276116. (Visited on 02/26/2020).

[10] Emily R. Oby et al. “Intracortical Brain–Machine Interfaces”. In: Neural Engineering. Ed. by Bin He. Cham: Springer International Publishing, 2020, pp. 185–221. isbn: 978-3-030-43395-6. doi: 10.1007/978-3-030-43395-6_5. (Visited on 07/30/2021).

[11] Beata Jarosiewicz et al. “Functional Network Reorganization during Learning in a Brain-Computer Interface Paradigm”. In: Proceedings of the National Academy of Sciences 105.49 (Dec. 2008), pp. 19486–19491. issn: 0027-8424, 1091-6490. doi: 10.1073/pnas.0808113105. (Visited on 02/04/2020).

[12] Steven M. Chase, Robert E. Kass, and Andrew B. Schwartz. “Behavioral and Neural Correlates of Visuomotor Adaptation Observed through a Brain-Computer Interface in Primary Motor Cortex”. In: Journal of Neurophysiology 108.2 (Apr. 2012), pp. 624–644. issn: 0022-3077. doi: 10.1152/jn.00371.2011. (Visited on 02/16/2020).

[13] Vikash Gilja et al. “A High-Performance Neural Prosthesis Enabled by Control Algorithm Design”. In: Nature Neuroscience 15.12 (Dec. 2012), pp. 1752–1757. issn: 1546-1726. doi: 10.1038/nn.3265. (Visited on 04/27/2020).

[14] Patrick T. Sadtler et al. “Neural Constraints on Learning”. In: Nature 512.7515 (Aug. 2014), pp. 423–426. issn: 1476-4687. doi: 10.1038/nature13665. (Visited on 02/13/2020).

[15] Karunesh Ganguly and Jose M. Carmena. “Emergence of a Stable Cortical Map for Neuroprosthetic Control”. In: PLOS Biology 7.7 (July 2009), e1000153. issn: 1545-7885. doi: 10.1371/journal.pbio.1000153. (Visited on 04/27/2020).

[16] Emily R. Oby et al. “New Neural Activity Patterns Emerge with Long-Term Learning”. In: Proceedings of the National Academy of Sciences 116.30 (2019), pp. 15210–15215.

[17] Robert Legenstein et al. “A Reward-Modulated Hebbian Learning Rule Can Explain Experimentally Observed Network Reorganization in a Brain Control Task”. In: Journal of Neuroscience 30.25 (June 2010), pp. 8400–8410. issn: 0270-6474, 1529-2401. doi: 10.1523/JNEUROSCI.4284-09.2010. (Visited on 03/13/2020).

[18] Ben Engelhard et al. “Neuronal Activity and Learning in Local Cortical Networks Are Modulated by the Action-Perception State”. In: bioRxiv (Feb. 2019), p. 537613. doi: 10.1101/537613. (Visited on 07/29/2020).

[19] Emil Wärnberg and Arvind Kumar. “Perturbing Low Dimensional Activity Manifolds in Spiking Neuronal Networks”. In: PLOS Computational Biology 15.5 (May 2019), e1007074. issn: 1553-7358. doi: 10.1371/journal.pcbi.1007074. (Visited on 04/27/2020).

[20] Barbara Feulner and Claudia Clopath. “Neural Manifold under Plasticity in a Goal Driven Learning Behaviour”. In: PLOS Computational Biology 17.2 (Feb. 2021), e1008621. issn: 1553-7358. doi: 10.1371/journal.pcbi.1008621. (Visited on 03/06/2021).

[21] Rodolphe Héliot et al. “Learning in Closed-Loop Brain–Machine Interfaces: Modeling and Experimental Validation”. In: IEEE Transactions on Systems, Man, and Cybernetics, Part B (Cybernetics) 40.5 (Oct. 2010), pp. 1387–1397. issn: 1941-0492. doi: 10.1109/TSMCB.2009.2036931.

[22] Justin Werfel, Xiaohui Xie, and H. Sebastian Seung. “Learning Curves for Stochastic Gradient Descent in Linear Feedforward Networks”. In: Advances in Neural Information Processing Systems. 2004, pp. 1197–1204.

[23] Naoki Hiratani et al. “On the Stability and Scalability of Node Perturbation Learning”. In: Advances in Neural Information Processing Systems 35 (Dec. 2022), pp. 31929–31941. (Visited on 05/24/2023).

[24] A. Aldo Faisal, Luc P. J. Selen, and Daniel M. Wolpert. “Noise in the Nervous System”. In: Nat Rev Neurosci 9.4 (Apr. 2008), pp. 292–303. issn: 1471-0048. doi: 10.1038/nrn2258. (Visited on 10/15/2023).

[25] Francis Crick. “The Recent Excitement about Neural Networks”. In: Nature 337.6203 (Jan. 1989), pp. 129–132. issn: 1476-4687. doi: 10.1038/337129a0. (Visited on 07/22/2020).

[26] Sergey Bartunov et al. “Assessing the Scalability of Biologically-Motivated Deep Learning Algorithms and Architectures”. In: Advances in Neural Information Processing Systems 31. Ed. by S. Bengio et al. Curran Associates, Inc., 2018, pp. 9368–9378. (Visited on 07/22/2020).

[27] Timothy P. Lillicrap et al. “Backpropagation and the Brain”. In: Nature Reviews Neuroscience 21.6 (June 2020), pp. 335–346. issn: 1471-0048. doi: 10.1038/s41583-020-0277-3. (Visited on 07/22/2020).

[28] EE Fetz and MA Baker. “Operantly Conditioned Patterns on Precentral Unit Activity and Correlated Responses in Adjacent Cells and Contralateral Muscles.” In: Journal of Neurophysiology 36.2 (Mar. 1973), pp. 179–204. issn: 0022-3077. doi: 10.1152/jn.1973.36.2.179. (Visited on 07/14/2020).

[29] Matthew D. Golub et al. “Learning by Neural Reassociation”. In: Nature Neuroscience 21.4 (Apr. 2018), pp. 607–616. issn: 1546-1726. doi: 10.1038/s41593-018-0095-3. (Visited on 11/11/2019).

[30] Jay A Hennig et al. “Constraints on Neural Redundancy”. In: eLife 7 (Aug. 2018). Ed. by Eric Shea-Brown and Timothy E Behrens, e36774. issn: 2050-084X. doi: 10.7554/eLife.36774. (Visited on 02/16/2020).

[31] Karunesh Ganguly et al. “Reversible Large-Scale Modification of Cortical Networks during Neuroprosthetic Control”. In: Nature Neuroscience 14.5 (May 2011), pp. 662–667. issn: 1546-1726. doi: 10.1038/nn.2797. (Visited on 04/13/2020).

[32] Saurabh Vyas et al. “Neural Population Dynamics Underlying Motor Learning Transfer”. In: Neuron 97.5 (Mar. 2018), 1177–1186.e3. issn: 0896-6273. doi: 10.1016/j.neuron.2018.01.040. (Visited on 02/16/2020).

[33] Steven M. Chase, Andrew B. Schwartz, and Robert E. Kass. “Latent Inputs Improve Estimates of Neural Encoding in Motor Cortex”. In: Journal of Neuroscience 30.41 (Oct. 2010), pp. 13873–13882. issn: 0270-6474, 1529-2401. doi: 10.1523/JNEUROSCI.2325-10.2010. (Visited on 07/14/2020).

[34] Eun Jung Hwang, Paul M. Bailey, and Richard A. Andersen. “Volitional Control of Neural Activity Relies on the Natural Motor Repertoire”. In: Current Biology 23.5 (Mar. 2013), pp. 353–361. issn: 0960-9822. doi: 10.1016/j.cub.2013.01.027. (Visited on 02/04/2020).

[35] Sofia Sakellaridi et al. “Intrinsic Variable Learning for Brain-Machine Interface Control by Human Anterior Intraparietal Cortex”. In: Neuron 102.3 (May 2019), 694–705.e3. issn: 0896-6273. doi: 10.1016/j.neuron.2019.02.012. (Visited on 02/16/2020).

[36] A. d’Avella and E. Bizzi. “Low Dimensionality of Supraspinally Induced Force Fields”. en. In: Proceedings of the National Academy of Sciences 95.13 (June 1998), pp. 7711–7714. issn: 0027-8424, 1091-6490. doi: 10.1073/pnas.95.13.7711.

[37] Andrea d’Avella, Philippe Saltiel, and Emilio Bizzi. “Combinations of Muscle Synergies in the Construction of a Natural Motor Behavior”. en. In: Nature Neuroscience 6.3 (Mar. 2003), pp. 300–308. issn: 1546-1726. doi: 10.1038/nn1010.

[38] Yuri P. Ivanenko et al. “Temporal Components of the Motor Patterns Expressed by the Human Spinal Cord Reflect Foot Kinematics”. In: Journal of Neurophysiology 90.5 (Nov. 2003), pp. 3555–3565. issn: 0022-3077. doi: 10.1152/jn.00223.2003.

[39] Emanuel Todorov. “Optimality Principles in Sensorimotor Control”. en. In: Nature Neuroscience 7.9 (Sept. 2004), pp. 907–915. issn: 1546-1726. doi: 10.1038/nn1309.

[40] Jason J. Kutch and Francisco J. Valero-Cuevas. “Challenges and New Approaches to Proving the Existence of Muscle Synergies of Neural Origin”. en. In: PLOS Computational Biology 8.5 (May 2012), e1002434. issn: 1553-7358. doi: 10.1371/journal.pcbi.1002434.

[41] Naveen Kuppuswamy and Christopher M. Harris. “Do Muscle Synergies Reduce the Dimensionality of Behavior?” English. In: Frontiers in Computational Neuroscience 8 (2014). issn: 1662-5188. doi: 10.3389/fncom.2014.00063.

[42] Mark M. Churchland et al. “Neural Population Dynamics during Reaching”. en. In: Nature 487.7405 (July 2012), pp. 51–56. issn: 1476-4687. doi: 10.1038/nature11129.

[43] Guillaume Hennequin, Tim P. Vogels, and Wulfram Gerstner. “Optimal Control of Transient Dynamics in Balanced Networks Supports Generation of Complex Movements”. en. In: Neuron 82.6 (June 2014), pp. 1394–1406. issn: 0896-6273. doi: 10.1016/j.neuron.2014.04.045.

[44] David Sussillo et al. “A Neural Network That Finds a Naturalistic Solution for the Production of Muscle Activity”. In: Nature Neuroscience 18.7 (July 2015), pp. 1025–1033. issn: 1546-1726. doi: 10.1038/nn.4042. (Visited on 04/22/2020).

[45] Abigail A. Russo et al. “Motor Cortex Embeds Muscle-like Commands in an Untangled Population Response”. en. In: Neuron 97.4 (Feb. 2018), 953–966.e8. issn: 0896-6273. doi: 10.1016/j.neuron.2018.01.004.

[46] Ta-Chu Kao, Mahdieh S. Sadabadi, and Guillaume Hennequin. “Optimal Anticipatory Control as a Theory of Motor Preparation: A Thalamo-Cortical Circuit Model”. en. In: Neuron 109.9 (May 2021), 1567–1581.e12. issn: 0896-6273. doi: 10.1016/j.neuron.2021.03.009.

[47] Mijail D. Serruya et al. “Instant Neural Control of a Movement Signal”. In: Nature 416.6877 (Mar. 2002), pp. 141–142. issn: 1476-4687. doi: 10.1038/416141a. (Visited on 08/16/2020).

[48] Leigh R. Hochberg et al. “Neuronal Ensemble Control of Prosthetic Devices by a Human with Tetraplegia”. In: Nature 442.7099 (July 2006), pp. 164–171. issn: 1476-4687. doi: 10.1038/nature04970. (Visited on 08/16/2020).

[49] Meel Velliste et al. “Cortical Control of a Prosthetic Arm for Self-Feeding”. In: Nature 453.7198 (June 2008), pp. 1098–1101. issn: 1476-4687. doi: 10.1038/nature06996. (Visited on 08/16/2020).

[50] Matthew D Golub, Byron M Yu, and Steven M Chase. “Internal Models for Interpreting Neural Population Activity during Sensorimotor Control”. In: eLife 4 (Dec. 2015). Ed. by Timothy Behrens, e10015. issn: 2050-084X. doi: 10.7554/eLife.10015. (Visited on 01/21/2020).

[51] Sergey D. Stavisky et al. “Motor Cortical Visuomotor Feedback Activity Is Initially Isolated from Downstream Targets in Output-Null Neural State Space Dimensions”. In: Neuron 95.1 (July 2017), 195–208.e9. issn: 0896-6273. doi: 10.1016/j.neuron.2017.05.023. (Visited on 04/28/2020).

[52] Maryam M. Shanechi et al. “Rapid Control and Feedback Rates Enhance Neuroprosthetic Control”. In: Nature Communications 8.1 (Jan. 2017), p. 13825. issn: 2041-1723. doi: 10.1038/ncomms13825. (Visited on 10/27/2020).

[53] W. T. Thach. “Correlation of Neural Discharge with Pattern and Force of Muscular Activity, Joint Position, and Direction of Intended next Movement in Motor Cortex and Cerebellum”. In: Journal of Neurophysiology 41.3 (May 1978), pp. 654–676. issn: 0022-3077. doi: 10.1152/jn.1978.41.3.654.

[54] Eberhard E. Fetz. “Are Movement Parameters Recognizably Coded in the Activity of Single Neurons?” en. In: Behavioral and Brain Sciences 15.4 (Dec. 1992), pp. 679–690. issn: 1469-1825, 0140-525X. doi: 10.1017/S0140525×00072599.

[55] Eberhard E. Fetz. “Volitional Control of Neural Activity: Implications for Brain–Computer Interfaces”. In: The Journal of Physiology 579.3 (2007), pp. 571–579. issn: 1469-7793. doi: 10.1113/jphysiol.2006.127142. (Visited on 07/14/2020).

[56] Stephen H. Scott. “Inconvenient Truths about Neural Processing in Primary Motor Cortex”. en. In: The Journal of Physiology 586.5 (2008), pp. 1217–1224. issn: 1469-7793. doi: 10.1113/jphysiol.2007.146068.

[57] Mohsen Omrani et al. “Perspectives on Classical Controversies about the Motor Cortex”. In: Journal of Neurophysiology 118.3 (June 2017), pp. 1828–1848. issn: 0022-3077. doi: 10.1152/jn.00795.2016.

[58] Juan A. Gallego et al. “Neural Manifolds for the Control of Movement”. en. In: Neuron 94.5 (June 2017), pp. 978–984. issn: 0896-6273. doi: 10.1016/j.neuron.2017.05.025.

[59] Francis R. Willett et al. “Hand Knob Area of Premotor Cortex Represents the Whole Body in a Compositional Way”. en. In: Cell 181.2 (Apr. 2020), 396–409.e26. issn: 0092-8674. doi: 10.1016/j.cell.2020.02.043.

[60] Peiran Gao et al. “A Theory of Multineuronal Dimensionality, Dynamics and Measurement”. en. In: bioRxiv (Nov. 2017), p. 214262. doi: 10.1101/214262.

[61] Xiao Zhou et al. “Distinct Types of Neural Reorganization during Long-Term Learning”. In: Journal of Neurophysiology 121.4 (Feb. 2019), pp. 1329–1341. issn: 0022-3077. doi: 10.1152/jn.00466.2018. (Visited on 03/13/2020).

[62] Marvin Minsky. “Steps toward Artificial Intelligence”. In: Proceedings of the IRE 49.1 (Jan. 1961), pp. 8–30. issn: 2162-6634. doi: 10.1109/JRPROC.1961.287775.

[63] Eberhard E. Fetz. “Operant Conditioning of Cortical Unit Activity”. In: Science 163.3870 (Feb. 1969), pp. 955–958. issn: 0036-8075, 1095-9203. doi: 10.1126/science.163.3870.955. (Visited on 08/07/2020).

[64] Aaron C. Koralek et al. “Corticostriatal Plasticity Is Necessary for Learning Intentional Neuroprosthetic Skills”. In: Nature 483.7389 (Mar. 2012), pp. 331–335. issn: 1476-4687. doi: 10.1038/nature10845. (Visited on 07/22/2020).

[65] Kelly B. Clancy et al. “Volitional Modulation of Optically Recorded Calcium Signals during Neuroprosthetic Learning”. In: Nature Neuroscience 17.6 (June 2014), pp. 807–809. issn: 1546-1726. doi: 10.1038/nn.3712. (Visited on 11/25/2019).

[66] Vivek R. Athalye et al. “Evidence for a Neural Law of Effect”. In: Science 359.6379 (2018), pp. 1024–1029.

[67] Robert Legenstein, Dejan Pecevski, and Wolfgang Maass. “A Learning Theory for Reward-Modulated Spike-Timing-Dependent Plasticity with Application to Biofeedback”. In: PLOS Computational Biology 4.10 (Oct. 2008), e1000180. issn: 1553-7358. doi: 10.1371/journal.pcbi.1000180. (Visited on 08/07/2020).

[68] Peter C. Humphreys et al. BCI Learning Phenomena Can Be Explained by Gradient-Based Optimization. Dec. 2022. doi: 10.1101/2022.12.08.519453. (Visited on 05/02/2023).

[69] Jordan A. Taylor, John W. Krakauer, and Richard B. Ivry. “Explicit and Implicit Contributions to Learning in a Sensorimotor Adaptation Task”. In: J. Neurosci. 34.8 (Feb. 2014), pp. 3023–3032. issn: 0270-6474, 1529-2401. doi: 10.1523/JNEUROSCI.3619-13.2014. (Visited on 07/21/2020).

[70] Aaron L. Wong et al. “Explicit Knowledge Enhances Motor Vigor and Performance: Motivation versus Practice in Sequence Tasks”. In: Journal of Neurophysiology 114.1 (July 2015), pp. 219– 232. issn: 0022-3077. doi: 10.1152/jn.00218.2015. (Visited on 01/26/2024).

[71] Alexandre Payeur, Amy L. Orsborn, and Guillaume Lajoie. Neural Manifolds and Gradient-Based Adaptation in Neural-Interface Tasks. Mar. 2023. doi: 10.1101/2023.03.11.532146. (Visited on 05/02/2023).

[72] Jeffrey A. Kleim et al. “Cortical Synaptogenesis and Motor Map Reorganization Occur during Late, But Not Early, Phase of Motor Skill Learning”. In: J. Neurosci. 24.3 (Jan. 2004), pp. 628–633. issn: 0270-6474, 1529-2401. doi: 10.1523/JNEUROSCI.3440-03.2004. (Visited on 03/13/2020).

[73] Xulu Sun et al. “Cortical Preparatory Activity Indexes Learned Motor Memories”. en. In: Nature 602.7896 (Feb. 2022), pp. 274–279. issn: 1476-4687. doi: 10.1038/s41586-021-04329-x.

[74] Darby M. Losey et al. “Learning leaves a memory trace in motor cortex”. In: Current Biology (2024). doi: 10.1016/j.cub.2024.03.003.

[75] James B. Heald, Máté Lengyel and Daniel M. Wolpert. “Contextual Inference Underlies the Learning of Sensorimotor Repertoires”. In: Nature 600.7889 (Dec. 2021), pp. 489–493. issn: 1476-4687. doi: 10.1038/s41586-021-04129-3. (Visited on 05/23/2023).

[76] Ta-Chu Kao and Guillaume Hennequin. “Neuroscience out of Control: Control-Theoretic Perspectives on Neural Circuit Dynamics”. en. In: Current Opinion in Neurobiology. Computational Neuroscience 58 (Oct. 2019), pp. 122–129. issn: 0959-4388. doi: 10.1016/j.conb.2019.09.001.

[77] Peiran Gao and Surya Ganguli. “On Simplicity and Complexity in the Brave New World of Large-Scale Neuroscience”. en. In: Current Opinion in Neurobiology. Large-Scale Recording Technology (32) 32 (June 2015), pp. 148–155. issn: 0959-4388. doi: 10.1016/j.conb.2015.04.003.

[78] Krishna V. Shenoy and Jose M. Carmena. “Combining Decoder Design and Neural Adaptation in Brain-Machine Interfaces”. In: Neuron 84.4 (Nov. 2014), pp. 665–680. issn: 0896-6273. doi: 10.1016/j.neuron.2014.08.038. (Visited on 10/07/2020).

[79] Serafeim Perdikis and Jose del R. Millan. “Brain-Machine Interfaces: A Tale of Two Learners”. In: IEEE Systems, Man, and Cybernetics Magazine 6.3 (July 2020), pp. 12–19. issn: 2333-942X. doi: 10.1109/MSMC.2019.2958200.

[80] Jane X. Wang et al. “Prefrontal Cortex as a Meta-Reinforcement Learning System”. In: Nature Neuroscience 21.6 (June 2018), pp. 860–868. issn: 1546-1726. doi: 10.1038/s41593-018-0147-8. (Visited on 07/14/2020).

[81] Ryan M. Neely et al. “Volitional Modulation of Primary Visual Cortex Activity Requires the Basal Ganglia”. In: Neuron 97.6 (Mar. 2018), 1356–1368.e4. issn: 0896-6273. doi: 10.1016/j.neuron.2018.01.051. (Visited on 11/25/2019).

[82] N. Vendrell-Llopis et al. “Ventral Striatum Uses a Temporal Difference Rule for Prediction during Neuroprosthetic Control”. In: 2019 9th International IEEE/EMBS Conference on Neural Engineering (NER). Mar. 2019, pp. 562–565. doi: 10.1109/NER.2019.8716982.

[83] Vivek R Athalye, Jose M Carmena, and Rui M Costa. “Neural Reinforcement: Re-Entering and Refining Neural Dynamics Leading to Desirable Outcomes”. In: Current Opinion in Neurobiology. Neurobiology of Behavior 60 (Feb. 2020), pp. 145–154. issn: 0959-4388. doi: 10.1016/j.conb.2019.11.023. (Visited on 02/26/2020).

[84] Barbara Feulner et al. “Small, Correlated Changes in Synaptic Connectivity May Facilitate Rapid Motor Learning”. In: Nat Commun 13.1 (Sept. 2022), p. 5163. issn: 2041-1723. doi: 10.1038/s41467-022-32646-w. (Visited on 05/02/2023).

[85] Ricky T. Q. Chen et al. “Neural Ordinary Differential Equations”. In: Advances in Neural Information Processing Systems 31. Curran Associates, Inc., 2018, pp. 6571–6583. (Visited on 08/02/2020).

[86] Richard H. Byrd et al. “A Limited Memory Algorithm for Bound Constrained Optimization”. In: SIAM Journal on Scientific Computing 16.5 (Sept. 1995), pp. 1190–1208. issn: 1064-8275. doi: 10.1137/0916069.

[87] A. P. Georgopoulos, A. B. Schwartz, and R. E. Kettner. “Neuronal Population Coding of Movement Direction”. en. In: Science 233.4771 (Sept. 1986), pp. 1416–1419. issn: 0036-8075, 1095-9203. doi: 10.1126/science.3749885.

## References

[1] Guillaume Hennequin, Tim P. Vogels, and Wulfram Gerstner. “Non-Normal Amplification in Random Balanced Neuronal Networks”. In: Phys. Rev. E 86.1 (July 2012), p. 011909. doi: 10.1103/PhysRevE.86.011909. (Visited on 04/15/2020).

[2] Guillaume Hennequin, Tim P. Vogels, and Wulfram Gerstner. “Optimal Control of Transient Dynamics in Balanced Networks Supports Generation of Complex Movements”. en. In: Neuron 82.6 (June 2014), pp. 1394–1406. issn: 0896-6273. doi: 10.1016/j.neuron.2014.04.045.

[3] David Sussillo et al. “A Neural Network That Finds a Naturalistic Solution for the Production of Muscle Activity”. In: Nature Neuroscience 18.7 (July 2015), pp. 1025–1033. issn: 1546-1726. doi: 10.1038/nn.4042. (Visited on 04/22/2020).

[4] E. Todorov and W. Li. “Optimal Control Methods Suitable for Biomechanical Systems”. In: Proceedings of the 25th Annual International Conference of the IEEE Engineering in Medicine and Biology Society (IEEE Cat. No.03CH37439). Vol. 2. Sept. 2003, 1758–1761 Vol.2. doi: 10.1109/IEMBS.2003.1279748.

[5] Ta-Chu Kao, Mahdieh S. Sadabadi, and Guillaume Hennequin. “Optimal Anticipatory Control as a Theory of Motor Preparation: A Thalamo-Cortical Circuit Model”. en. In: Neuron 109.9 (May 2021), 1567–1581.e12. issn: 0896-6273. doi: 10.1016/j.neuron.2021.03.009.

[6] Diederik P. Kingma and Jimmy Ba. “Adam: A Method for Stochastic Optimization”. In: arXiv:1412.6980 [cs] (Jan. 2017). 1412.6980 [cs]. (Visited on 03/13/2021).

[7] Friedrich Schuessler et al. “The Interplay between Randomness and Structure during Learning in RNNs”. In: Advances in Neural Information Processing Systems. Vol. 33. Curran Associates, Inc., 2020, pp. 13352–13362. (Visited on 01/26/2024).

[8] Francesca Mastrogiuseppe and Srdjan Ostojic. “Linking Connectivity, Dynamics, and Computations in Low-Rank Recurrent Neural Networks”. In: Neuron 99.3 (Aug. 2018), 609–623.e29. issn: 0896-6273. doi: 10.1016/j.neuron.2018.07.003. (Visited on 01/24/2020).

[9] Laureline Logiaco, L. F. Abbott, and Sean Escola. “Thalamic Control of Cortical Dynamics in a Model of Flexible Motor Sequencing”. en. In: Cell Reports 35.9 (June 2021), p. 109090. issn: 2211-1247. doi: 10.1016/j.celrep.2021.109090.

[10] Beata Jarosiewicz et al. “Functional Network Reorganization during Learning in a Brain-Computer Interface Paradigm”. In: Proceedings of the National Academy of Sciences 105.49 (Dec. 2008), pp. 19486–19491. issn: 0027-8424, 1091-6490. doi: 10.1073/pnas.0808113105. (Visited on 02/04/2020).

[11] Karunesh Ganguly et al. “Reversible Large-Scale Modification of Cortical Networks during Neuroprosthetic Control”. In: Nature Neuroscience 14.5 (May 2011), pp. 662–667. issn: 1546-1726. doi: 10.1038/nn.2797. (Visited on 04/13/2020).

[12] Huibert Kwakernaak and Raphael Sivan. Linear Optimal Control Systems. Vol. 1. Wiley-interscience New York, 1972.

[13] Emanuel Todorov and Michael I. Jordan. “Optimal Feedback Control as a Theory of Motor Coordination”. en. In: Nature Neuroscience 5.11 (Nov. 2002), pp. 1226–1235. issn: 1546-1726. doi: 10.1038/nn963.

[14] Matthew D Golub, Byron M Yu, and Steven M Chase. “Internal Models for Interpreting Neural Population Activity during Sensorimotor Control”. In: eLife 4 (Dec. 2015). Ed. by Timothy Behrens, e10015. issn: 2050-084X. doi: 10.7554/eLife.10015. (Visited on 01/21/2020).

[15] Sergey D. Stavisky et al. “Motor Cortical Visuomotor Feedback Activity Is Initially Isolated from Downstream Targets in Output-Null Neural State Space Dimensions”. In: Neuron 95.1 (July 2017), 195–208.e9. issn: 0896-6273. doi: 10.1016/j.neuron.2017.05.023. (Visited on 04/28/2020).

[16] Maryam M. Shanechi et al. “Rapid Control and Feedback Rates Enhance Neuroprosthetic Control”. In: Nature Communications 8.1 (Jan. 2017), p. 13825. issn: 2041-1723. doi: 10.1038/ncomms13825. (Visited on 10/27/2020).

[17] Michael E. Tipping and Christopher M. Bishop. “Probabilistic Principal Component Analysis”. In: Journal of the Royal Statistical Society: Series B (Statistical Methodology) 61.3 (1999), pp. 611–622. issn: 1467-9868. doi: 10.1111/1467-9868.00196. (Visited on 11/12/2020).

[18] Patrick T. Sadtler et al. “Neural Constraints on Learning”. In: Nature 512.7515 (Aug. 2014), pp. 423–426. issn: 1476-4687. doi: 10.1038/nature13665. (Visited on 02/13/2020).

[19] Matthew D. Golub et al. “Learning by Neural Reassociation”. In: Nature Neuroscience 21.4 (Apr. 2018), pp. 607–616. issn: 1546-1726. doi: 10.1038/s41593-018-0095-3. (Visited on 11/11/2019).

[20] Ake Björck and Gene H. Golub. “Numerical Methods for Computing Angles between Linear Subspaces”. en. In: Mathematics of Computation 27.123 (1973), pp. 579–594. issn: 0025-5718, 1088-6842. doi: 10.1090/S0025-5718-1973-0348991-3.

